# Mitochondrial genomes of two parasitic *Cuscuta* species lack clear evidence of horizontal gene transfer and retain unusually fragmented *ccmF_C_* genes

**DOI:** 10.1101/2021.04.15.439983

**Authors:** Benjamin M Anderson, Kirsten Krause, Gitte Petersen

## Abstract

**Background:** The intimate association between parasitic plants and their hosts favours the exchange of genetic material, potentially leading to horizontal gene transfer (HGT) between plants. With the recent publication of several parasitic plant nuclear genomes, there has been considerable focus on such non-sexual exchange of genes. To enhance the picture on HGT events in a widely distributed parasitic genus, *Cuscuta* (dodders), we assembled and analyzed the organellar genomes of two recently sequenced species, *C. australis* and *C. campestris*, making this the first account of complete mitochondrial genomes (mitogenomes) for this genus.

**Results:** The mitogenomes are 265,696 and 275,898 bp in length and contain a typical set of mitochondrial genes, with ten missing or pseudogenized genes often lost from angiosperm mitogenomes. Each mitogenome also possesses a structurally unusual *ccmF_C_* gene, which exhibits splitting of one exon and a shift to trans-splicing of its intron. Based on phylogenetic analysis of mitochondrial genes from across angiosperms and similarity-based searches, there is little to no indication of HGT into the *Cuscuta* mitogenomes. A few candidate regions for plastome-to-mitogenome transfer were identified, with one suggestive of possible HGT.

**Conclusions:** The lack of HGT is surprising given examples from the nuclear genomes, and may be due in part to the relatively small size of the *Cuscuta* mitogenomes, limiting the capacity to integrate foreign sequences.

## Background

Horizontal gene transfer (HGT) may be broadly defined as the movement of genetic material between species by a means other than normal reproduction (e.g. fusion of gametes). The process has been inferred within and between both prokaryotes [1] and eukaryotes [2], and is now thought to potentially have a role in adaptation, as shown in recent examples from plants [3, 4]. While the concept is sometimes extended to include transfers between cellular compartments (e.g. chloroplast to mitochondrion), we here refer to HGT as exchange of genetic material between organisms and consider it distinct from intracellular transfers.

Mechanisms that facilitate HGT between plants have been postulated to include direct contact via e.g. wounds/grafting [5], parasitic haustoria [6], epiphytic interactions [7] and illegitimate pollination [8]. Indirect routes have also been suggested, such as via soil, bacteria, viruses or endophytic fungi [8]. Early cases of detected HGT in plants [9, 10] were principally found in their mitochondrial genomes (mitogenomes), possibly partly as a result of the greater sequence availability at the time compared to the nuclear genome. Cellular mechanisms such as active DNA uptake [11] and organelle fusion and fission [12, 13] have been referenced to explain why HGT is more prevalent in the mitogenome, at least compared with the extensively sequenced chloroplast genome (plastome). While examples of HGT from nuclear genomes are accumulating, mitogenome examples still remain a substantial portion of the evidence for HGT in plants.

A group of plants whose lifestyle makes them more likely to experience HGT are parasites [14, 15], largely because of their close physical interaction with their hosts via specialised connective organs called haustoria. Parasitic plants rely on other plants for water and often for a source of carbon, having some degree of photosynthetic ability (hemiparasites) or having lost it (holoparasites). The altered need for photosynthetic capacity has impacted the size and gene content of plastomes of many parasitic plants [16, 17]. Parasites in the family Viscaceae also have markedly divergent and reduced gene content in their mitogenomes [18–20], but such a reduction was not observed in other groups of parasites [20, 21]. The more common feature associated with parasitic plant mitochondria is the presence of foreign DNA. An extreme example of incorporated foreign DNA in a mitogenome is *Lophophytum mirabile* (Balanophoraceae), which is inferred to have acquired approximately 80% of its genes and 60% of its sequence from its host [22, 23]. Another example includes members of Rafflesiaceae, for which up to 40% of their mitochondrial genes are inferred to have been transferred, with many of the putatively transferred genes shown to be actively transcribed [24], though whether the inferred transfers are indeed foreign and/or functional has recently been questioned [25].

Parasitic plants in the genus *Cuscuta* are twining stem parasites with reduced/absent leaves that rely on their host plants for both water and a carbon source obtained via haustorial connections [26]. Photosynthetic ability is variably retained in the genus, and there is a diversity of chloroplast structure and plastome gene content [27–31]. For example, while some species are known to photosynthesize to some degree, carbon fixation in the bright yellow species *C. campestris* is below the compensation point [27] making it unable to survive and grow, let alone complete its life cycle, without its hosts. The range of possible host species for *Cuscuta* varies from narrow to especially broad with examples including members of Fabaceae, Apiaceae, Amaranthaceae, Asteraceae, Solanaceae, Linaceae, Ericaceae, Rutaceae and Amaryllidaceae among others [26]. The haustoria that *Cuscuta* plants use to obtain nutrients from their hosts are capable of both phloem and xylem connections [32] and are also capable of transmitting genetic material in the form of RNA [6] and DNA [4, 33], which together with the phylogenetically broad host range makes *Cuscuta* a potentially ideal study system for exploring the phenomenon of HGT.

A number of studies have implicated *Cuscuta* species as sources and recipients of HGT events. Studies of host *Plantago* [34, 35] and *Geranium* [36] species show evidence of multiple mitochondrial genes likely obtained from *Cuscuta* parasites. Multiple transcriptomic surveys identified candidate nuclear genes in *Cuscuta* that may have been acquired by HGT [37–39]. Nuclear genome assemblies coupled with transcriptomic work have also revealed numerous candidates for HGT in *Cuscuta campestris* [4, 33]. A recent report of an *Agrobacterium* gene in three *Cuscuta* species could also represent an HGT from another plant, though it could have also been acquired directly from the bacterium [40]. To date the evidence suggests there is active exchange of genetic information between *Cuscuta* and its hosts, with donations of mitochondrial genes and acquisitions of multiple nuclear genes, but not yet any indication of imported mitochondrial genetic material.

While nuclear genomes for *Cuscuta australis* [41] and *C. campestris* [33] have been assembled, our knowledge of the mitogenome in *Cuscuta* is limited to a subset of genes from *Cuscuta gronovii* [36]. Given the lack of knowledge about *Cuscuta* mitogenomes and the expectation that the mitogenome is a prime location to look for HGT, we set out to assemble and characterise mitogenomes for *Cuscuta australis* and *C. campestris*. The complete genomes will enable us to assess the evidence for HGT, and secondarily intracellular transfers between genomic compartments, in *Cuscuta* mitogenomes, and test the unspoken assumption that examples of HGT in *Cuscuta* should be abundant.

## Methods

We downloaded from GenBank portions of previously published datasets from whole genome sequencing studies of *Cuscuta australis* and *C. campestris* [33, 41]. For *C. australis*, we downloaded 150 bp paired-end reads from an Illumina 350 bp insert library (SRA accession SRR5851367; 155 M reads) and PacBio long reads (SRA accessions SRR5851361–6; 711 k reads). For *C. campestris*, we downloaded 301 bp paired-end reads from Illumina 800 bp insert libraries (SRA accessions ERR1917169–71; 9.5 M reads) and PacBio long reads (SRA accessions ERR1910865–7; 490 k reads). Illumina short reads were evaluated with FastQC [42] before and after read deduplication, adapter and low quality base trimming, error correction and read merger using Clumpify, BBDuk, BBMerge and Tadpole in the BBMap package v. 38.08 [43; https://sourceforge.net/projects/bbmap/]. As part of initial assembly, long reads were corrected then trimmed using default settings in Canu v. 1.8 [44]. After initial quality control, the datasets consisted of 1 M merged and 122 M unmerged short reads and 399 k long reads for *C. australis*, and 316 k merged and 7 M unmerged short reads and 79 k long reads for *C. campestris*.

For transcriptome data of *C. australis* [41], we downloaded 150 bp paired-end reads for various tissues (SRR6664647–54; 411 M reads). After read deduplication and adapter and low quality base trimming using Clumpify and BBDuk, 268 M reads were retained with an average read length of 142 bp (72% of reads were untrimmed). Transcriptomic reads for *Ipomoea nil* were obtained from three sources: the 1KP project [45], ERR2040616; a transcriptome study [46], SRR1772255; and a genome project [47], DRR024544–9. Reads were also deduplicated and trimmed for adapters and low quality bases, with 157 M reads retained with an average read length of 97 bp (81% of reads were untrimmed).

### Organelle genome assemblies

We approached organellar assembly by running separate short and long read *de novo* assemblies, searching resulting contigs using BLASTN v. 2.6.0+ [48–50] with default parameters for hits to reference genomes (see Tables S1 and S2, Additional file 1), then using those contigs with hits as targets for further mapping with BBMap and assembly.

Short read datasets were first filtered using the Kmer Analysis Toolkit (KAT) v. 2.4.2 [51] to select a high coverage subset of reads for easier assembly. Coverage targets were based on mapping reads to sequences from reference plastome (*Cuscuta exaltata*, NC_009963) and mitogenome (*Ipomoea nil*, NC_031158) CDS. Based on the mapping, we chose kmer depth filters of 500–25000 selecting ~35 M reads and 30–2100 selecting ~700 k reads for *C. australis* and *C. campestris*, respectively. The filtered read sets were assembled using Unicycler v. 0.4.8 [52], which when given only short reads optimizes a SPAdes v. 3.14.0 [53] assembly.

Long read datasets were assembled in Canu v. 1.8 [44] using default settings and with genome size estimates of 280 M and 550 M for *C. australis* and *C. campestris*, respectively. In order to complete one run, we set the stopOnLowCoverage parameter to 0.1 for *C. campestris*.

We examined contigs from the short and long read assemblies in Bandage v. 0.8.1 [54] to identify those with BLAST (blastn megablast default parameters) hits to the reference genomes and which had similar depths (for distinguishing plastid and mitochondrial contigs). We considered as candidates contigs with > 50% coverage of hits to the respective references, and in cases of less than 50% coverage, whether read depth was similar to depth of contigs having greater coverage of hits. Contigs with hits and identified as putatively plastid or mitochondrial were used as targets for mapping the full sets of short reads to create input datasets for Unicycler. The mapped reads were assembled along with long reads (corrected and trimmed with Canu) in Unicycler, which assembles long reads with miniasm [55] and Racon v. 1.4.3 [56] and uses the assembly to resolve repeats and create bridges in the SPAdes small read assembly graph.

Plastome assemblies were obtained after a single round of Unicycler, producing single circular contigs. In order to complement the assembly approach, we also used NOVOPlasty v. 3.7.2 [57] with the deduplicated and adapter-trimmed (but not error corrected or merged) short read datasets. We used an *rbcL* sequence from *Cuscuta exaltata* as a seed, which NOVOPlasty uses to select an initial matching read and begin iterative extension until it can circularize an assembly. The resulting two options per assembly (different configurations across the inverted repeats) were then compared with BLAST to the assemblies from Unicycler to verify the sequences were identical.

The mitogenome assemblies were more challenging, given the presence of long repeats and linear and circular contigs that in some cases could be joined at multiple locations, but this differed between assemblies. For *C. australis*, a second round of Unicycler was run after extending contigs by up to 100 kbp using Tadpole (part of the BBMap package) with settings el=100000 er=100000 mode=extend k=31 and the full short read dataset, and then mapping short reads again. Simplifying the assembly graph included removing small contigs that were fully present in a larger contig or were too short (e.g. <100 bp). We then checked long read contigs from the initial assembly in Bandage to ensure they were covered by hits from the remaining assembly. Since there were four linear and four circular contigs, and since the circular contigs did not share repeats that could be used to merge them, we were unable to merge all contigs into a single contig. Repeats are present (though there are no long repeats shared between circles) and there are alternative paths through the assembly, so our representation is one of multiple possible and does not reflect the actual structure of the mitochondrial genome in the plants. While we were not able to merge the contigs in *C. australis*, we could join circular contigs for the assembly of *C. campestris*. For *C. campestris*, the first round of Unicycler produced four circular contigs along with three small linear contigs (linear totalling <5 kb) which hit the plastome or had low depth and were absent from the long read contigs. The short linear contigs were not included in further assembly. The four circles had shared long repeats, and these were examined using long reads to determine if multiple configurations were present, suggesting the separate contigs are sometimes contiguous. Long reads were mapped and aligned to extracted repeats with Minimap2 v. 2.14-r883 [55], Samtools v. 1.7 [58] and MAFFT v. 7.310 [59] and using a custom Python script including ‘pysam’ (https://github.com/pysam-developers/pysam). When the reads indicated multiple configurations across the repeat, we manually joined the circles. The circles all shared repeats that showed multiple configurations, so we were able to merge the four circles into a single circular contig.

### Read mapping and annotation

In order to check the mitogenome assemblies for clear errors, we conservatively mapped short and long reads then manually compared the resulting BAM files in IGV v. 2.6+ [60]. Short reads were mapped using BBMap with settings pairedonly=t minid=0.9 ambiguous=random maxindel=100 strictmaxindel=t pairlen=1000. Long reads were mapped using Minimap2 in mode ‘map-pb’ and filtered with Samtools with option -F 2308, followed by filtering the resulting BAM file with a custom Python script using ‘pysam’ to only retain mappings where reads covered the target for at least 80% of their length. Long reads were used to check structural assembly and cases of insertions, where high coverage plastid reads can lead to misassembly of the mitogenomic sequence. The long reads passing through an insertion can be used to identify the lower-coverage short reads that are mitochondrial and allow for manual correction. For *C. australis*, only one location suggested a small (28 bp) sequence that needed to be deleted (absent in most long reads). For *C. campestris*, one location of a plastome insert needed three single base pair corrections and a small (9 bp) deletion to match the long reads that spanned the region rather than the higher depth plastid reads.

Plastomes and mitogenomes were annotated using custom scripts to BLAST (BLASTN, BLASTX, default search settings) against nucleotide and protein databases built using reference plastid (14 from 13 genera and 13 families, see Table S1, Additional file 1) and mitochondrial (172 from 130 genera and 62 families, see Table S2, Additional file 1) genomes, an approach similar to that used in Mitofy [61]. For mitogenomes, we also used the references to build protein alignments for constructing a HMMER v. 3.3 [62; http://hmmer.org] database that we used to search open reading frames (ORFs) found using ORFfinder v. 0.4.3 (NCBI; https://www.ncbi.nlm.nih.gov/orffinder/). Following searches, hits were examined and adjusted manually, with input from MFAnnot v. 1.35 (https://github.com/BFL-lab/Mfannot) to select intron boundaries. Transfer RNAs were also detected and checked with tRNAscan-SE 2.0 [63]. Pseudogenes were subjectively annotated if hits had clear frameshifts and/or internal stop codons, and gene copies broken by repeats or alternative paths in the assembly were annotated as fragments.

Dispersed repeats were detected by searching the organellar genomes against themselves using BLASTN with an E value of 1e-10 and filtering out hits less than 50 bp and lower than 90% identity. Overlapping hits were then filtered with a custom python script, and the percentage repeat content calculated. For visualization, the genomes were searched against themselves using Circoletto [64] with an E value chosen to approximate a 90% identity threshold (1e-70 and 1e-40 for *C. australis* and *C. campestris*, respectively). As a proxy for recombinational activity, we used long repeats (>500 bp, >90% identity) and mapped long reads. A random selection of up to 50 long reads that mapped to each repeat were aligned with the repeat and evaluated manually for whether they showed indications of incongruence on either side of the repeat. The proportion of the mapped reads showing incongruence was used as a proxy for recombinational activity. Tandem repeat content was determined using Tandem Repeat Finder v. 4.09 [65] with recommended settings: 2 7 7 80 10 50 500. Organelle genome figures were based on output from OGDRAW [66].

To investigate the transcription and splicing of the mitochondrial *ccmF_C_* gene, we mapped mRNA reads using BBMap with the setting pairedonly=t against three DNA sequences manually stitched together and separated by N’s from the annotated *ccmF_C_* locations in the *C. australis* assembly. We also aligned *ccmF_C_* DNA sequences from both *Cuscuta* species with *ccmF_C_* genes and CDS from reference mitogenomes using MAFFT to determine the locations of the exon and intron boundaries. We visualized alignments and mapping in Jalview v. 2.11.0 [67] and IGV v. 2.8.0 [60]. Intron domains were roughly determined by iteratively running the Mfold web server [68] using the ccmFci829 from *Nicotiana tabacum* (NC_006581). RNA editing sites were predicted with PREP-MT [69] to compare with those observed in the transcriptomic reads. We also aligned *Ipomoea nil* transcriptomic sequences to similar *ccmF_C_* DNA sequences from its mitogenome to compare with *Cuscuta*.

To evaluate the notable gene structure of the final portion of *nad1*, we aligned sequences from *Ipomoea nil, C. australis* and *C. campestris* using Mauve v. snapshot_2015-02-13 [70]. We also mapped transcriptomic reads from *C. australis* to *nad1* exons separated by 100 Ns to evaluate splicing and whether there was evidence for an intact product.

### Potential intracellular transfers

To detect the presence of transfers from a plastome to the mitogenome, we searched each mitogenome assembly against the reference plastomes (Table S1, Additional file 1) using BLASTN with default parameters. We also searched the mitochondrial ORFs against a HMMER database built with the reference plastomes to detect whether there were intact portions of plastid genes present and to interpret blast hits. Broader searches used the NCBI non-redundant nucleotide database (https://blast.ncbi.nlm.nih.gov/). We evaluated plastome hits of interest phylogenetically using top NCBI results in RAxML v. 8.2.12 [71] with rapid bootstrapping and a search for best tree in a single run, opting for 1000 bootstrap replicates under the GTRGAMMA model.

To determine whether any missing mitochondrial genes were present in the nuclear genome, and whether there were other transfers between the mitogenome and the nuclear genome, we conducted BLAST searches of the assemblies and of the reference mitogenomes (Table S2, Additional file 1) against the two published nuclear genomes (*C. australis*: GCA_003260385.1, *C. campestris*: PRJEB19879). For assessment of total hit coverage of the assemblies, we ignored hits <250 bp and with <80% identity. For searches for missing genes, we ignored hits with <50% identity.

### Horizontal gene transfer

We investigated HGT using a phylogenetic approach, in which we built phylogenetic trees for each mitochondrial gene present in *Cuscuta* (as well as one based on all genes concatenated) and examined whether *Cuscuta* sequences were supported as grouping with their close relatives (particularly *Ipomoea nil*) or with more distantly related plants. Using reference mitogenomes (see Table S2, Additional file 1) and the *Cuscuta* assemblies along with data from *C. gronovii* (GenBank accessions KP940494–514), we extracted all CDS protein translations and corresponding nucleotide sequences, as well as nucleotide sequences for ribosomal RNAs. We removed some sequences in cases of high divergence (e.g. for *Viscum scurruloideum*) and when alignments suggested problematic annotations, and in some cases we had to reverse complement ribosomal RNA sequences. For each gene, we created protein alignments with ClustalOmega v. 1.2.4 [72] using default settings. We then converted the protein alignments back to nucleotide alignments with PAL2NAL v. 14 [73] using default settings. Alignments were run through PREP-MT [69] to predict editing sites, with sites having a prediction score of >=0.5 removed prior to analysis. Alignments were checked manually for apparent misalignments and problems with annotation around intron boundaries, and the *cox1* co-conversion tract [74] was removed. For ribosomal RNA sequences, we aligned with MAFFT v. 7.310 using options --localpair and --maxiterate 1000. For each alignment we used a custom python script to eliminate positions with >50% missing data, then ran RAxML v. 8.2.12 [71] with rapid bootstrapping and a search for best tree in a single run, opting for 1000 bootstrap replicates under the GTRGAMMA model. Finally, we plotted the resulting trees using R package ape [75], rooting them on non-angiosperm seed plants, and manually examined whether *Cuscuta* sequences grouped together, with their close relative *Ipomoea nil*, and with Solanales or whether they grouped elsewhere with bootstrap support >80%.

To explore intergenic sequences, we searched the reference mitogenomes against the *Cuscuta* assemblies using BLASTN with default parameters, removing hits <250 bp and with <80% identity. For clearer display, we used custom scripts to filter BLAST hits so that hits of equivalent length (within 50 bp) or less and more than 2% lower identity than top hits were removed along the assembly. We plotted hits using the R package Sushi v. 1.26.0 [76]. We then manually examined hits in intergenic regions, looking for where the top hits were not from Solanales and evaluating whether the hits were suggestive of longer, potentially transferred tracts.

In cases of possibly chimeric gene sequences based on top blast hits, we assessed evidence for conversion using GENECONV v. 1.81 [77].

## Results

### Organelle genomes

The plastomes were assembled as single circular molecules having typical quadripartite structure with two inverted repeats separating small and large single copy regions (see Figs. S1 and S2, Additional file 2). The plastomes are 85,263 and 86,749 bp, with GC contents of 37.8 and 37.7% for *C. australis* and *C. campestris*, respectively. Beyond the inverted repeats (14,070 and 14,348 bp in *C. australis* and *C. campestris*, respectively), there are no dispersed repeats longer than 50 bp and with identity >90%. Tandem repeats make up approximately 0.2 and 0.4% of the plastomes of *C. australis* and *C. campestris*, respectively. Similar to other reduced plastomes of *Cuscuta* species such as *C. gronovii* [29] and *C. obtusiflora* [28], the plastomes are missing all *ndh* genes (a pseudogenized copy of *ndhB* is present in *C. campestris), infA, matK, psaI*, all *rpo* genes, *rpl23, rpl32*, and *rps16*. Missing tRNAs include *trnA*-UGC, *trnG*-UCC, *trnI*-GAU, *trnK*-UUU, *trnR*-ACG, and *trnV*-UAC.

The mitogenomes were assembled as 265,696 and 275,898 bp, with GC contents of 44.0 and 44.3% for *C. australis* and *C. campestris*, respectively (Fig. 1). Dispersed repeats (see Table 1; Fig. S3, Additional file 2) comprise approximately 21.8 and 34.1% while tandem repeats make up approximately 5.0 and 2.5% of the mitogenomes of *C. australis* and *C. campestris*, respectively. Read depth was largely consistent across the assemblies, except for drops at ends of contigs and in some locations possibly representing alternative conformations (see Fig. S4, Additional file 2). Recombinational activity as assessed with long repeats varied from 10–48% (average = 29%, n=6) and 33–47% (average = 37%, n=12) for *C. australis* and *C. campestris*, respectively (see Tables S3 and S4, Additional file 2). For each mitogenome, we annotated 31 protein-coding genes, 3 ribosomal RNA genes, and 19 tRNAs (some in multiple copies; Table 1). Compared to *C. australis, C. campestris* has an extra *trnQ*-UUG of plastome origin (within a possibly transferred region), but *trnT*-UGU is missing. Common missing or pseudogenized genes for both genomes include *sdh3* and *sdh4, rpl2* and *rpl10*, and six *rps* genes (*rps2, 7, 10, 11, 14, 19*). Some extra fragments of already present genes broken up by possible alternative paths through the assembly include *atp1* and *cox1* for both species, and *atp6, rps12* and *rps3* for *C. campestris*.

**Fig. 1.**
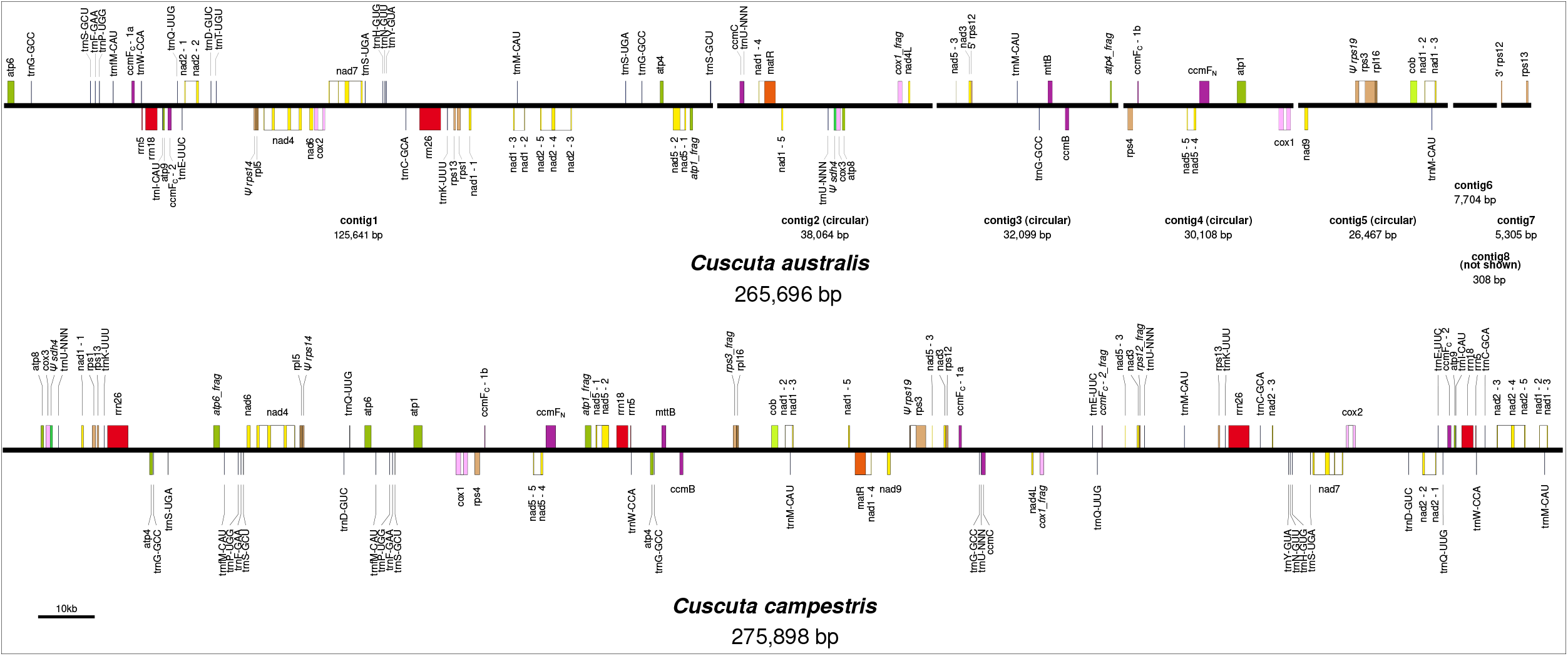
Linear representations of mitogenomes of *Cuscuta australis* and *C. campestris*. The assembly of *C. australis* consists of eight contigs (divided by gaps), while that of *C. campestris* is represented as a single circle. Pseudogenes are indicated with a capital psi (Ψ). Fragmented genes have “_frag” at the end of their labels. Transcription is to the right above the lines, and to the left below the lines.

**Table 1.**
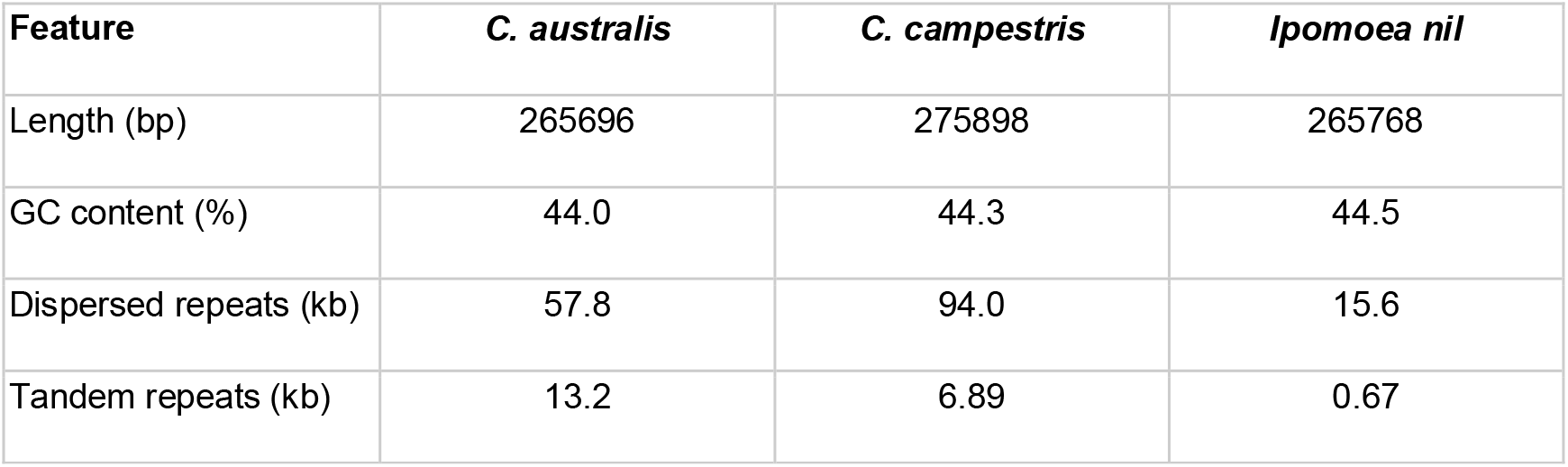
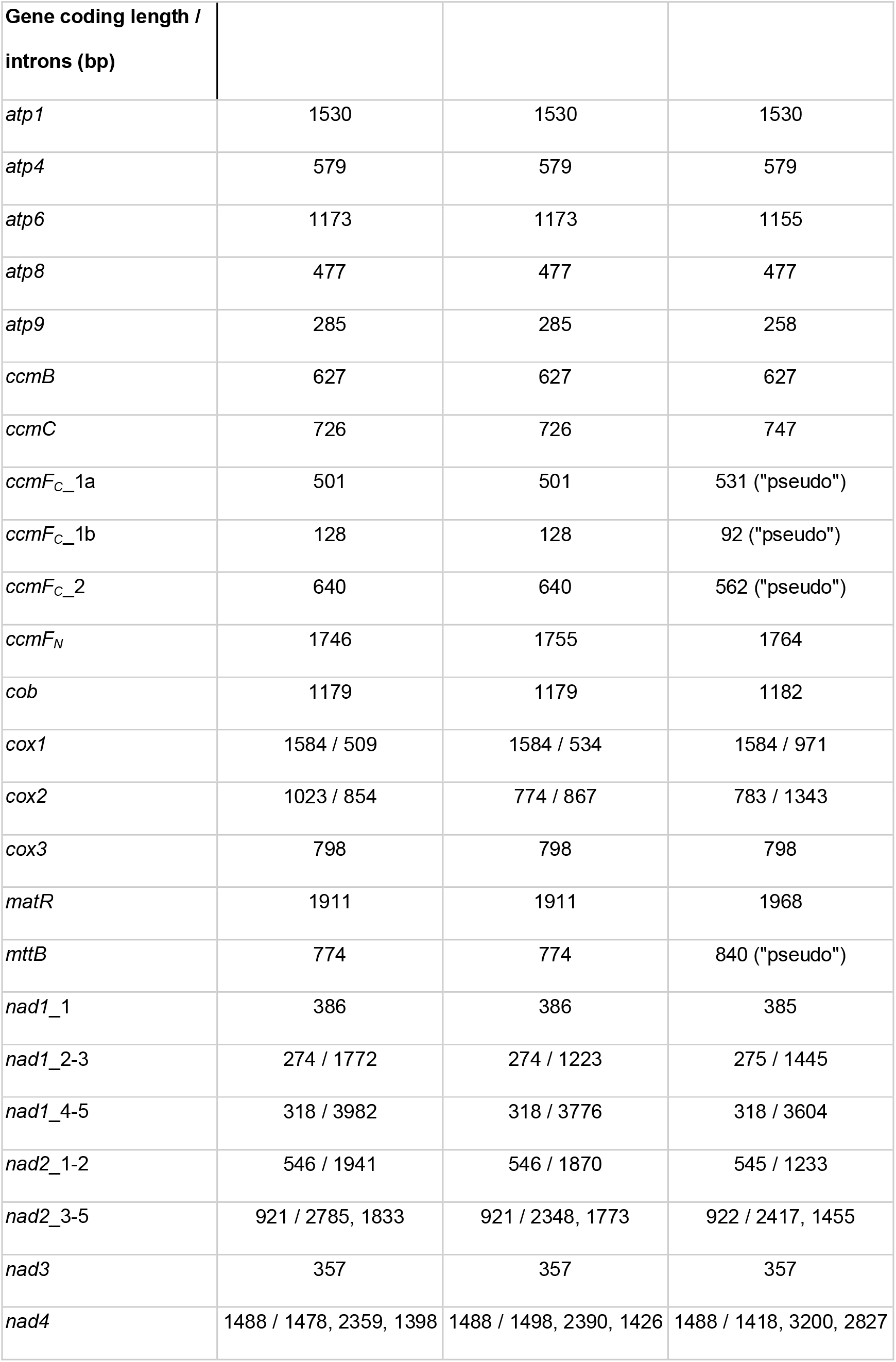
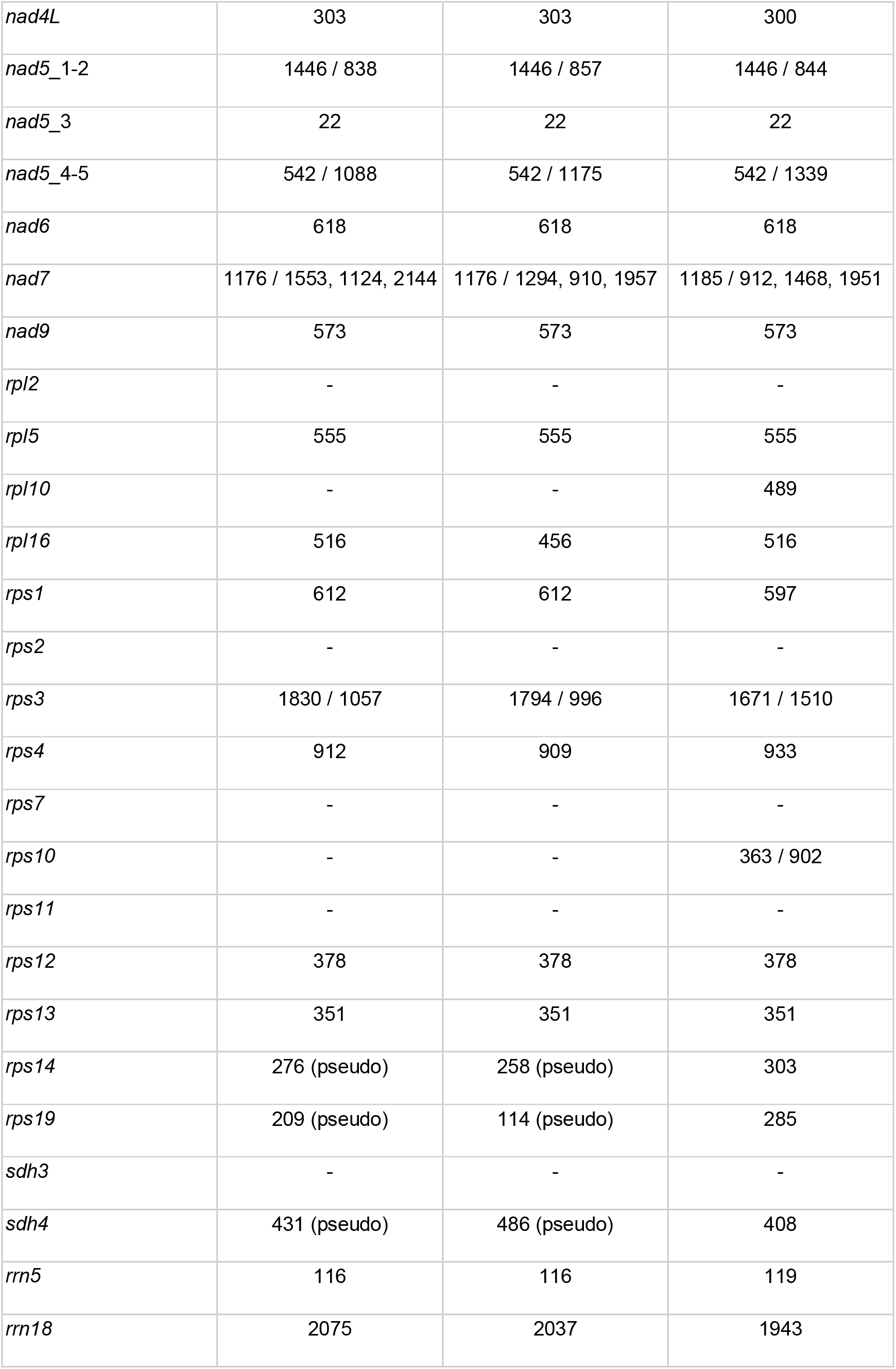
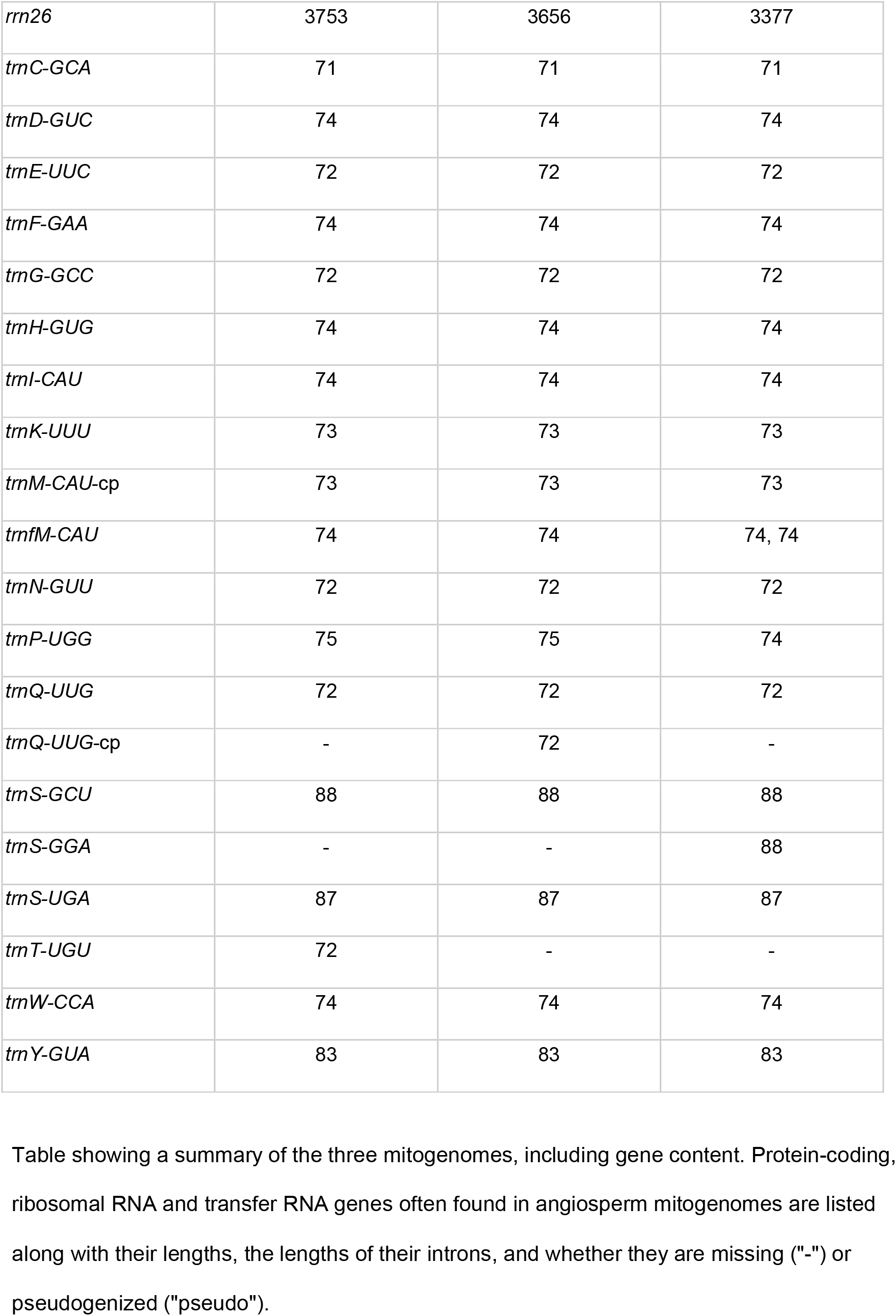
Features of the mitogenomes of *Cuscuta australis, C. campestris* and their close relative *Ipomoea nil* (NC_031158).

The *ccmF_C_* gene was problematic to annotate in both species (annotated as a pseudogene in *C. gronovii* and *Ipomoea nil*) and may not be properly transcribed, as it appears to be both trans-spliced and missing a portion of the first exon. Based on alignments with other angiosperms, including the intron-less *Platycodon grandiflorus* (NC_035958), it appears that the *ccmF_C_* gene is split across three locations in both of the *Cuscuta* mitogenomes (Fig. 2). A large portion of exon 1 is separated from the 3’ end of exon 1 and most of the group II intron. The remainder of the intron plus exon 2 are in a third location. Mapping mRNA reads resulted in low coverage of the *ccmF_C_* gene, but provided some support for successful trans-splicing of the 3’ end of exon 1 and exon 2 (see Fig. 2). There was no indication of an mRNA product that included both portions of exon 1, however, making it unclear whether the majority of exon 1 is actually translated into part of a functional product. PREP-MT predicted 14 editing sites, three of which didn’t have transcriptome support, and missed four sites that appeared to be RNA edited based on transcriptome read mapping (see Fig. S5, Additional file 2). Mapping mRNA reads against other mitochondrial gene sequences (e.g. *atp6, ccmC*; data not shown) recovered similarly low read depths as the mapping to *ccmF_C_*, suggesting read depth in this case is unreliable for effectively determining relative expression of mitochondrial genes. For accurate investigation of mitochondrial transcription of the genes of interest, other techniques like mitochondrial run-on transcription assays [78] would be more suitable.

**Fig. 2.**
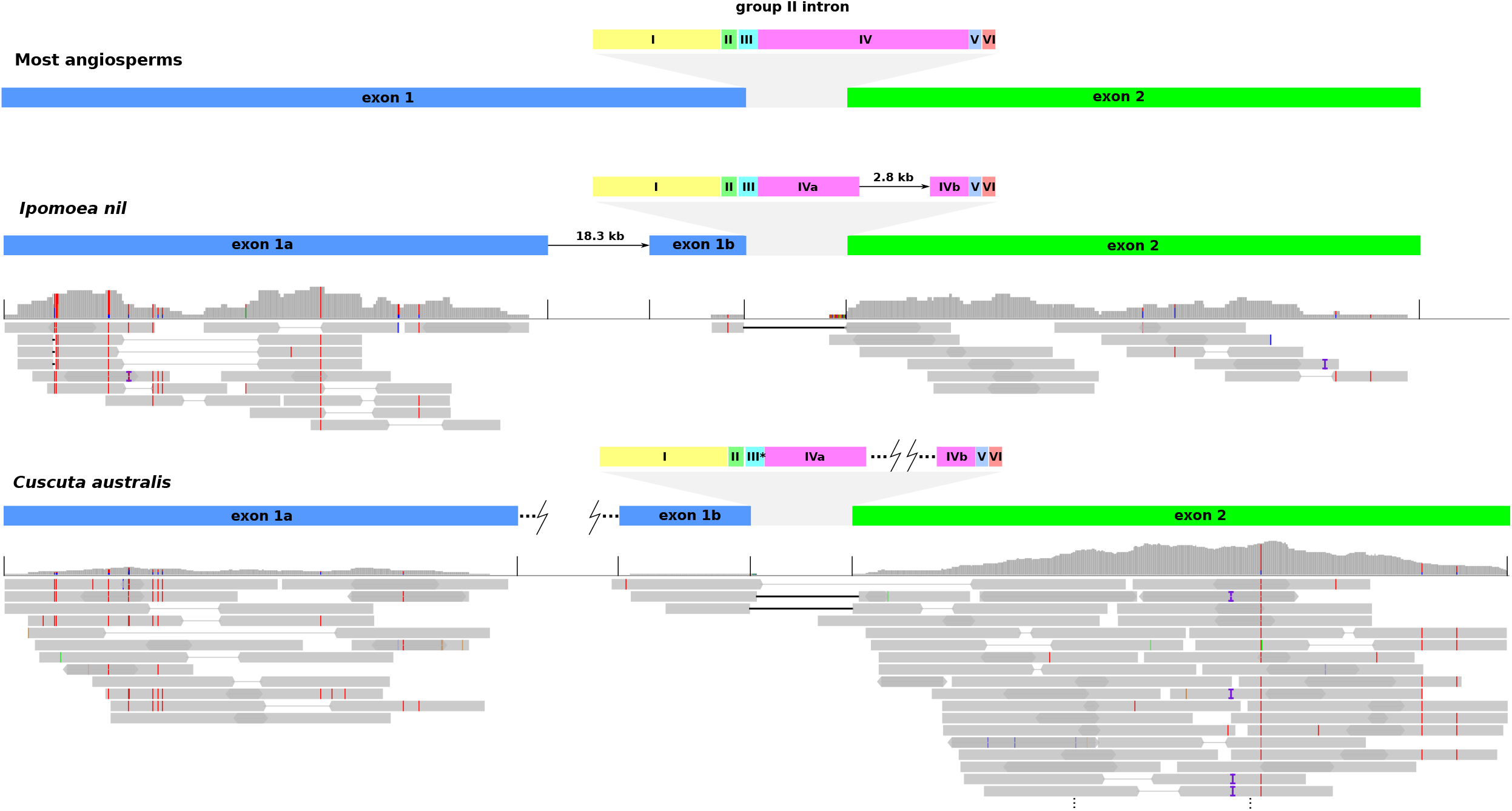
Schematic of *ccmF_C_* gene structure in *Cuscuta australis* and its close relative *Ipomoea nil* compared to typical structure in angiosperms. Transcriptomic reads from both species are mapped to three DNA sequences with high similarity to *ccmF_C_* from *Cuscuta australis* and three from *Ipomoea nil*. Red lines indicate mismatches in the mapped RNA reads and correspond to putative RNA editing sites. Light grey lines connect read pairs, while solid black lines indicate a break in a single read. Group II intron domains are denoted with Roman numerals. *The third domain in *Cuscuta* may not fold properly given its sequence divergence.

Another odd gene structure found in both *Cuscuta* species is the final portion of *nad1*, namely exon 4, *matR* and exon 5 (see Fig. S6 Additional file 3). In the two *Cuscuta* assemblies, exon 5 is found in the reverse orientation on the opposite strand, while exon 4 is located on the same strand as *matR*. This may indicate nad1i728 has shifted to trans-splicing (as seen in some other angiosperms, e.g. *Nicotiana tabacum*, NC_006581.1), but if that is the case, it seems odd that exon 5 is in such close proximity to the rest of the intron and exon 4.

### Potential intracellular transfers

Using reference plastomes, we plotted plastid hits on the mitogenome assemblies (see Figs. S7 and S8, Additional file3). Excluding hits that occurred in the ribosomal genes or *atp1*, which are similar to plastid ribosomal genes and *atpA*, we found 8 regions with hits, representing approximately 0.6 and 1.1% of the mitogenomes of *C. australis* and *C. campestris*, respectively (Table 2).

**Table 2.**
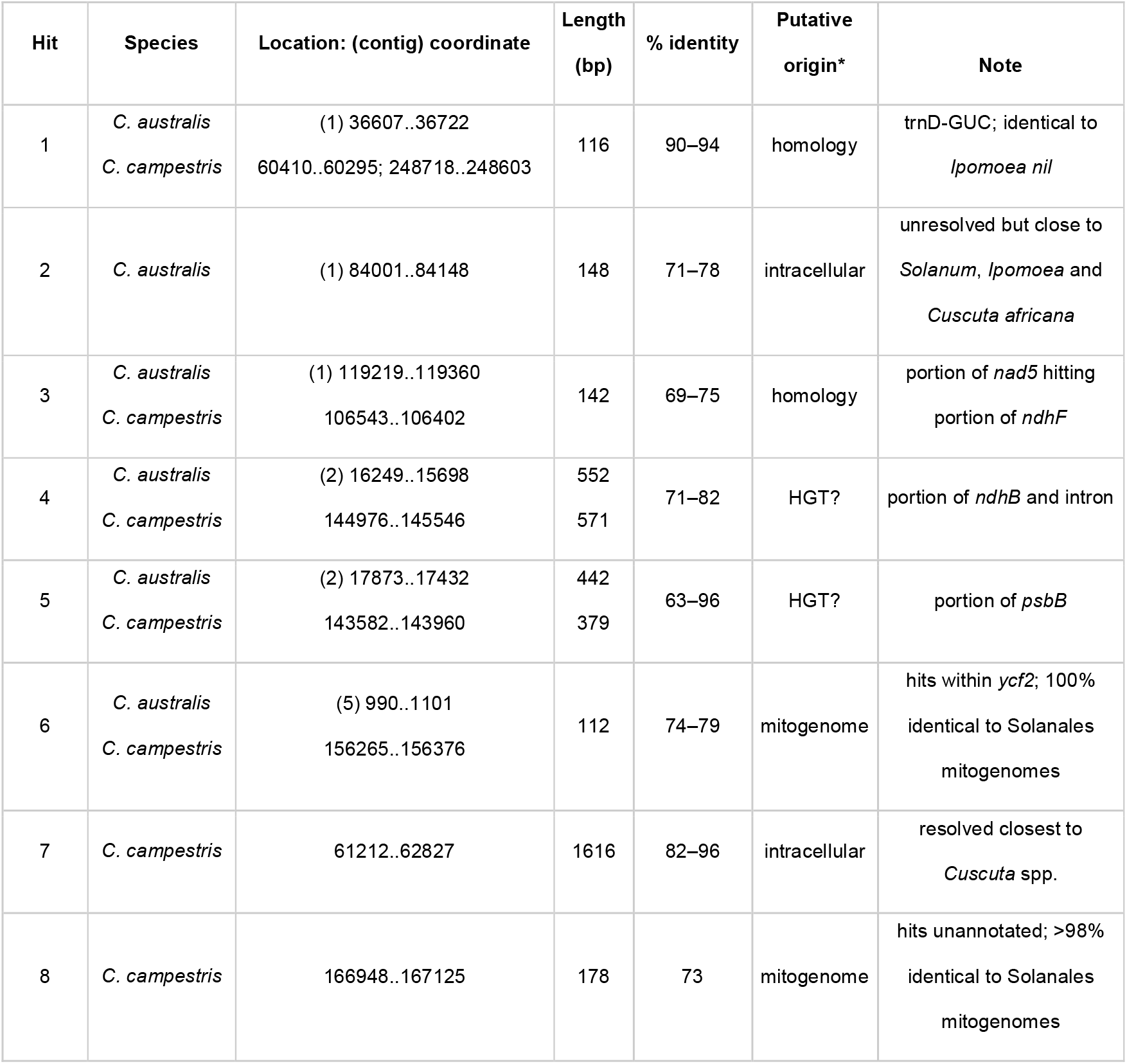
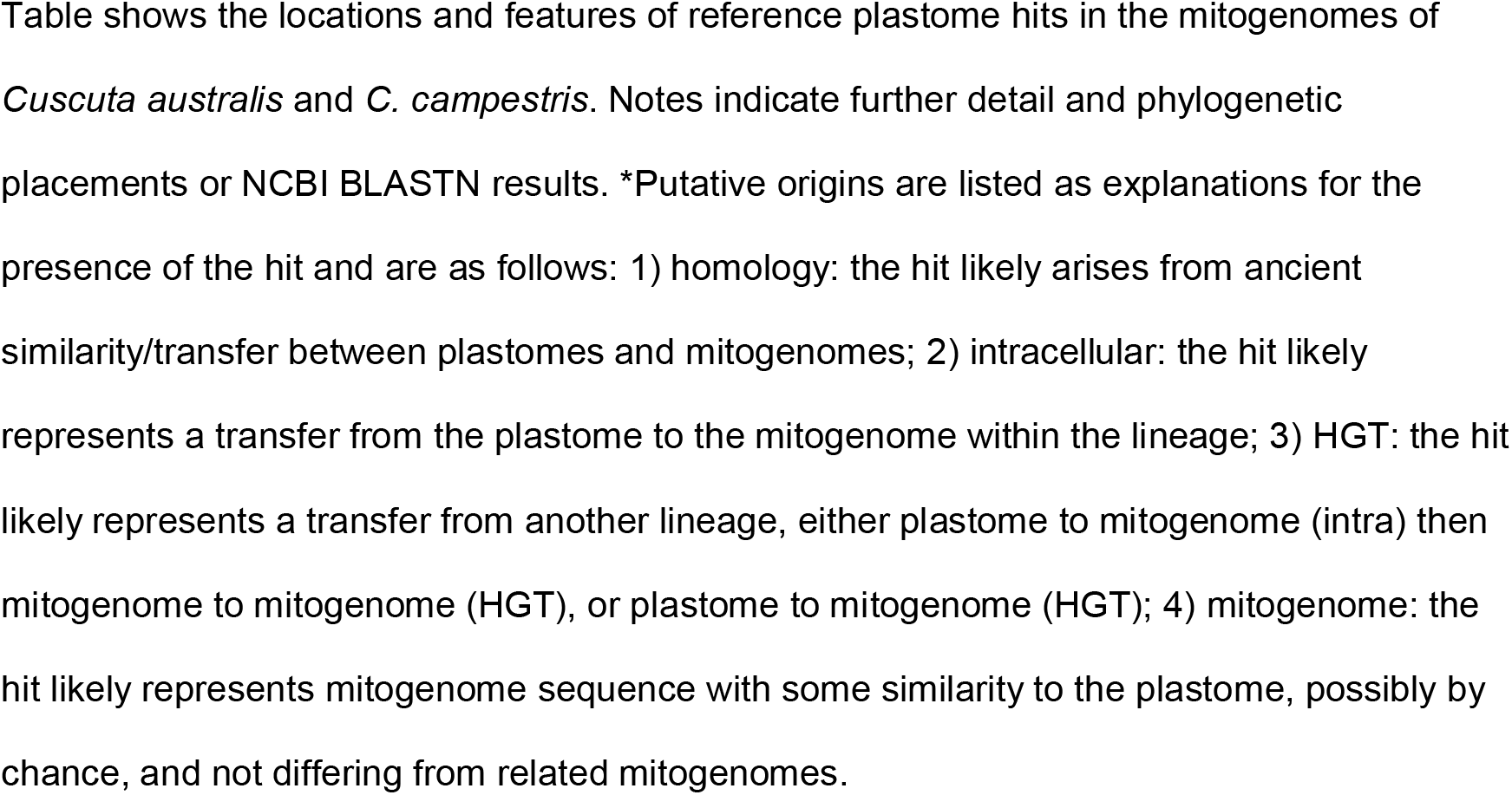
Reference plastome hits in the mitogenomes of *Cuscuta australis* and *C. campestris*.

Phylogenetic trees for hits 2 and 7 (see Fig. S9, Additional file 3) support the mitochondrial sequences grouping with *Cuscuta* or *Ipomoea* chloroplast sequences, consistent with intracellular transfer. Both species share two regions corresponding to HMMER hits to plastid genes *ndhB* and *psbB* (hits 4 and 5). Phylogenetic analysis of hits 4 and 5 are partly unresolved but suggestive of potential HGT, as the two *Cuscuta* species lack intact copies of *ndhB* in their plastomes (hit 4) and the *psbB* hits group closer to Malpighiales than other *Cuscuta* chloroplast sequences. In both *Cuscuta* species, the *psbB* and *ndhB* hit regions are within 2 kb of each other, possibly indicating a larger transferred region. Bootstrap support is low in the *ndhB* tree, with taxa from asterids and rosids, making it unclear whether the two hits are associated with the same potential donor lineage.

Searches of the nuclear genomes produced markedly different results for the two *Cuscuta* species. In *C. australis*, the ~260 Mbp assembly hit the mitochondrial assembly for cumulatively ~27% of its length, but in *C. campestris*, the ~477 Mbp assembly hit the mitochondrial assembly for ~99.8% of its length or essentially the entire assembly. Three scaffolds with lengths of 172, 43 and 21 kb are essentially entirely mitochondrial. These scaffolds could tentatively indicate recent large-scale integration of mitochondrial sequences into the nuclear genome, but could also be simple assembly artifacts since mitochondrial contaminations are difficult to identify.

Potential candidates for missing mitochondrial genes were identified in both nuclear genomes for *rps2, rps10, rps11, rps14, rps19, sdh3* and *sdh4* (Table 3). All the candidates had intact open reading frames except for *sdh3*, which had no determined start codon. Phylogenetic analysis of the candidates with copies from the reference mitogenomes and top NCBI BLASTN hits consistently put the nuclear copies on longer branches relative to the mitogenome copies (see Figs. S10–13, Additional file 3). In some cases, the trees suggest vertical inheritance of a nuclear copy (*rps2, rps11, rps14, sdh3*) but in three cases it is unclear and could represent HGT (*rps10, rps19, sdh4*), though long branch attraction may obscure phylogenetic relationships. Searching for indications of mitochondrial targeting sequences with Predotar v. 1.04 [79] indicated possible mitochondrial targeting (p 0.21–0.44) for *rps10, rps14, rps19* and *sdh3* (if an ‘M’ is added in place of the undetermined start codon).

**Table 3.** Nuclear candidate copies of missing mitochondrial genes in *Cuscuta australis* and *C. campestris*.

| Mitochondrial gene | Nuclear copy length (bp) | Reference average length (bp) | Putative origin |
| --- | --- | --- | --- |
| *rps2* | 618 | 978 | vertical; nuclear copy or UTP--glucose-1-phosphate uridylyltransferase |
| *rps10* | 387 | 354 | unclear; possible HGT |
| *rps11* | 432 | 486 | vertical; nuclear copy |
| *rps14* | 321 | 295 | vertical; nuclear copy |
| *rps19* | 273 | 286 | unclear; possible HGT |
| *sdh3* | 321, 324 | 343 | vertical; nuclear copy |
| *sdh4* | 288 | 418 | unclear; possible HGT |

Table showing the nuclear candidate copies, including the lengths of reference mitochondrial genes. The putative origin is based on phylogenetic analysis of top NCBI BLASTN hits together with reference mitogenomes.

### Horizontal gene transfer

Phylogenetic analysis of the concatenation of all mitochondrial genes strongly supports the grouping of *Cuscuta* within Solanales (Fig. 3). Based on the individual gene trees with sampling across seed plants, the majority of genes (26/34, 76%) in the three *Cuscuta* species are strongly supported (>80% bootstrap support) as grouping together and sister to their close autotrophic relative *Ipomoea nil*, a pattern consistent with vertical rather than horizontal inheritance (see Fig. S14, Additional file 4). Exceptions to this pattern are *atp9* (71% sister to *Ipomoea), ccmB* (unsupported), *cox2* (66%), *nad3* (unsupported), *nad4L* (not sister to *Ipomoea*, but 80% support together with *Ipomoea* in Solanales), *rpl16* (68%), *rps12* (unsupported), and *rrn5* (unresolved) (note that *ccmF_C_* and *rrn5* from *C. gronovii* were not included). In the cases with no support for a sister relationship with *Ipomoea*, there is no strong support for a grouping elsewhere, as would be expected for HGT.

**Fig. 3.**
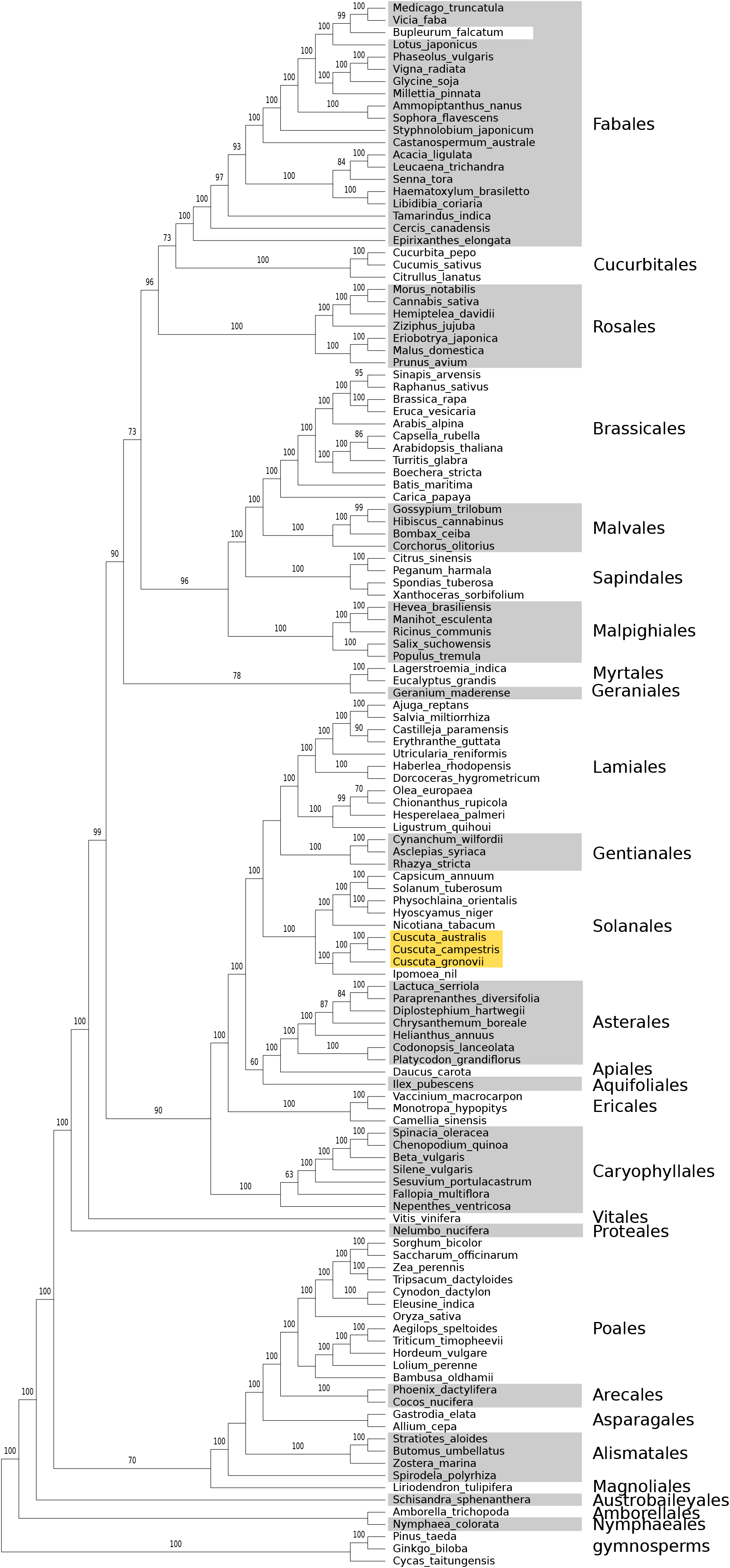
Maximum likelihood cladogram based on concatenation of all known mitochondrial genes present in *Cuscuta* together with representatives from angiosperm orders with fully assembled mitogenomes. Orders are indicated at the right along with the outgroup gymnosperms, and the location of *Cuscuta* is highlighted in yellow. Bootstrap support of 60% or greater is shown above branches. The sequence for *Bupleurum falcatum* (NC_035962.1) is likely misidentified.

Intergenic sequences generally lacked clear evidence of longer (>500 bp) contiguous groups of hits to single orders outside Solanales (see Figs. S7 and S8, Additional file 3). Some exceptions could be identified, including hits to Asterales, Caryophyllales, Fabales, Gentianales, Lamiales and Vitales, totalling approximately 7.4 and 4.7% of the mitogenomes of *C. australis* and *C. campestris*, respectively.

In portions of some genes, blast hits with higher identity to other orders than to Solanales were found (*cox2* exon 1, *nad4* exon 3, and *nad5* exon 5), so we ran GENECONV to detect possible conversion. In two cases there were some sequences showing significant (P < 0.05) signal of conversion (4 sequences for *cox2* and 2 for *nad5*), but not involving the *Cuscuta* sequences. For *nad4*, there were many significant comparisons, but all but one involved *Vicia faba* (KC189947), suggesting it may have undergone gene conversion. There was no significant indication that the *Cuscuta* sequences were chimeric.

## Discussion

Recently sequenced genomes of parasitic *Cuscuta* species revealed extensive HGT events into their nuclear genomes from a broad range of non-parasitic angiosperms that serve as their hosts. It was a reasonable expectation that there would be high transfer activity of genetic material into the mitogenomes of these parasites as well. The organellar genomes of the two *Cuscuta* species that we have assembled and annotated here, however, are not clearly different in gene content from other *Cuscuta* plastomes or typical angiosperm mitogenomes. Based on phylogenetic analysis, we found little evidence for HGT into the mitogenomes of the two species, an unexpected result given the parasitic lifestyle of *Cuscuta* and evidence for HGT candidates in their nuclear genomes. In the process of examining the mitogenomes, we also identified a novel structure of the *ccmF_C_* gene in *Ipomoea* and *Cuscuta*, indicative of a shift to trans-splicing of the group II intron and possibly a shorter product.

### Gene losses and possible intracellular transfers

All of the ten genes missing from the two *Cuscuta* mitogenomes are commonly found to be lost during angiosperm evolution [80, 81]. That *rps2* and *rps11* are missing is consistent with the inference of their ancient losses in angiosperms [80]. For seven of the ten missing or pseudogenized genes, potential candidates for nuclear copies were found. Transfers of mitochondrial genes to the nucleus have been inferred to be numerous across plant evolution and seem to be the primary mode of functional gene loss from the mitogenome [82]. Whether the gene products from the nuclear candidates are functional or imported into the mitochondrion is uncertain, but tests for targeting sequences suggested some of them might be targeted to the mitochondrion. The nuclear candidates are on longer branches in phylogenetic trees compared to the mitochondrial copies, suggesting higher substitution rates. This is consistent with typical observations of higher substitution rates in the nuclear genome compared to the mitogenome [83], and of higher rates in nuclear copies of genes transferred from the mitogenome compared to the mitogenome copies [84], though exceptions exist, such as in *Silene* [85].

The three missing mitochondrial genes not detected in the nuclear genomes are *rpl2, rpl10* and *rps7*. Losses of *rpl2* and *rps7* are frequent among angiosperms, but only for *rpl2* are losses commonly associated with functional transfers to the nucleus [36, 80, 86, 87]. Losses of *rpl10* are less common and have been associated with functional replacement by a duplicated nuclear gene normally encoding chloroplast RPL10 in some monocot and Brassicaceae lineages [81, 88, 89]. Searching ORFs from both nuclear genomes for possible copies of RPL10 suggested the presence of one (*C. australis*) or two (*C. campestris*) genes more similar to the chloroplast-targeted copy from *Arabidopsis* (NP_196855) than to the mitochondrial-targeted one (NP_187843). Targeting predictions from Predotar for the copies in *Cuscuta* provided no support for targeting the chloroplast or the mitochondrion.

Plastid transfers appear to be limited in the mitogenomes, with a 148 bp hit in *C. australis* and a 1.6 kb hit in *C. campestris*. The other main candidate regions included hits to *psbB* and *ndhB*, but based on phylogenetic analysis appear to be more similar to regions in distantly related plants. The trees are inconclusive as to the origin of the region, but suggest a horizontal transfer, possibly via an intracellular plastome transfer to the donor’s mitogenome and then a plant to plant transfer between mitogenomes, a common route for plastome to mitogenome HGT [90]. The close association of the *psbB* and *ndhB* hits might indicate a transfer of a larger contiguous block of DNA from a donor with the genes in close proximity, but this is speculative and the donor lineage is not clear from our search and trees.

### Lack of horizontal gene transfer

Phylogenetic evidence for HGT in the *Cuscuta* mitogenome assemblies is lacking, with only a few genes having <80% bootstrap support for a close relationship to *Ipomoea nil*, and those exceptions also with negligible support for alternative relationships. BLASTN-based assessment of intergenic sequences did uncover a few longer candidates for possible HGT, but it is unclear how reliable the similarity search is for uncovering the origin of those sequences. It is possible they represent vertically-inherited mitogenomic sequences. Given the dynamic nature of mitogenome size and structure in angiosperms [85, 91–93], our limited sampling of lineages may not include more similar sequences from more closely related plants. The HGT candidates also lack embedded genes for the most part, making it difficult to determine homology and to further support an origin. In cases where BLAST hits to another order cover genes (*cox3*, *nad4L, rps13*), phylogenetic analysis of the genes generally contradicts the BLAST-based assignment. The aforementioned presence of sequences of chloroplast origin similar to regions in distantly related plants may also indicate that some HGT has occurred, possibly earlier in the evolution of *Cuscuta* given the region is shared by all the *Cuscuta* sequences we examined.

Why the mitogenomes lack clear evidence of HGT is unclear, given it is prevalent in the nuclear genome of *Cuscuta*. One possible explanation may be related to the size of the mitogenomes we have assembled, which are on the low side for angiosperms. Based on the reference set of mitogenomes (with 168 angiosperms), the average size of the angiosperm mitogenomes is roughly 670 kb, while the assemblies are less than half that. Given there has been a correspondence observed between mitogenome size and the amount of incorporated plastid transfers [90] and that plant mitochondrial genomes appear to have relaxed restriction on the size of intergenic spacers [e.g. 94], it may be that larger mitogenomes with longer intergenic regions allow for the incorporation of more foreign DNA. Conversely, the smaller mitogenomes in *Cuscuta* may experience more restriction on what can be successfully incorporated. Other explanations could include population sizes and demographic effects that make it more or less likely to retain incorporations of foreign DNA in a species.

It is unclear why there has not been an extensive expansion of intergenic material in *Cuscuta* or *Ipomoea* for that matter (265,768 bp). The wide range of sizes of angiosperm mitogenomes are difficult to explain, but recombination rates have been put forward as one potential factor, albeit a complex one [85]. In a study of the massive differences in size amongst *Silene* mitogenomes, Sloan et al. [85] found that the larger mitogenomes experienced lower recombinational activity (<20%) and had more divergent repeats, suggesting reduced gene conversion. The assembled mitogenomes had relatively high recombinational activity across longer repeat sequences as assessed with long reads (often >20% and >40% for *C. australis* and *C. campestris*, respectively), though with small sample sizes. The difficulty in assembling the mitogenomes and the multiple potential paths through the assemblies are also consistent with multiple conformations across highly similar or identical repeats, possibly indicating ongoing recombination and conversion.

### Curious structure of *ccmF_C_* and its trans-splicing intron

Both *Cuscuta* species and their close autotrophic relative *Ipomoea nil* appear to share a different structure of their *ccmF_C_* genes than other plants. In all three cases, the first exon is split into two pieces, and the same has happened for the group II intron, making transsplicing of the intron necessary in order to join the now remotely located parts of the gene. In angiosperm mitogenomes, most introns are group II introns (having six domains) and require multiple protein factors, including some encoded in the nuclear genome, for proper splicing [95]. While the exact mechanism of trans-splicing for group II introns and the splicing factors required remain unclear, this type of splicing has been observed in a number of other angiosperm mitochondrial genes, namely *nad1, nad2* and *nad5* [96] and *cox2* [97]. Typically, the break point for the two pieces of the intron that are trans-spliced is in domain IV [96, 98], which is where the intron is broken in *Cuscuta* and *Ipomoea*. The evidence presented here, namely the mRNA reads that indicate a product joined from two disparate locations in the genome and the break in domain IV, suggests this is the fifth such example of a gene with a group II intron that has shifted to trans-splicing.

Angiosperm *ccmF_C_* genes examined so far typically have intact rather than broken group II introns. For splicing of the intact intron, an essential nuclear-encoded splicing factor, WTF9, has been identified which shares an RNA-binding domain with another factor involved in intron splicing in plastids [99]. In examining the role of WTF9, des Francs-Small et al. [99] also elucidated the role of *ccmF_C_* in cytochrome biogenesis, showing that proper splicing of the intron for the formation of a complete protein was essential to the generation of cytochromes c and c1 in *Arabidopsis*. In addition, splicing-impaired mutants had higher levels of alternative oxidase, suggesting problems with the respiratory chain [99]. It appears that being able to produce a fully intact *ccmF_C_* product is important for respiratory function, which makes the break in the first exon in *Cuscuta* and *Ipomoea* puzzling. While there is evidence that the intron is trans-spliced and connects the 3’ portion of exon 1 with exon 2, there is no indication that the larger 5’ portion of exon 1 is connected for translation into the final product. Transcriptomic reads show RNA editing in all parts of the gene (see Fig. 2 and Fig. S4, Additional file 2), similar to predictions from PREP-MT, but whether this indicates an intact product is unclear. Pseudogenization of *ccmF_C_* has only been reported for *Silene conica* amongst angiosperms [85], though there was indication that a truncated form of the protein might still be expressed and functional. Normally, the full *ccmF_C_* product is thought to be integrated into a larger complex potentially involved in heme ligation [100]. Another possible explanation for the process might be a continued trend of gene breakup, similar to the fission of *ccmF_N_* in *Trifolium*, Brassicaceae and *Allium* [101]. The functionality of at least part of the *ccmF_C_* gene would be consistent with its conservation in both *Ipomoea* and *Cuscuta*, though their copies are divergent relative to other members of Solanales (see Fig. S14, Additional file 4). Gene loci coding for homologs of the WTF9 splicing factor are present in *C. campestris* and *C. australis* (data not shown), excluding this as a reason for the switch. How *Cuscuta* and *Ipomoea* are able to function effectively with their *ccmF_C_* structure is unclear, and additional work would be helpful to characterise and evaluate the *ccmF_C_* protein products in this group of plants.

While the assembly approach is not guaranteed to have captured the entirety of the mitogenomes (for instance, the *Cuscuta australis* assembly consists of multiple contigs, and the *C. campestris* assembly was circularized from multiple smaller circular contigs), the search approach makes it unlikely that any missing sequences would carry substantially different phylogenetic signals of HGT than the bulk of the assemblies or that they would harbour intact foreign genes or a contiguous copy of *ccmF_C_*. Alternative structural arrangement of the contigs and construction of the assemblies are also unlikely to affect our conclusions on HGT, but could have some impact on how we have interpreted the structure of *ccmF_C_*. The lack of mRNA support for a fully intact *ccmF_C_* product, however, supports the assembly and interpretation of a fragmented gene structure.

## Conclusions

Our search of the assembled mitogenomes of *Cuscuta australis* and *C. campestris* revealed little to no indication of HGT, contrary to our expectations based on examples from their nuclear genomes. This finding suggests that elevated rates of HGT into mitogenomes, where the bulk of examples from parasitic plants have been found to date, is not always the case. The observed amount of HGT may also be influenced by species-specific factors such as mitogenome size and recombination frequency that allow for any DNA uptake to be retained.

## Abbreviations

CDS: coding sequence
HGT: horizontal gene transfer
ORF: open reading frame

## Declarations

### Ethics approval and consent to participate

Not applicable.

### Consent for publication

Not applicable.

### Availability of data and materials

Organelle genomes are deposited in GenBank under accessions BK059221 and BK059222 for the *C. australis* and *C. campestris* plastomes, and BK059197–BK059204 and BK016277 for the *C. australis* and *C. campestris* mitogenomes, respectively.

Accession numbers for reference sequences downloaded from Genbank are provided in Additional file 1. Sequence alignments used in phylogenetic analysis are included as Additional file 5.

### Authors’ contributions

BMA and GP conceived and designed the study. KK provided access to genomic data. BMA conducted bioinformatic assembly and analysis. BMA drafted the manuscript. All authors contributed to critically revising the manuscript. All authors read and approved the final version.

## Acknowledgements

Not applicable.

## Additional files

Additional file 1: supplementary Tables S1 and S2. (pdf)

Additional file 2: supplementary Figs. S1–S5, Tables S3 and S4. (pdf)

Additional file 3: supplementary Figs. S6–S13. (pdf)

Additional file 4: supplementary Fig. S14. (pdf)

Additional file 5: fasta alignments used in phylogenetic analyses. (zip)

