## Additional file 1 for "Mitochondrial genomes of two parasitic *Cuscuta* species lack clear evidence of horizontal gene transfer and retain unusually fragmented *ccmF_C_* genes"

### Additional file 1: supplementary Tables S1 and S2

Table S1. Reference plastomes used for annotation and BLASTN searches and their GenBank accession numbers.

| **Order** | **Family** | **Genus** | **Specific_ep** | **Accession number** |
|---|---|---|---|---|
| Asterales | Asteraceae | Helianthus | annuus | NC_007977.1 |
| Boraginales | Lennoaceae | Pholisma | arenarium | NC_039719.1 |
| Cucurbitales | Cucurbitaceae | Cucurbita | pepo | NC_038229.1 |
| Ericales | Theaceae | Camellia | petelotii | NC_024661. |
| Fabales | Fabaceae | Glycine | max | NC_007942.1 |
| Lamiales | Lamiaceae | Ocimum | basilicum | NC_035143.1 |
| Magnoliales | Magnoliaceae | Liriodendron | tulipifera | NC_008326.1 |
| Malvales | Malvaceae | Gossypium | barbadense | NC_008641.1 |
| Poales | Poaceae | Oryza | sativa | NC_008155.1 |
| Proteales | Nelumbonaceae | Nelumbo | nucifera | NC_025339.1 |
| Santalales | Schoepfiaceae | Schoepfia | jasminodora | NC_034228.1 |
| Solanales | Convolvulaceae | Cuscuta | exaltata | NC_009963.1 |
| Solanales | Convolvulaceae | Cuscuta | obtusiflora | NC_009949.1 |
| Vitales | Vitaceae | Vitis | vinifera | NC_007957.1 |

Table S2. Reference mitogenomes used for annotation, BLASTN searches and phylogenetic analysis and their GenBank accession numbers.

| **Order** | **Family** | **Genus** | **Specific_ep** | **Accession number** |
|---|---|---|---|---|
| Alismatales | Araceae | Spirodela | polyrhiza | NC_017840.1 |
| Alismatales | Butomaceae | Butomus | umbellatus | NC_021399.1 |
| Alismatales | Hydrocharitaceae | Stratiotes | aloides | NC_035317.1 |
| Alismatales | Zosteraceae | Zostera | marina | NC_035345.1 |
| Amborellales | Amborellaceae | Amborella | trichopoda | KF754799.1–803.1 |
| Apiales | Apiaceae | Bupleurum | falcatum | NC_035962.1 |
| Apiales | Apiaceae | Daucus | carota | NC_017855.1 |
| Aquifoliales | Aquifoliaceae | Ilex | pubescens | NC_045078.1 |
| Arecales | Arecaceae | Cocos | nucifera | NC_031696.1 |
| Arecales | Arecaceae | Phoenix | dactylifera | NC_016740.1 |
| Asparagales | Amaryllidaceae | Allium | cepa | NC_030100.1 |
| Asparagales | Orchidaceae | Gastrodia | elata | MF070084.1–102.1 |
| Asterales | Asteraceae | Chrysanthemum | boreale | NC_039757.1 |
| Asterales | Asteraceae | Diplostephium | hartwegii | NC_034354.1 |
| Asterales | Asteraceae | Helianthus | annuus | NC_023337.1 |
| Asterales | Asteraceae | Lactuca | saligna | NC_042406.1 |
| Asterales | Asteraceae | Lactuca | sativa | NC_042756.1 |
| Asterales | Asteraceae | Lactuca | serriola | NC_042378.1 |
| Asterales | Asteraceae | Paraprenanthes | diversifolia | MN661146.1 |
| Asterales | Campanulaceae | Codonopsis | lanceolata | NC_037949.1 |
| Asterales | Campanulaceae | Platycodon | grandiflorus | NC_035958.1 |
| Austrobaileyales | Schisandraceae | Schisandra | sphenanthera | NC_042758.1 |
| Brassicales | Bataceae | Batis | maritima | NC_024429.1 |
| Brassicales | Brassicaceae | Arabidopsis | thaliana | NC_037304.1 |
| Brassicales | Brassicaceae | Arabis | alpina | NC_037070.1 |
| Brassicales | Brassicaceae | Boechera | stricta | NC_042143.1 |
| Brassicales | Brassicaceae | Brassica | carinata | NC_016120.1 |
| Brassicales | Brassicaceae | Brassica | juncea | NC_016123.1 |
| Brassicales | Brassicaceae | Brassica | napus | NC_008285.1 |
| Brassicales | Brassicaceae | Brassica | nigra | NC_029182.1 |
| Brassicales | Brassicaceae | Brassica | oleracea | NC_016118.1 |
| Brassicales | Brassicaceae | Brassica | oxyrrhina | AP018041.1 |
| Brassicales | Brassicaceae | Brassica | rapa | NC_016125.1 |
| Brassicales | Brassicaceae | Capsella | rubella | NC_042883.1 |
| Brassicales | Brassicaceae | Eruca | vesicaria | KF442616.1 |
| Brassicales | Brassicaceae | Raphanus | sativus | NC_018551.1 |

Table S2 continued.

| Order | Family | Genus | Specific_ep | Accession number |
|---|---|---|---|---|
| Brassicales | Brassicaceae | Schrenkiella | parvula | KT988071.2 |
| Brassicales | Brassicaceae | Sinapis | arvensis | NC_031896.1 |
| Brassicales | Brassicaceae | Turritis | glabra | LC325489.1 |
| Brassicales | Caricaceae | Carica | papaya | NC_012116.1 |
| Caryophyllales | Aizoaceae | Sesuvium | portulacastrum | MN683736.1 |
| Caryophyllales | Caryophyllaceae | Silene | conica | JF750490.1–629.1 |
| Caryophyllales | Caryophyllaceae | Silene | latifolia | NC_014487.1 |
| Caryophyllales | Caryophyllaceae | Silene | noctiflora | JF750431.1–489.1 |
| Caryophyllales | Caryophyllaceae | Silene | vulgaris | JF750427.1–30.1 |
| Caryophyllales | Chenopodiaceae | Beta | macrocarpa | NC_015994.1 |
| Caryophyllales | Chenopodiaceae | Beta | vulgaris | NC_002511.2 |
| Caryophyllales | Chenopodiaceae | Chenopodium | quinoa | NC_041093.1 |
| Caryophyllales | Chenopodiaceae | Spinacia | oleracea | NC_035618.1 |
| Caryophyllales | Nepenthaceae | Nepenthes | ventricosa | NC_039531.1 |
| Caryophyllales | Polygonaceae | Fallopia | multiflora | MF611850.1–1.1 |
| Cucurbitales | Cucurbitaceae | Citrullus | lanatus | NC_014043.1 |
| Cucurbitales | Cucurbitaceae | Cucumis | melo | MG947207.1–9.1 |
| Cucurbitales | Cucurbitaceae | Cucumis | sativus | NC_016004.1–6.1 |
| Cucurbitales | Cucurbitaceae | Cucurbita | pepo | NC_014050.1 |
| Cycadales | Cycadaceae | Cycas | taitungensis | NC_010303.1 |
| Ericales | Ericaceae | Monotropa | hypopitys | MK990822.1–3.1 |
| Ericales | Ericaceae | Vaccinium | macrocarpon | NC_023338.1 |
| Ericales | Theaceae | Camellia | sinensis | NC_043914.1 |
| Fabales | Fabaceae | Acacia | ligulata | NC_040998.1 |
| Fabales | Fabaceae | Ammopiptanthus | mongolicus | NC_039660.1 |
| Fabales | Fabaceae | Ammopiptanthus | nanus | MH127920.1 |
| Fabales | Fabaceae | Castanospermum | australe | MK426679.1 |
| Fabales | Fabaceae | Cercis | canadensis | MN017226.1 |
| Fabales | Fabaceae | Glycine | max | NC_020455.1 |
| Fabales | Fabaceae | Glycine | soja | NC_039768.1 |
| Fabales | Fabaceae | Haematoxylum | brasiletto | NC_045040.1 |
| Fabales | Fabaceae | Leucaena | trichandra | NC_039738.1 |
| Fabales | Fabaceae | Libidibia | coriaria | NC_045039.1 |
| Fabales | Fabaceae | Lotus | japonicus | NC_016743.2 |
| Fabales | Fabaceae | Medicago | truncatula | NC_029641.1 |
| Fabales | Fabaceae | Millettia | pinnata | NC_016742.1 |
| Fabales | Fabaceae | Phaseolus | vulgaris | NC_045135.1 |
| Fabales | Fabaceae | Senna | occidentalis | NC_038221.1 |

Table S2 continued.

| Order | Family | Genus | Specific_ep | Accession number |
|---|---|---|---|---|
| Fabales | Fabaceae | Senna | tora | NC_038053.1 |
| Fabales | Fabaceae | Sophora | flavescens | NC_043897.1 |
| Fabales | Fabaceae | Styphnolobium | japonicum | NC_039596.1 |
| Fabales | Fabaceae | Tamarindus | indica | NC_045038.1 |
| Fabales | Fabaceae | Vicia | faba | KC189947.1 |
| Fabales | Fabaceae | Vigna | angularis | NC_021092.1 |
| Fabales | Fabaceae | Vigna | radiata | NC_015121.1 |
| Fabales | Polygalaceae | Epirixanthes | elongata | MG783394.1 |
| Gentianales | Apocynaceae | Asclepias | syriaca | NC_022796.1 |
| Gentianales | Apocynaceae | Cynanchum | wilfordii | MF611847.1–9.1 |
| Gentianales | Apocynaceae | Rhazya | stricta | NC_024293.1 |
| Geraniales | Geraniaceae | Geranium | maderense | NC_027000.1 |
| Ginkgoales | Ginkgoaceae | Ginkgo | biloba | NC_027976.1 |
| Lamiales | Gesneriaceae | Boea | hygrometrica | NC_016741.1 |
| Lamiales | Gesneriaceae | Haberlea | rhodopensis | MH757117.1 |
| Lamiales | Lamiaceae | Ajuga | reptans | NC_023103.1 |
| Lamiales | Lamiaceae | Salvia | miltiorrhiza | NC_023209.1 |
| Lamiales | Lentibulariaceae | Utricularia | reniformis | NC_034982.1 |
| Lamiales | Oleaceae | Chionanthus | rupicola | MG372115.1 |
| Lamiales | Oleaceae | Hesperelaea | palmeri | NC_031323.1 |
| Lamiales | Oleaceae | Ligustrum | quihoui | MN723864.1 |
| Lamiales | Oleaceae | Olea | europaea | MG372117.1 |
| Lamiales | Orobanchaceae | Castilleja | paramensis | NC_031806.1 |
| Lamiales | Phrymaceae | Mimulus | guttatus | NC_018041.1 |
| Magnoliales | Magnoliaceae | Liriodendron | tulipifera | NC_021152.1 |
| Malpighiales | Euphorbiaceae | Hevea | brasiliensis | AP014526.1 |
| Malpighiales | Euphorbiaceae | Manihot | esculenta | NC_045136.1 |
| Malpighiales | Euphorbiaceae | Ricinus | communis | NC_015141.1 |
| Malpighiales | Salicaceae | Populus | alba | NC_041085.1 |
| Malpighiales | Salicaceae | Populus | davidiana | NC_035157.1 |
| Malpighiales | Salicaceae | Populus | tremula | NC_028096.1 |
| Malpighiales | Salicaceae | Salix | purpurea | NC_029693.1 |
| Malpighiales | Salicaceae | Salix | suchowensis | NC_029317.1 |
| Malvales | Malvaceae | Bombax | ceiba | NC_038052.1 |
| Malvales | Malvaceae | Corchorus | capsularis | NC_031359.1 |
| Malvales | Malvaceae | Corchorus | olitorius | NC_031360.1 |
| Malvales | Malvaceae | Gossypium | arboreum | NC_035073.1 |

Table S2 continued.

| Order | Family | Genus | Specific_ep | Accession number |
|---|---|---|---|---|
| Malvales | Malvaceae | Gossypium | barbadense | NC_028254.1 |
| Malvales | Malvaceae | Gossypium | davidsonii | NC_035075.1 |
| Malvales | Malvaceae | Gossypium | harknessii | NC_027407.1 |
| Malvales | Malvaceae | Gossypium | hirsutum | NC_027406.1 |
| Malvales | Malvaceae | Gossypium | raimondii | NC_029998.1 |
| Malvales | Malvaceae | Gossypium | thurberi | NC_035074.1 |
| Malvales | Malvaceae | Gossypium | trilobum | NC_035076.1 |
| Malvales | Malvaceae | Hibiscus | cannabinus | NC_035549.1 |
| Myrtales | Lythraceae | Lagerstroemia | indica | NC_035616.1 |
| Myrtales | Myrtaceae | Eucalyptus | grandis | NC_040010.1 |
| Nymphaeales | Nymphaeaceae | Nymphaea | colorata | NC_037468.1 |
| Pinales | Pinaceae | Pinus | Taeda | NC_039746.1 |
| Poales | Poaceae | Aegilops | speltoides | NC_022666.1 |
| Poales | Poaceae | Bambusa | oldhamii | EU365401.1 |
| Poales | Poaceae | Cynodon | dactylon | MK175054.1 |
| Poales | Poaceae | Eleusine | indica | NC_040989.1 |
| Poales | Poaceae | Hordeum | vulgare | MN127975.1 |
| Poales | Poaceae | Lolium | perenne | JX999996.1 |
| Poales | Poaceae | Oryza | coarctata | MG429050.1 |
| Poales | Poaceae | Oryza | minuta | NC_029816.1 |
| Poales | Poaceae | Oryza | rufipogon | NC_013816.1 |
| Poales | Poaceae | Oryza | sativa | NC_007886.1 |
| Poales | Poaceae | Saccharum | officinarum | LC107874.1–5.1 |
| Poales | Poaceae | Sorghum | bicolor | NC_008360.1 |
| Poales | Poaceae | Tripsacum | dactyloides | NC_008362.1 |
| Poales | Poaceae | Triticum | aestivum | NC_036024.1 |
| Poales | Poaceae | Triticum | timopheevii | NC_022714.1 |
| Poales | Poaceae | Zea | luxurians | NC_008333.1 |
| Poales | Poaceae | Zea | mays | NC_007982.1 |
| Poales | Poaceae | Zea | perennis | NC_008331.1 |
| Proteales | Nelumbonaceae | Nelumbo | nucifera | NC_030753.1 |
| Rosales | Cannabaceae | Cannabis | sativa | NC_029855.1 |
| Rosales | Moraceae | Morus | notabilis | NC_041177.1 |
| Rosales | Rhamnaceae | Ziziphus | jujuba | NC_029809.1 |
| Rosales | Rosaceae | Eriobotrya | japonica | NC_045228.1 |
| Rosales | Rosaceae | Malus | hupehensis | KR534606.1 |
| Rosales | Rosaceae | Malus | x-domestica | NC_018554.1 |

Table S2 continued.

| Order | Family | Genus | Specific_ep | Accession number |
|---|---|---|---|---|
| Rosales | Rosaceae | Prunus | avium | NC_044768.1 |
| Rosales | Ulmaceae | Hemiptelea | davidii | MN061667.1 |
| Santalales | Balanophoraceae | Lophophytum | mirabile | KU992322.1–80.1; KX792461.1 |
| Santalales | Viscaceae | Viscum | album | NC_029039.1 |
| Santalales | Viscaceae | Viscum | scurruloideum | KT022222.1–3.1 |
| Sapindales | Anacardiaceae | Spondias | mombin | NC_045035.1 |
| Sapindales | Anacardiaceae | Spondias | tuberosa | NC_045036.1 |
| Sapindales | Nitrariaceae | Peganum | harmala | MK431826.1 |
| Sapindales | Rutaceae | Citrus | sinensis | NC_037463.1 |
| Sapindales | Sapindaceae | Xanthoceras | sorbifolium | MK333231.1 |
| Saxifragales | Cynomoriaceae | Cynomorium | coccineum | KX270753.1–801.1 |
| Solanales | Convolvulaceae | Ipomoea | nil | NC_031158.1 |
| Solanales | Solanaceae | Capsicum | annuum | NC_024624.1 |
| Solanales | Solanaceae | Hyoscyamus | niger | NC_026515.1 |
| Solanales | Solanaceae | Nicotiana | attenuata | NC_036467.1 |
| Solanales | Solanaceae | Nicotiana | sylvestris | NC_029805.1 |
| Solanales | Solanaceae | Nicotiana | tabacum | NC_006581.1 |
| Solanales | Solanaceae | Physochlaina | orientalis | NC_044153.1 |
| Solanales | Solanaceae | Solanum | commersonii | MF989960.1–1.1 |
| Solanales | Solanaceae | Solanum | lycopersicum | NC_035963.1 |
| Solanales | Solanaceae | Solanum | pennellii | NC_035964.1 |
| Solanales | Solanaceae | Solanum | tuberosum | MN104801.1–3.1 |
| Vitales | Vitaceae | Vitis | vinifera | NC_012119.1 |
| Welwitschiales | Welwitschiaceae | Welwitschia | mirabilis | NC_029130.1 |
