## Additional file 2 for "Mitochondrial genomes of two parasitic *Cuscuta* species lack clear evidence of horizontal gene transfer and retain unusually fragmented *ccmF_C_* genes"

### Additional file 2: supplementary Figs. S1–S5, Tables S3 & S4

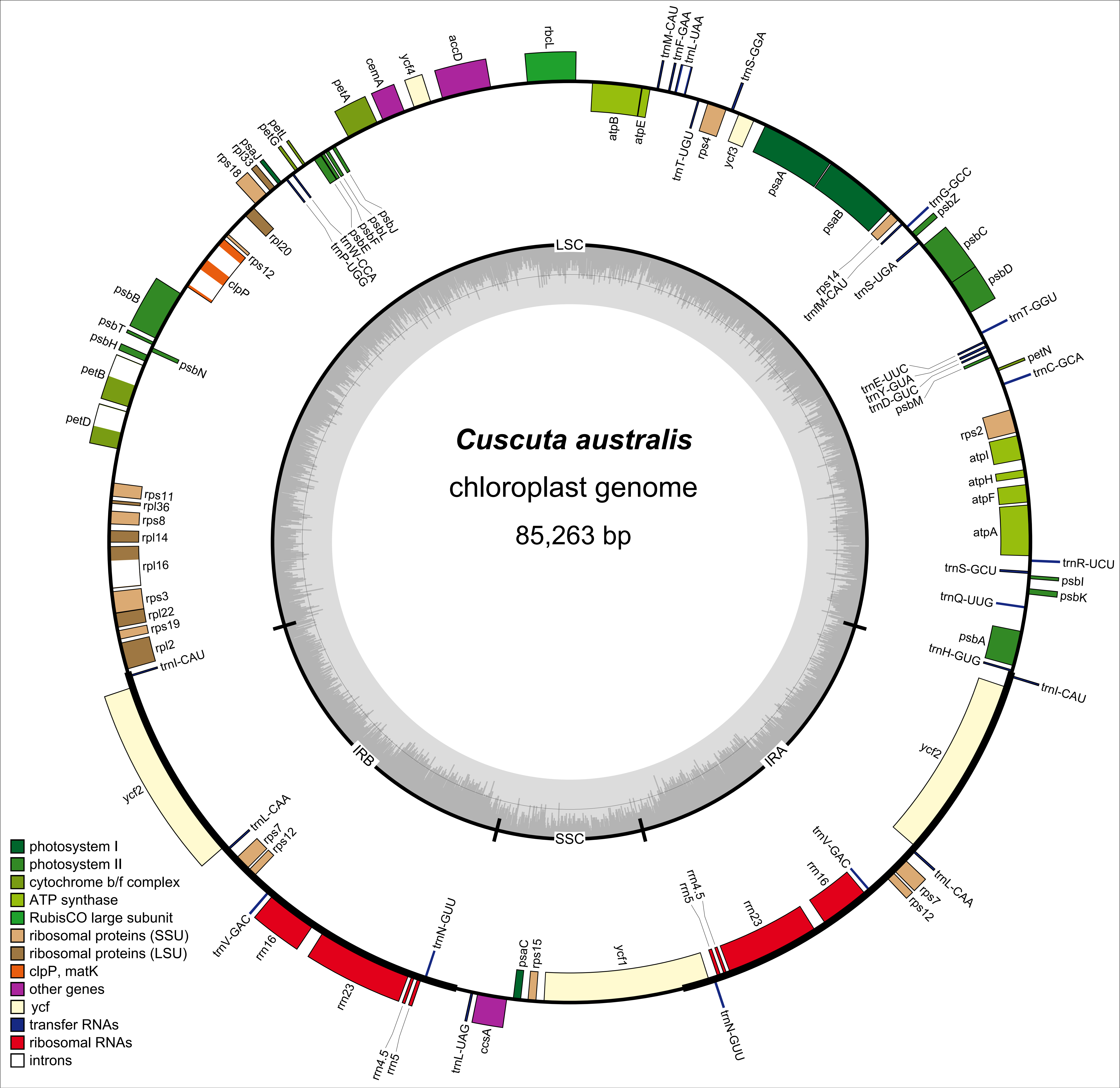
Cuscuta australis
chloroplast genome
85,263 bp
LSC
IRA
IRB
SSC
rbcL
trnM-CAU
trnF-GAA
trnL-UAA
trnS-GGA
atpB
atpE
trnT-UGU
rps4
ycf3
psaA
psaB
rps14
trnfM-CAU
trnG-GCC
psbZ
trnS-UGA
psbC
psbD
trnT-GGU
petN
trnC-GCA
trnE-UUC
trnY-GUA
trnD-GUC
psbM
rps2
atpI
atpH
atpF
atpA
trnR-UCU
trnS-GCU
psbI
psbK
trnQ-UUG
psbA
trnH-GUG
trnI-CAU
ycf2
trnL-CAA
rps7
rps12
trnV-GAC
rrn16
rrn23
rrn4.5
rrn5
trnN-GUU
ycf1
rps15
psaC
ccsA
trnL-UAG
trnN-GUU
rrn23
rrn4.5
rrn5
rrn16
trnV-GAC
rps12
rps7
trnL-CAA
ycf2
trnI-CAU
rpl2
rps19
rpl22
rps3
rpl16
rpl14
rps8
rpl36
rps11
petD
petB
psbH
psbN
psbT
psbB
clpP
rps12
rpl20
rps18
rpl33
psaJ
petG
petL
trnP-UGG
trnW-CCA
psbE
psbF
psbL
psbJ
petA
cemA
ycf4
accD
photosystem I
photosystem II
cytochrome b/f complex
ATP synthase
RubisCO large subunit
ribosomal proteins (SSU)
ribosomal proteins (LSU)
clpP, matK
other genes
ycf
transfer RNAs
ribosomal RNAs
introns

Fig. S1 (previous page). Chloroplast genome of *Cuscuta australis*. Transcription is counterclockwise around the outside of the circle and clockwise inside. The grey inner graph is GC content. Abbreviations: LSC = large single copy region, SSC = small single copy region, IRA/B = inverted repeat A/B.

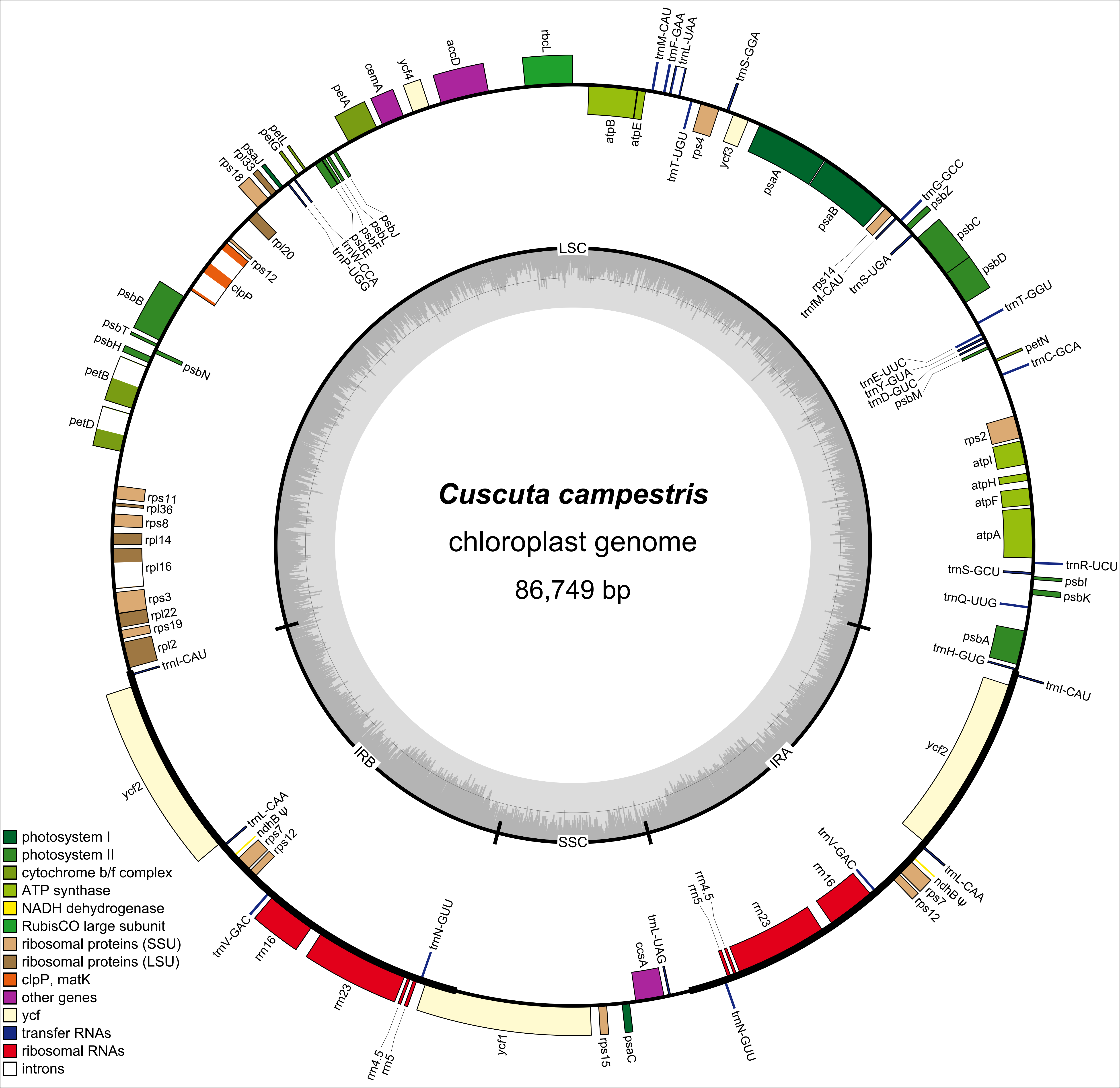

Cuscuta campestris
chloroplast genome
86,749 bp
LSC
IRA
SSC
IRB
rbcL
accD
ycf4
cemA
petA
petL
petG
psaJ
rpl33
rps18
rpl20
rps12
clpP
psbB
psbT
psbH
psbN
petB
petD
psbJ
psbL
psbF
psbE
trnW-CCA
trnP-UGG
trnM-CAU
trnF-GAA
trnL-UAA
trnS-GGA
atpB
atpE
trnT-UGU
rps4
ycf3
psaA
psaB
trnG-GCC
psbZ
psbC
psbD
rps14
trnfM-CAU
trnS-UGA
trnT-GGU
petN
trnC-GCA
trnE-UUC
trnY-GUA
trnD-GUC
psbM
rps2
atpI
atpH
atpF
atpA
trnR-UCU
trnS-GCU
psbI
psbK
trnQ-UUG
psbA
trnH-GUG
trnI-CAU
ycf2
rps11
rpl36
rps8
rpl14
rpl16
rps3
rpl22
rps19
rpl2
trnL-CAA
ndhB ψ
rps7
trnV-GAC
rrn16
rrn23
rrn4.5
rrn5
trnN-GUU
ycf1
rps15
psaC
ccsA
trnL-UAG
photosystem I
photosystem II
cytochrome b/f complex
ATP synthase
NADH dehydrogenase
RubisCO large subunit
ribosomal proteins (SSU)
ribosomal proteins (LSU)
clpP, matK
other genes
ycf
transfer RNAs
ribosomal RNAs
introns

Fig. S2 (previous page). Chloroplast genome of *Cuscuta campestris*. Transcription is counterclockwise around the outside of the circle and clockwise inside. The grey inner graph is GC content. Abbreviations: LSC = large single copy region, SSC = small single copy region, IRA/B = inverted repeat A/B.

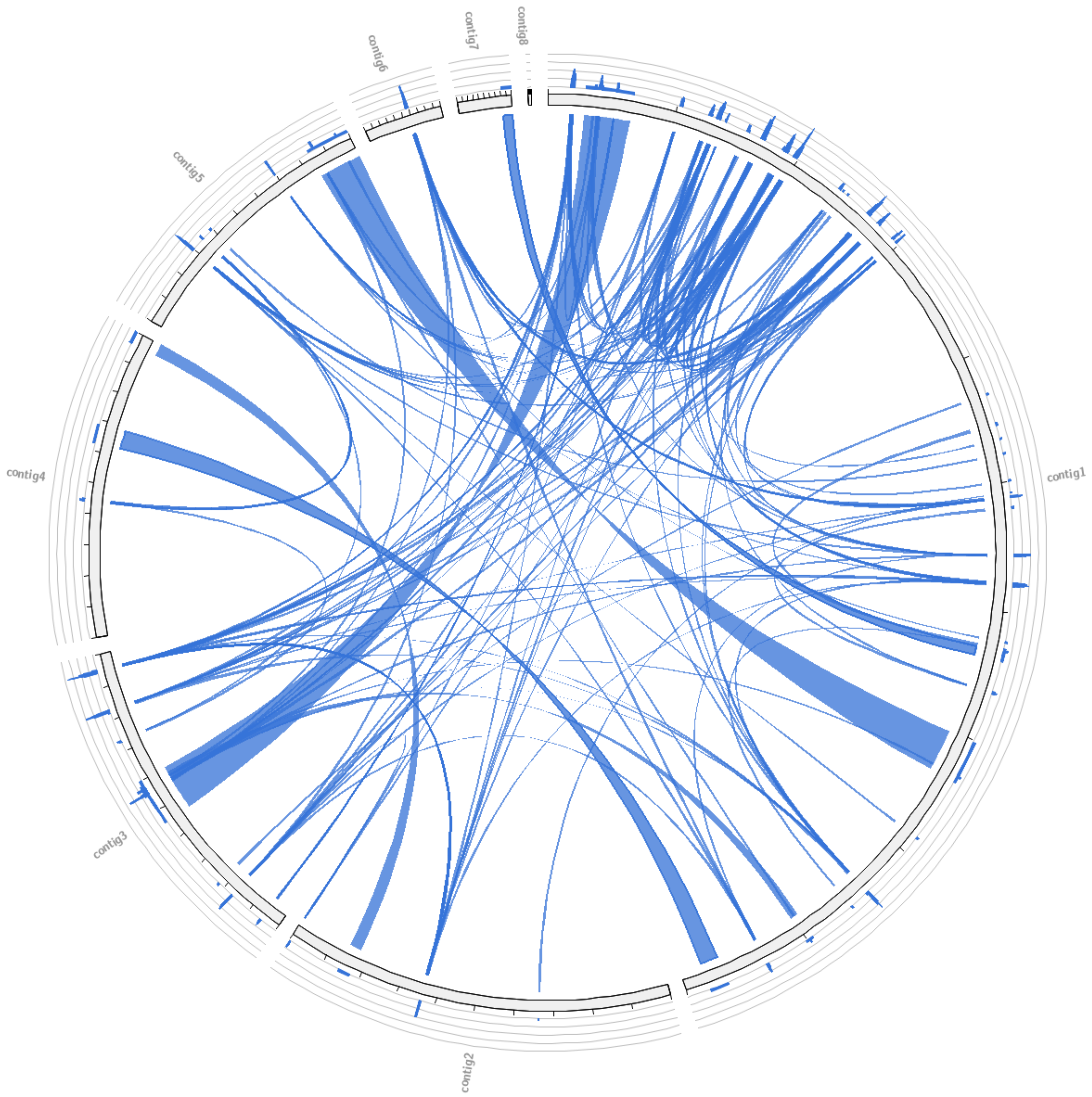
contig8
contig7
contig6
contig5
contig4
contig3
contig2
contig1

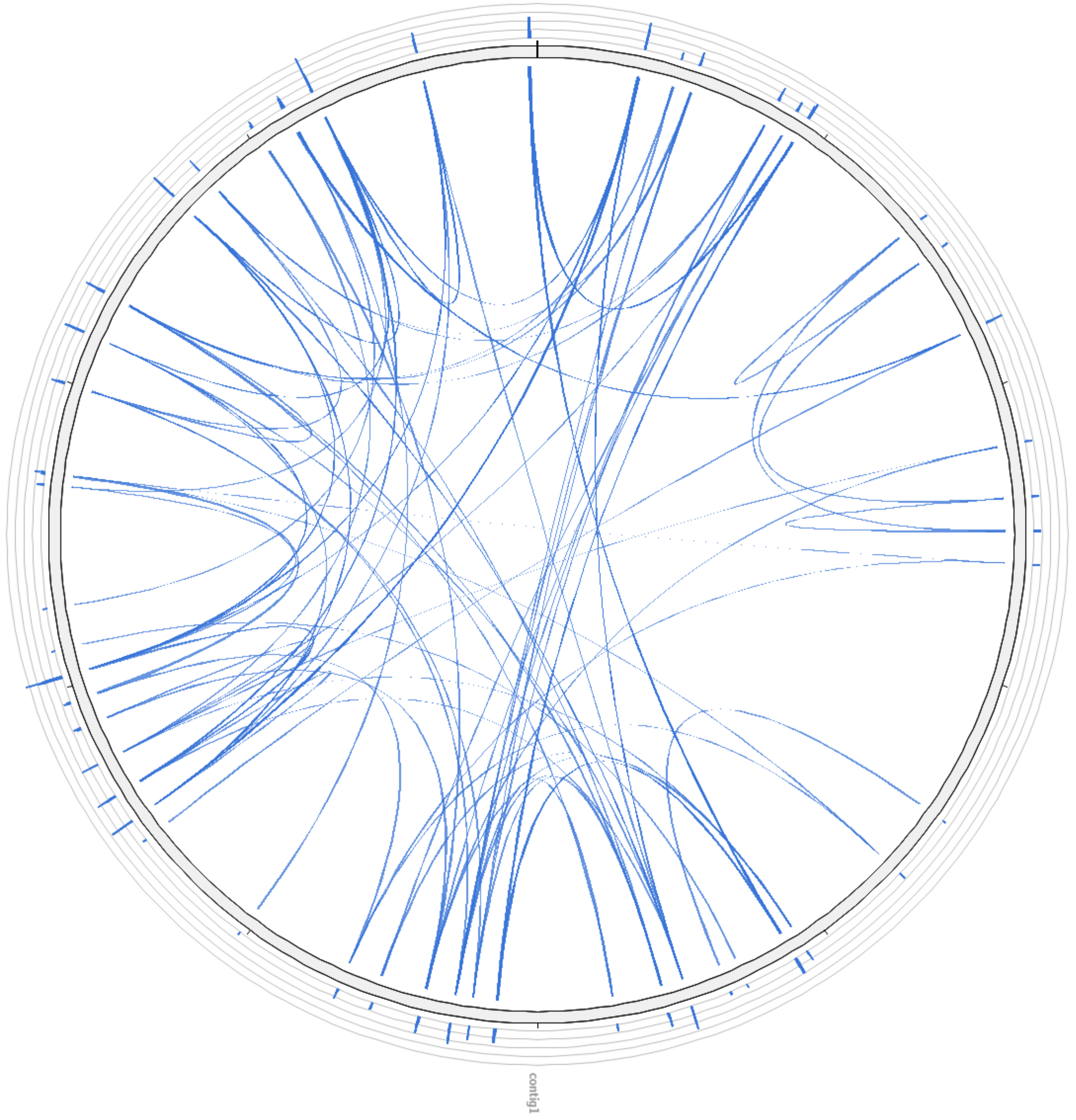
contig1

Fig. S3 (previous page). Dispersed repeats in the mitogenomes of *Cuscuta australis* (left) and *C. campestris* (right). Generated with Circoletto (http://tools.bat.infspire.org/circoletto/).

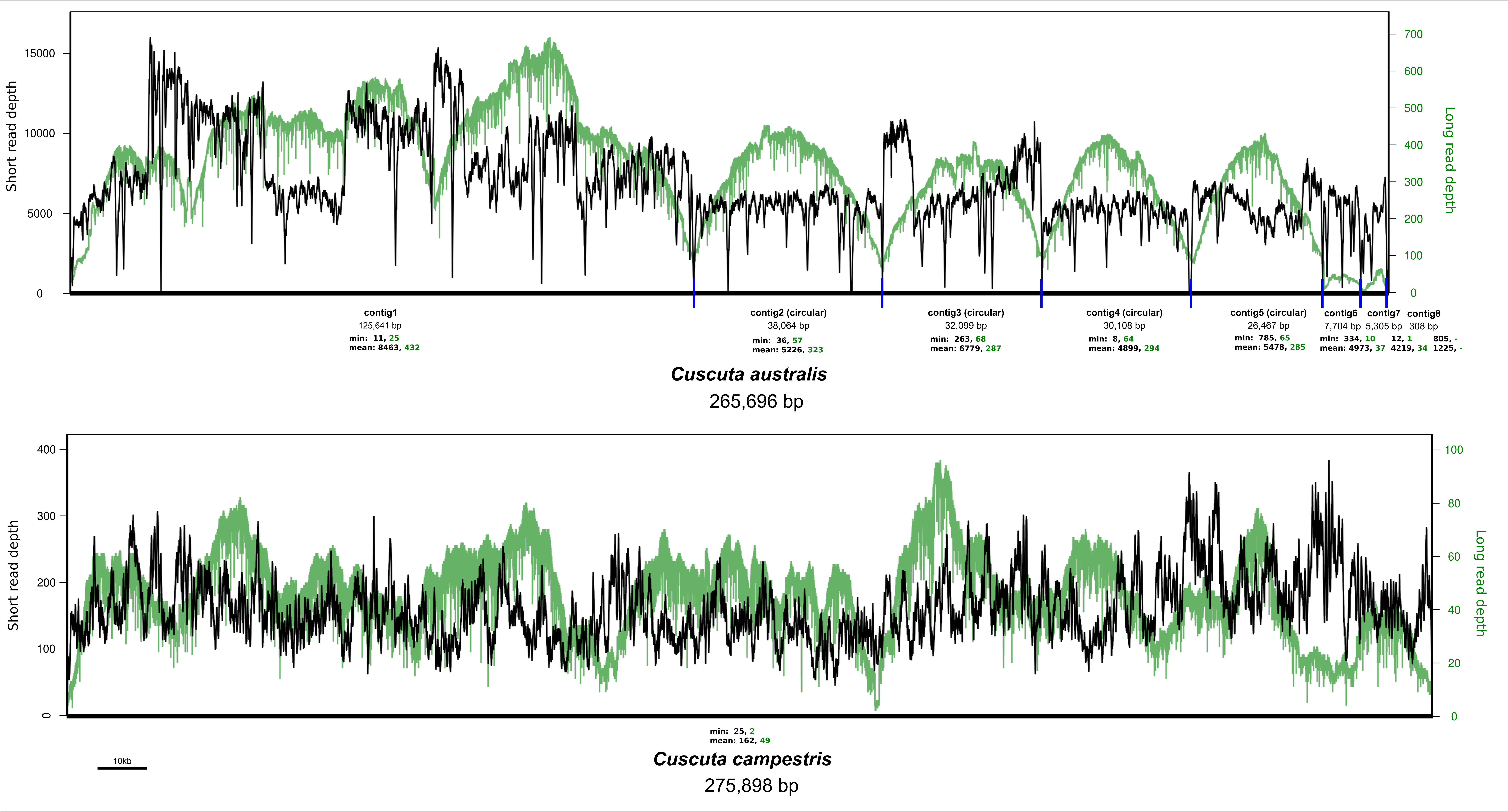

Short read depth
Long read depth
15000
10000
5000
0
700
600
500
400
300
200
100
contig1
125,641 bp
min: 11, 25
mean: 8463, 432
contig2 (circular)
38,064 bp
min: 36, 57
mean: 5226, 323
contig3 (circular)
32,099 bp
min: 263, 68
mean: 6779, 287
contig4 (circular)
30,108 bp
min: 8, 64
mean: 4899, 294
contig5 (circular)
26,467 bp
min: 785, 65
mean: 5478, 285
contig6
7,704 bp
min: 334, 10
mean: 4973, 37
contig7
5,305 bp
12, 1
4219, 34
contig8
308 bp
805, -
1225, -
Cuscuta australis
265,696 bp
400
300
200
100
80
60
40
20
min: 25, 2
mean: 162, 49
10kb
Cuscuta campestris
275,898 bp

Fig. S4 (previous page). Read depth for mitogenome assemblies of *Cuscuta australis* (top) and *C. campestris* (bottom). Short read depth is in black while long read depth is in green. Breakpoints between contigs in *C. australis* are blue lines. Minimum and mean read depths are indicated, with colours of text matching respective read depth.

**Table S3. Repeats recombination analysis for *Cuscuta australis*. The analysis is based on non-overlapping repeats >500 bp and >90% identity (blastn evalue 1e-10), and mapping long reads to those repeats.**

| Repeat | Locus 1 contig | Locus 1 start | Locus 1 end | Locus 2 contig | Locus 2 start | Locus 2 end | Strand | Length (bp) | identity (%) | Reads spanning | # incongruous | % recombined | Note |
|---|---|---|---|---|---|---|---|---|---|---|---|---|---|
| 1 | 3 | 14672 | 19276 | 1 | 3688 | 8292 | - | 4605 | 100.00 | 50 | 14 | 28 | |
| 2 | 5 | 22248 | 26467 | 1 | 88197 | 92416 | - | 4220 | 100.00 | 49 | 16 | 33 | |
| 3 | 4 | 18771 | 20608 | 1 | 121593 | 123430 | - | 1838 | 100.00 | 47 | 16 | 34 | |
| 4 | 4 | 28923 | 30108 | 2 | 31462 | 32647 | - | 1186 | 100.00 | 50 | 11 | 22 | |
| 5 | 7 | 4350 | 5305 | 1 | 79142 | 80097 | - | 956 | 100.00 | 50 | N/A | | hit is at end of contig (alternative path) |
| 6 | 1 | 23546 | 24337 | 1 | 72459 | 73278 | + | 792 | 96.59 | 48 | 5 | 10 | length locus 2 = 820 |
| 7 | 1 | 4229 | 4979 | 1 | 112453 | 113203 | + | 751 | 100.00 | 46 | 22 | 48 | |
| 8 | 1 | 15179 | 15726 | 1 | 125094 | 125641 | + | 548 | 100.00 | 50 | N/A | | hit is at end of contig (alternative path) |
| | | | | | | | | | | | | **29** | **Average** |

**Table S4. Repeats recombination analysis for *Cuscuta campestris*. The analysis is based on non-overlapping repeats >500 bp and >90% identity (blastn evalue 1e-10), and mapping long reads to those repeats.**

| Repeat | Locus 1 start | Locus 1 end | Locus 2 start | Locus 2 end | Strand | Length (bp) | identity (%) | Reads spanning | # incongruous | % recombined | Note |
|---|---|---|---|---|---|---|---|---|---|---|---|
| 1 | 16591 | 23986 | 214870 | 222265 | + | 7396 | 100.00 | 25 | 10 | 40 | |
| 2 | 37606 | 43001 | 64407 | 69802 | + | 5396 | 100.00 | 40 | 15 | 38 | |
| 3 | 220913 | 225483 | 260669 | 265239 | + | 4571 | 100.00 | 32 | 7 | 22 | third alternative connection |
| 4 | 22634 | 27068 | 111195 | 115629 | + | 4435 | 99.98 | 46 | 20 | 43 | |
| 5 | 108143 | 112547 | 257617 | 262021 | + | 4405 | 99.98 | 29 | 12 | 41 | |
| 6 | 138207 | 142395 | 271638 | 275898 | + | 4189 | 98.31 | 40 | 16 | 40 | length locus 2 = 4261 |
| 7 | 164143 | 167027 | 198327 | 201209 | + | 2885 | 99.83 | 48 | 18 | 38 | length locus 2 = 2883 |
| 8 | 71283 | 73704 | 101616 | 104037 | + | 2422 | 99.96 | 48 | 22 | 46 | |
| 9 | 58370 | 60787 | 246678 | 249126 | + | 2418 | 98.69 | 45 | 21 | 47 | length locus 2 = 2449 |
| 10 | 192601 | 194502 | 253720 | 255621 | + | 1902 | 100.00 | 50 | 23 | 46 | |
| 11 | 81543 | 82685 | 183437 | 184579 | + | 1143 | 100.00 | 50 | 5 | 10 | possible third alternative connection |
| 12 | 129048 | 129774 | 162559 | 163285 | + | 727 | 100.00 | 49 | 16 | 33 | |
| | | | | | | | | | | 37 | Average all |
| | | | | | | | | | | **41** | **Average two connections** |

Tables S3 and S4 (previous page). Details of long repeats used for a proxy of recombinational activity in mitogenomes of *Cuscuta australis* and *C. campestris*. In two cases (highlighted) there were more than two alternative orientations inferred (the repeat occurs more than twice and involves connections to multiple contigs/locations in the genome), and in two cases in *C. australis*, the repeats hit the ends of contigs and could not be assessed.

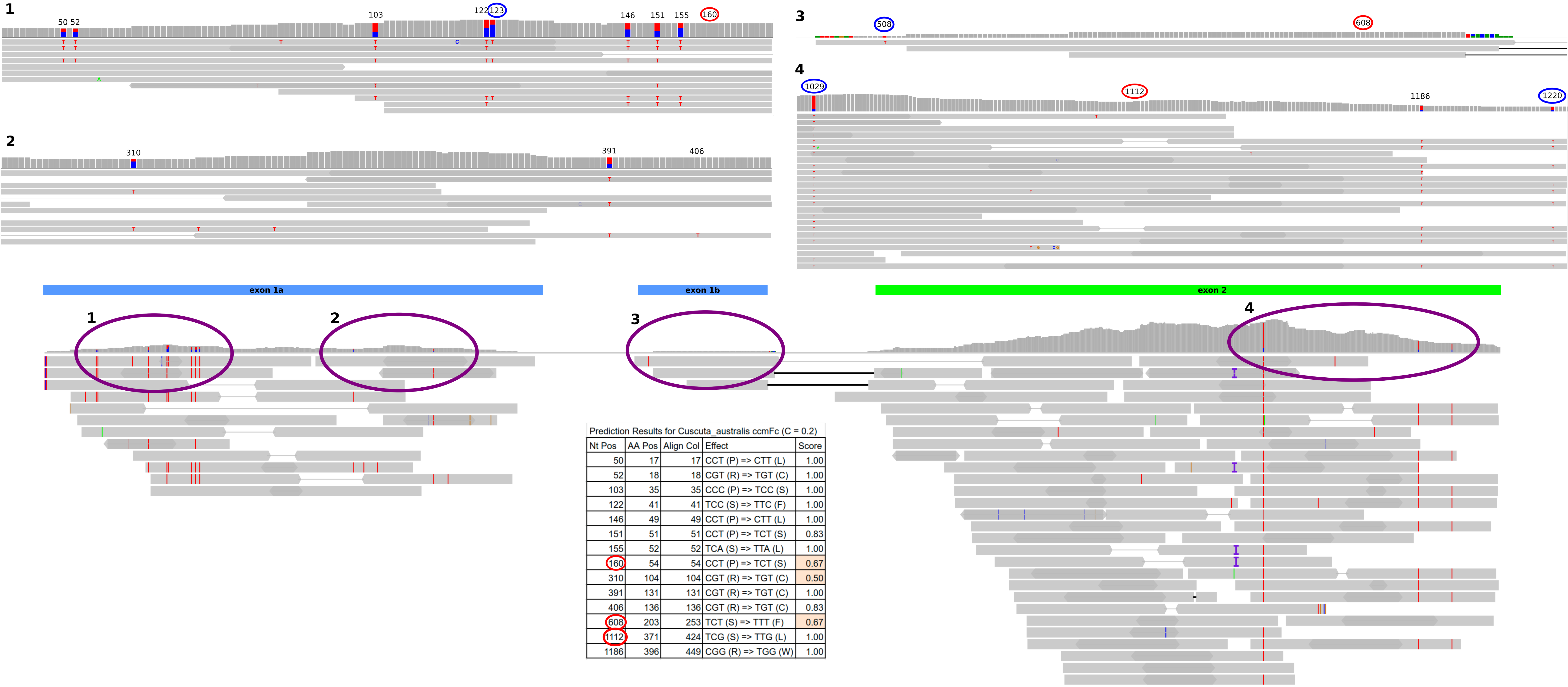


Prediction Results for Cuscuta_australis ccmFc (C = 0.2)

| Nt Pos | AA Pos | Align Col | Effect | Score |
|---|---|---|---|---|
| 50 | 17 | 17 | CCT (P) => CTT (L) | 1.00 |
| 52 | 18 | 18 | CGT (R) => TGT (C) | 1.00 |
| 103 | 35 | 35 | CCC (P) => TCC (S) | 1.00 |
| 122 | 41 | 41 | TCC (S) => TTC (F) | 1.00 |
| 146 | 49 | 49 | CCT (P) => CTT (L) | 1.00 |
| 151 | 51 | 51 | CCT (P) => TCT (S) | 0.83 |
| 155 | 52 | 52 | TCA (S) => TTA (L) | 1.00 |
| 160 | 54 | 54 | CCT (P) => TCT (S) | 0.67 |
| 310 | 104 | 104 | CGT (R) => TGT (C) | 0.50 |
| 391 | 131 | 131 | CGT (R) => TGT (C) | 1.00 |
| 406 | 136 | 136 | CGT (R) => TGT (C) | 0.83 |
| 608 | 203 | 253 | TCT (S) => TTT (F) | 0.67 |
| 1112 | 371 | 424 | TCG (S) => TTG (L) | 1.00 |
| 1186 | 396 | 449 | CGG (R) => TGG (W) | 1.00 |

Fig. S5 (previous page). Edited sites in *ccmF*$_C$ from *Cuscuta australis* based on mapped transcriptomic reads and predicted by PREP-MT. Exons are indicated above the mapped reads, with circles corresponding to panels above showing the mismatches in more detail. Positions along the alignment are numbered, with positions not predicted by PREP-MT circled in blue, and those predicted but not observed by mapping circled in red. The table shows details of the results from PREP-MT.
