## Additional file 3 for "Mitochondrial genomes of two parasitic *Cuscuta* species lack clear evidence of horizontal gene transfer and retain unusually fragmented *ccmF_C_* genes"

### Additional file 3: supplementary Figs. S6–S13

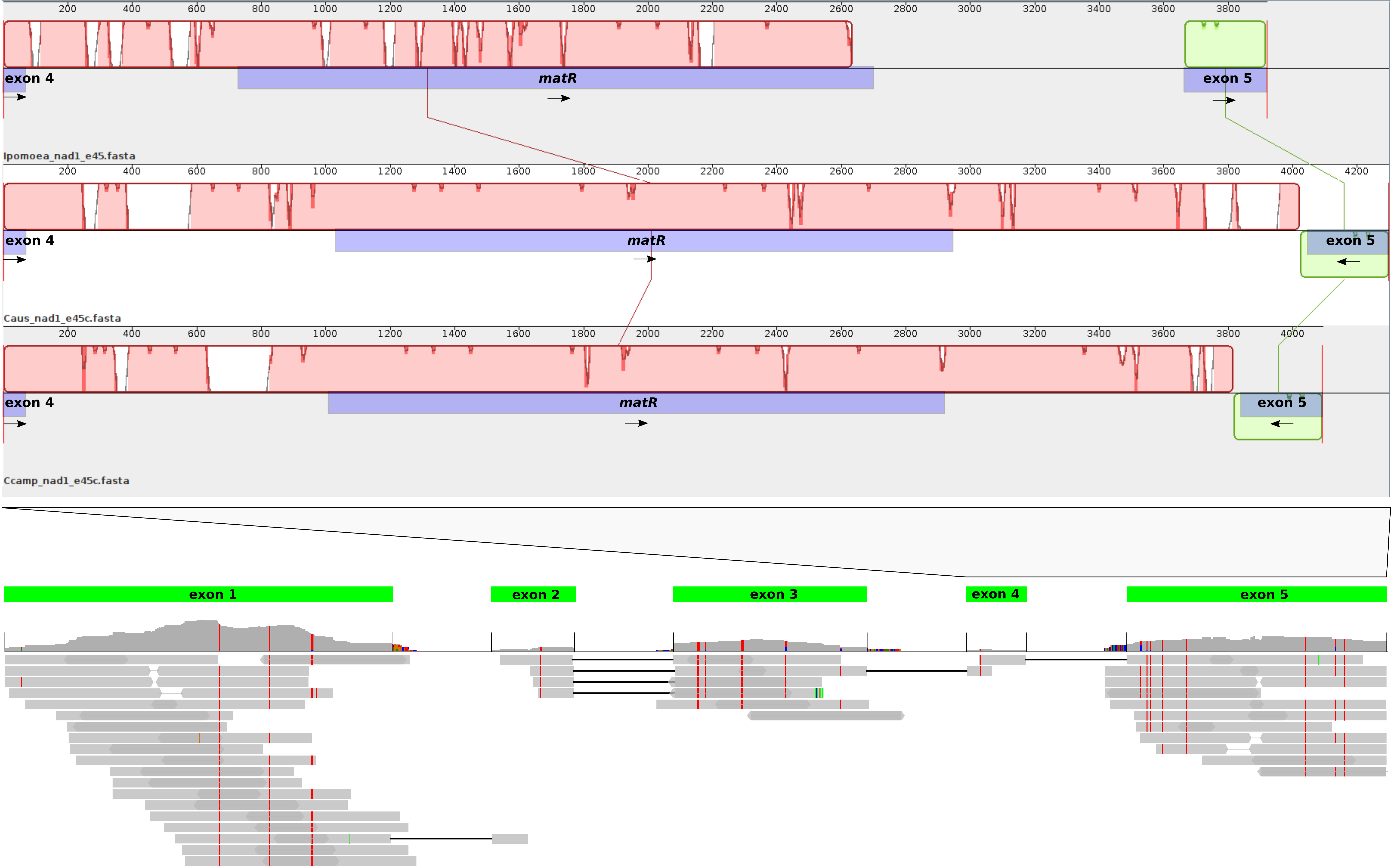
Ipomoea_nad1_e45.fasta
Caus_nad1_e45c.fasta
Ccamp_nad1_e45c.fasta
exon 4
matR
exon 5
exon 1
exon 2
exon 3
exon 4
exon 5

Fig. S6 (previous page). Schematic of the structure of *nad1* in *Cuscuta australis* based on DNA alignment with *C. campestris* and *Ipomoea nil* and alignment of transcriptomic reads. At top, a Mauve alignment of the section from exon4 to exon5 (containing *matR*) is shown for the three species (from top to bottom: *Ipomoea nil*, *Cuscuta australis*, *C. campestris*). Direction of transcription is shown with arrows. Exon structure is shown below with mRNA reads aligned to the DNA sequences separated by 100 Ns. Black lines demarcate the exonic sequences. Coloured lines in aligned reads indicate mismatches, corresponding to putative editing sites in most cases.

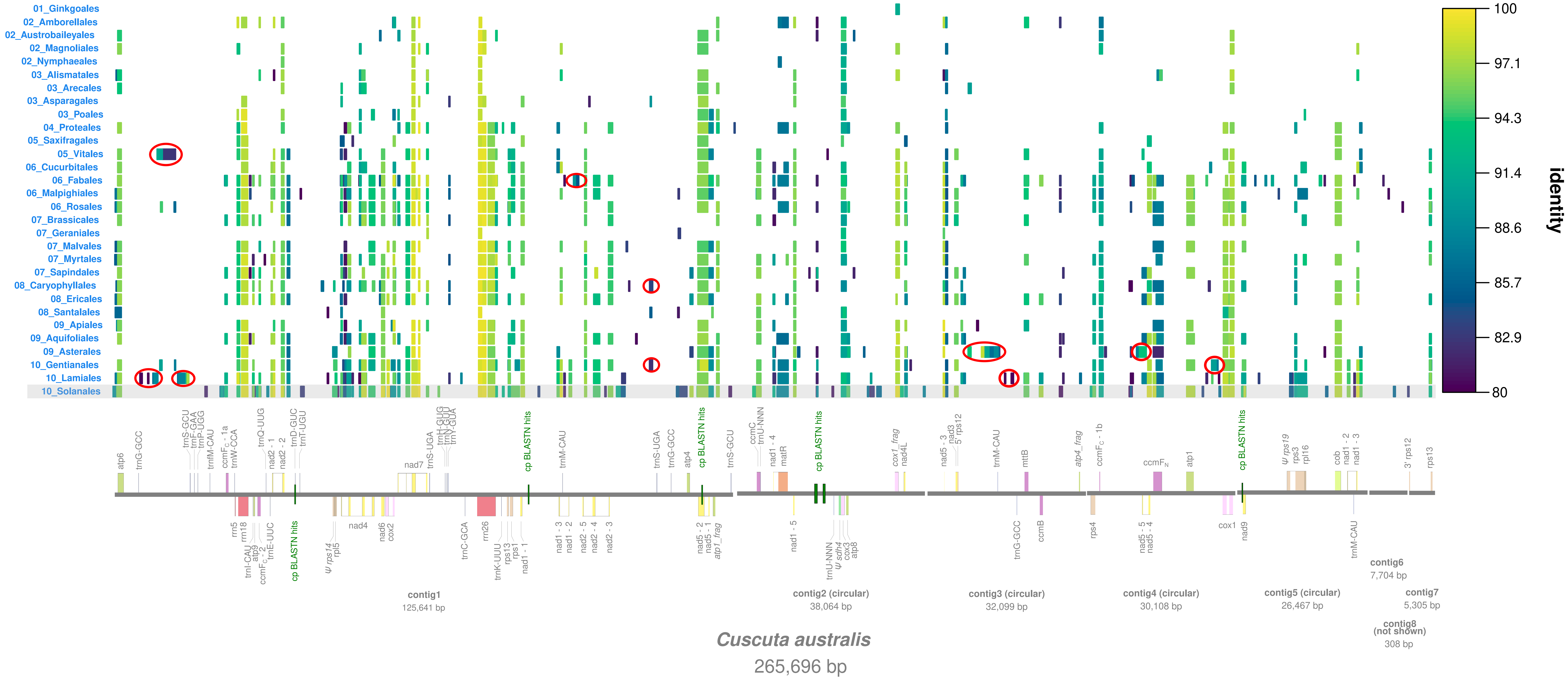
01_Ginkgoales
02_Amborellales
02_Austrobaileyales
02_Magnoliales
02_Nymphaeales
03_Alismatales
03_Arecales
03_Asparagales
03_Poales
04_Proteales
05_Saxifragales
05_Vitales
06_Cucurbitales
06_Fabales
06_Malpighiales
06_Rosales
07_Brassicales
07_Geraniales
07_Malvales
07_Myrtales
07_Sapindales
08_Caryophyllales
08_Ericales
08_Santalales
09_Apiales
09_Aquifoliales
09_Asterales
10_Gentianales
10_Lamiales
10_Solanales
100
97.1
94.3
91.4
88.6
85.7
82.9
80
identity
atp6
trnG-GCC
trnS-GCU
trnF-GAA
trnP-UGG
trnfM-CAU
ccmFC - 1a
trnW-CCA
trnQ-UUG
nad2 - 1
nad2 - 2
trnD-GUC
trnT-UGU
nad7
trnS-UGA
trnH-GUG
trnN-GUU
trnY-GUA
cp BLASTN hits
trnM-CAU
trnS-UGA
trnG-GCC
atp4
cp BLASTN hits
trnS-GCU
ccmC
trnU-NNN
nad1 - 4
matR
cp BLASTN hits
cox1_frag
nad4L
nad5 - 3
nad3
5' rps12
trnM-CAU
mttB
atp4_frag
ccmFC - 1b
ccmFN
atp1
cp BLASTN hits
ψ rps19
rps3
rpl16
cob
nad1 - 2
nad1 - 3
3' rps12
rps13
rrn5
rrn18
trnI-CAU
atp9
ccmFC - 2
trnE-UUC
cp BLASTN hits
ψ rps14
rpl5
nad4
nad6
cox2
trnC-GCA
rrn26
trnK-UUU
rps13
rps1
nad1 - 1
nad1 - 3
nad1 - 2
nad2 - 5
nad2 - 4
nad2 - 3
nad5 - 2
nad5 - 1
atp1_frag
nad1 - 5
trnU-NNN
ψ sdh4
cox3
atp8
trnG-GCC
ccmB
rps4
nad5 - 5
nad5 - 4
cox1
nad9
trnM-CAU
contig1
125,641 bp
contig2 (circular)
38,064 bp
contig3 (circular)
32,099 bp
contig4 (circular)
30,108 bp
contig5 (circular)
26,467 bp
contig6
7,704 bp
contig7
5,305 bp
contig8
(not shown)
308 bp
Cuscuta australis
265,696 bp

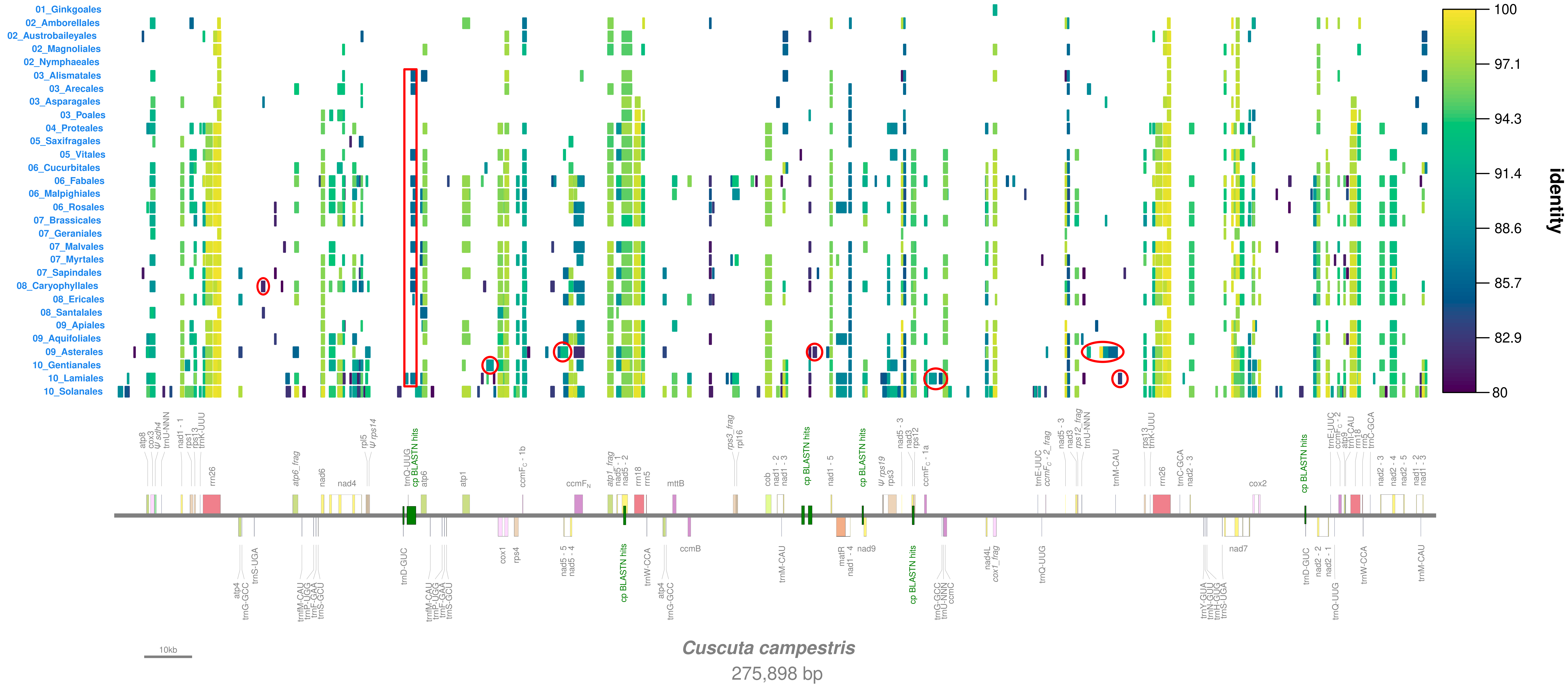

01_Ginkgoales
02_Amborellales
02_Austrobaileyales
02_Magnoliales
02_Nymphaeales
03_Alismatales
03_Arecales
03_Asparagales
03_Poales
04_Proteales
05_Saxifragales
05_Vitales
06_Cucurbitales
06_Fabales
06_Malpighiales
06_Rosales
07_Brassicales
07_Geraniales
07_Malvales
07_Myrtales
07_Sapindales
08_Caryophyllales
08_Ericales
08_Santalales
09_Apiales
09_Aquifoliales
09_Asterales
10_Gentianales
10_Lamiales
10_Solanales
100
97.1
94.3
91.4
88.6
85.7
82.9
80
identity
rrn26
nad4
atp6
atp1
cox1
rps4
ccmB
mttB
nad7
cox2
matR
nad9
cp BLASTN hits
10kb
Cuscuta campestris
275,898 bp

Figs. S7 and S8 (previous two pages). BLASTN hits for reference mitogenomes and plastomes plotted along the mitogenomes of *Cuscuta australis* and *C. campestris*. Plastome hits are shown in green along the linear representations of the mitogenomes, while hits to reference mitogenomes are shown by order above. The percentage identity of the mitogenomic hits is coloured based on the legend at the right. Solanales is shaded, as the order to which *Cuscuta* belongs. Hits >500 bp in regions not hit by Solanales are circled in red as possible HGT candidates.

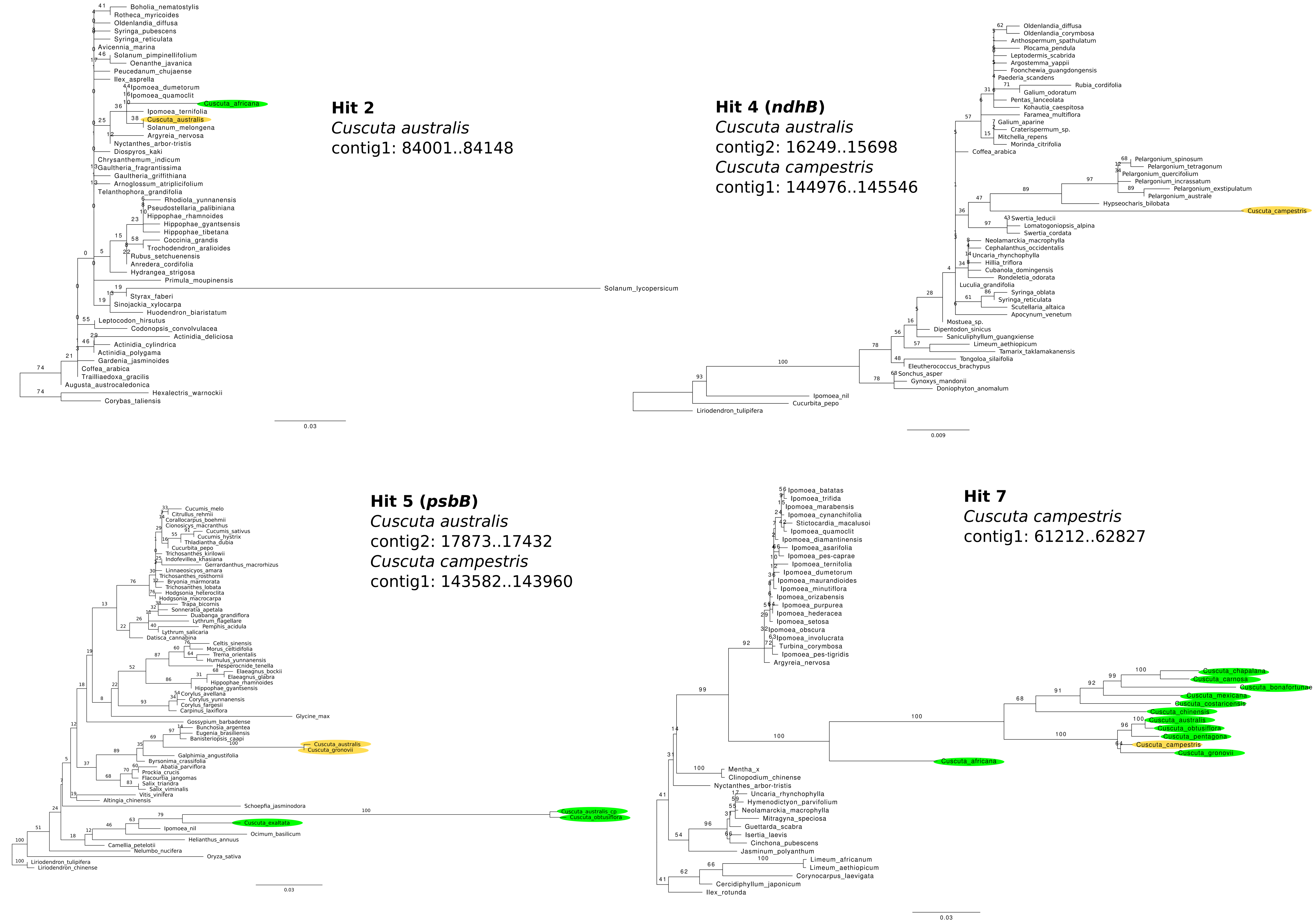
Hit 2
Cuscuta australis
contig1: 84001..84148
Boholia_nematostylis
Rotheca_myricoides
Oldenlandia_diffusa
Syringa_pubescens
Syringa_reticulata
Avicennia_marina
Solanum_pimpinellifolium
Oenanthe_javanica
Peucedanum_chujaense
Ilex_asprella
Ipomoea_dumetorum
Ipomoea_quamoclit
Cuscuta_africana
Ipomoea_ternifolia
Cuscuta_australis
Solanum_melongena
Argyreia_nervosa
Nyctanthes_arbor-tristis
Diospyros_kaki
Chrysanthemum_indicum
Gaultheria_fragrantissima
Gaultheria_griffithiana
Arnoglossum_atriplicifolium
Telanthophora_grandifolia
Rhodiola_yunnanensis
Pseudostellaria_palibiniana
Hippophae_rhamnoides
Hippophae_gyantsensis
Hippophae_tibetana
Coccinia_grandis
Trochodendron_aralioides
Rubus_setchuenensis
Anredera_cordifolia
Hydrangea_strigosa
Primula_moupinensis
Solanum_lycopersicum
Styrax_faberi
Sinojackia_xylocarpa
Huodendron_biaristatum
Leptocodon_hirsutus
Codonopsis_convolvulacea
Actinidia_deliciosa
Actinidia_cylindrica
Actinidia_polygama
Gardenia_jasminoides
Coffea_arabica
Trailliaedoxa_gracilis
Augusta_austrocaledonica
Hexalectris_warnockii
Corybas_taliensis
Hit 4 (ndhB)
Cuscuta australis
contig2: 16249..15698
Cuscuta campestris
contig1: 144976..145546
Oldenlandia_diffusa
Oldenlandia_corymbosa
Anthospermum_spathulatum
Plocama_pendula
Leptodermis_scabrida
Argostemma_yappii
Foonchewia_guangdongensis
Paederia_scandens
Rubia_cordifolia
Galium_odoratum
Pentas_lanceolata
Kohautia_caespitosa
Faramea_multiflora
Galium_aparine
Craterispermum_sp.
Mitchella_repens
Morinda_citrifolia
Coffea_arabica
Pelargonium_spinosum
Pelargonium_tetragonum
Pelargonium_quercifolium
Pelargonium_incrassatum
Pelargonium_exstipulatum
Pelargonium_australe
Hypseocharis_bilobata
Cuscuta_campestris
Swertia_leducii
Lomatogoniopsis_alpina
Swertia_cordata
Neolamarckia_macrophylla
Cephalanthus_occidentalis
Uncaria_rhynchophylla
Hillia_triflora
Cubanola_domingensis
Rondeletia_odorata
Luculia_grandifolia
Syringa_oblata
Syringa_reticulata
Scutellaria_altaica
Apocynum_venetum
Mostuea_sp.
Dipentodon_sinicus
Saniculiphyllum_guangxiense
Limeum_aethiopicum
Tamarix_taklamakanensis
Tongoloa_silaifolia
Eleutherococcus_brachypus
Sonchus_asper
Gynoxys_mandonii
Doniophyton_anomalum
Ipomoea_nil
Cucurbita_pepo
Liriodendron_tulipifera
Hit 5 (psbB)
Cuscuta australis
contig2: 17873..17432
Cuscuta campestris
contig1: 143582..143960
Cucumis_melo
Citrullus_rehmii
Corallocarpus_boehmii
Cionosicys_macranthus
Cucumis_sativus
Cucumis_hystrix
Thladiantha_dubia
Cucurbita_pepo
Trichosanthes_kirilowii
Indofevillea_khasiana
Gerrardanthus_macrorhizus
Linnaeosicyos_amara
Trichosanthes_rosthornii
Bryonia_marmorata
Trichosanthes_lobata
Hodgsonia_heteroclita
Hodgsonia_macrocarpa
Trapa_bicornis
Sonneratia_apetala
Duabanga_grandiflora
Lythrum_flagellare
Pemphis_acidula
Lythrum_salicaria
Datisca_cannabina
Celtis_sinensis
Morus_celtidifolia
Trema_orientalis
Humulus_yunnanensis
Hesperocnide_tenella
Elaeagnus_bockii
Elaeagnus_glabra
Hippophae_rhamnoides
Hippophae_gyantsensis
Corylus_avellana
Corylus_yunnanensis
Corylus_fargesii
Carpinus_laxiflora
Glycine_max
Gossypium_barbadense
Bunchosia_argentea
Eugenia_brasiliensis
Banisteriopsis_caapi
Cuscuta_australis
Cuscuta_gronovii
Galphimia_angustifolia
Byrsonima_crassifolia
Abatia_parviflora
Prockia_crucis
Flacourtia_jangomas
Salix_triandra
Salix_viminalis
Vitis_vinifera
Altingia_chinensis
Schoepfia_jasminodora
Cuscuta_australis_cp
Cuscuta_obtusiflora
Cuscuta_exaltata
Ipomoea_nil
Ocimum_basilicum
Helianthus_annuus
Camellia_petelotii
Nelumbo_nucifera
Oryza_sativa
Liriodendron_tulipifera
Liriodendron_chinense
Hit 7
Cuscuta campestris
contig1: 61212..62827
Ipomoea_batatas
Ipomoea_trifida
Ipomoea_marabensis
Ipomoea_cynanchifolia
Stictocardia_macalusoi
Ipomoea_quamoclit
Ipomoea_diamantinensis
Ipomoea_asarifolia
Ipomoea_pes-caprae
Ipomoea_ternifolia
Ipomoea_dumetorum
Ipomoea_maurandioides
Ipomoea_minutiflora
Ipomoea_orizabensis
Ipomoea_purpurea
Ipomoea_hederacea
Ipomoea_setosa
Ipomoea_obscura
Ipomoea_involucrata
Turbina_corymbosa
Ipomoea_pes-tigridis
Argyreia_nervosa
Cuscuta_chapalana
Cuscuta_carnosa
Cuscuta_bonafortunae
Cuscuta_mexicana
Cuscuta_costaricensis
Cuscuta_chinensis
Cuscuta_australis
Cuscuta_obtusiflora
Cuscuta_pentagona
Cuscuta_campestris
Cuscuta_gronovii
Cuscuta_africana
Mentha_x
Clinopodium_chinense
Nyctanthes_arbor-tristis
Uncaria_rhynchophylla
Hymenodictyon_parvifolium
Neolamarckia_macrophylla
Mitragyna_speciosa
Guettarda_scabra
Isertia_laevis
Cinchona_pubescens
Jasminum_polyanthum
Limeum_africanum
Limeum_aethiopicum
Corynocarpus_laevigata
Cercidiphyllum_japonicum
Ilex_rotunda

Fig. S9 (previous page). Phylogenetic trees of plastid hits in mitogenomes of *Cuscuta australis* and *C. campestris*, analysed with top BLASTN hits from NCBI. All support values from 1000 rapid bootstrap replicates are shown above branches. Hits correspond to Table 2 and include the location within each mitogenome. *Cuscuta* sequences from plastomes are highlighted in green, while *Cuscuta* mitogenomic sequences are highlighted in yellow. Scale bars indicate inferred nucleotide substitutions per site.

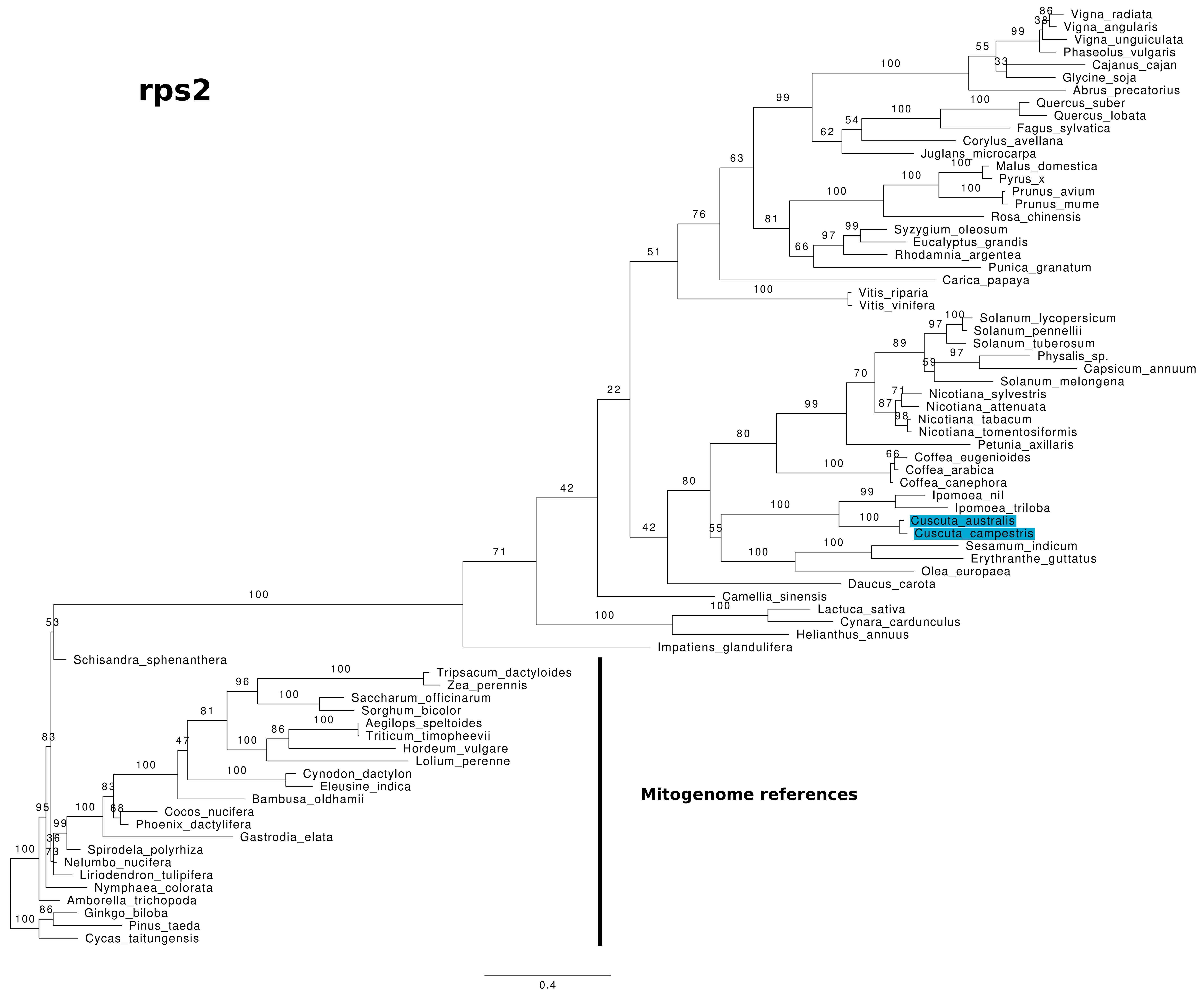
rps2
Vigna_radiata
Vigna_angularis
Vigna_unguiculata
Phaseolus_vulgaris
Cajanus_cajan
Glycine_soja
Abrus_precatorius
Quercus_suber
Quercus_lobata
Fagus_sylvatica
Corylus_avellana
Juglans_microcarpa
Malus_domestica
Pyrus_x
Prunus_avium
Prunus_mume
Rosa_chinensis
Syzygium_oleosum
Eucalyptus_grandis
Rhodamnia_argentea
Punica_granatum
Carica_papaya
Vitis_riparia
Vitis_vinifera
Solanum_lycopersicum
Solanum_pennellii
Solanum_tuberosum
Physalis_sp.
Capsicum_annuum
Solanum_melongena
Nicotiana_sylvestris
Nicotiana_attenuata
Nicotiana_tabacum
Nicotiana_tomentosiformis
Petunia_axillaris
Coffea_eugenioides
Coffea_arabica
Coffea_canephora
Ipomoea_nil
Ipomoea_triloba
Cuscuta_australis
Cuscuta_campestris
Sesamum_indicum
Erythranthe_guttatus
Olea_europaea
Daucus_carota
Camellia_sinensis
Lactuca_sativa
Cynara_cardunculus
Helianthus_annuus
Impatiens_glandulifera
Schisandra_sphenanthera
Tripsacum_dactyloides
Zea_perennis
Saccharum_officinarum
Sorghum_bicolor
Aegilops_speltoides
Triticum_timopheevii
Hordeum_vulgare
Lolium_perenne
Cynodon_dactylon
Eleusine_indica
Bambusa_oldhamii
Cocos_nucifera
Phoenix_dactylifera
Gastrodia_elata
Spirodela_polyrhiza
Nelumbo_nucifera
Liriodendron_tulipifera
Nymphaea_colorata
Amborella_trichopoda
Ginkgo_biloba
Pinus_taeda
Cycas_taitungensis
Mitogenome references
0.4

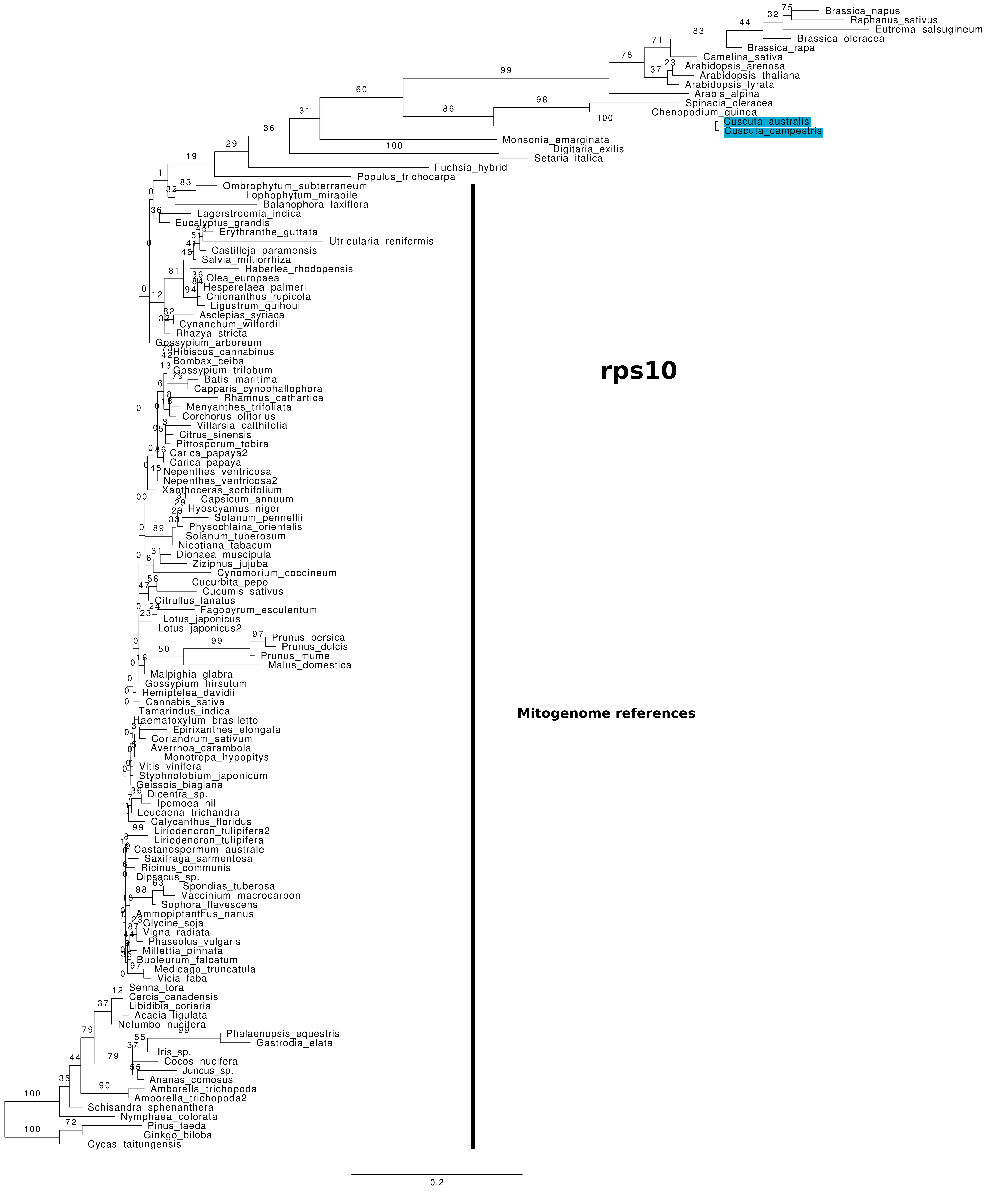
rps10
Brassica_napus
Raphanus_sativus
Eutrema_salsugineum
Brassica_oleracea
Brassica_rapa
Camelina_sativa
Arabidopsis_arenosa
Arabidopsis_thaliana
Arabidopsis_lyrata
Arabis_alpina
Spinacia_oleracea
Chenopodium_quinoa
Cuscuta_australis
Cuscuta_campestris
Monsonia_emarginata
Digitaria_exilis
Setaria_italica
Fuchsia_hybrid
Populus_trichocarpa
Ombrophytum_subterraneum
Lophophytum_mirabile
Balanophora_laxiflora
Lagerstroemia_indica
Eucalyptus_grandis
Erythranthe_guttata
Utricularia_reniformis
Castilleja_paramensis
Salvia_miltiorrhiza
Haberlea_rhodopensis
Olea_europaea
Hesperelaea_palmeri
Chionanthus_rupicola
Ligustrum_quihoui
Asclepias_syriaca
Cynanchum_wilfordii
Rhazya_stricta
Gossypium_arboreum
Hibiscus_cannabinus
Bombax_ceiba
Gossypium_trilobum
Batis_maritima
Capparis_cynophallophora
Rhamnus_cathartica
Menyanthes_trifoliata
Corchorus_olitorius
Villarsia_calthifolia
Citrus_sinensis
Pittosporum_tobira
Carica_papaya2
Carica_papaya
Nepenthes_ventricosa
Nepenthes_ventricosa2
Xanthoceras_sorbifolium
Capsicum_annuum
Hyoscyamus_niger
Solanum_pennellii
Physochlaina_orientalis
Solanum_tuberosum
Nicotiana_tabacum
Dionaea_muscipula
Ziziphus_jujuba
Cynomorium_coccineum
Cucurbita_pepo
Cucumis_sativus
Citrullus_lanatus
Fagopyrum_esculentum
Lotus_japonicus
Lotus_japonicus2
Prunus_persica
Prunus_dulcis
Prunus_mume
Malus_domestica
Malpighia_glabra
Gossypium_hirsutum
Hemiptelea_davidii
Cannabis_sativa
Tamarindus_indica
Haematoxylum_brasiletto
Epirixanthes_elongata
Coriandrum_sativum
Averrhoa_carambola
Monotropa_hypopitys
Vitis_vinifera
Styphnolobium_japonicum
Geissois_biagiana
Dicentra_sp.
Ipomoea_nil
Leucaena_trichandra
Calycanthus_floridus
Liriodendron_tulipifera2
Liriodendron_tulipifera
Castanospermum_australe
Saxifraga_sarmentosa
Ricinus_communis
Dipsacus_sp.
Spondias_tuberosa
Vaccinium_macrocarpon
Sophora_flavescens
Ammopiptanthus_nanus
Glycine_soja
Vigna_radiata
Phaseolus_vulgaris
Millettia_pinnata
Bupleurum_falcatum
Medicago_truncatula
Vicia_faba
Senna_tora
Cercis_canadensis
Libidibia_coriaria
Acacia_ligulata
Nelumbo_nucifera
Phalaenopsis_equestris
Gastrodia_elata
Iris_sp.
Cocos_nucifera
Juncus_sp.
Ananas_comosus
Amborella_trichopoda
Amborella_trichopoda2
Schisandra_sphenanthera
Nymphaea_colorata
Pinus_taeda
Ginkgo_biloba
Cycas_taitungensis
Mitogenome references
0.2

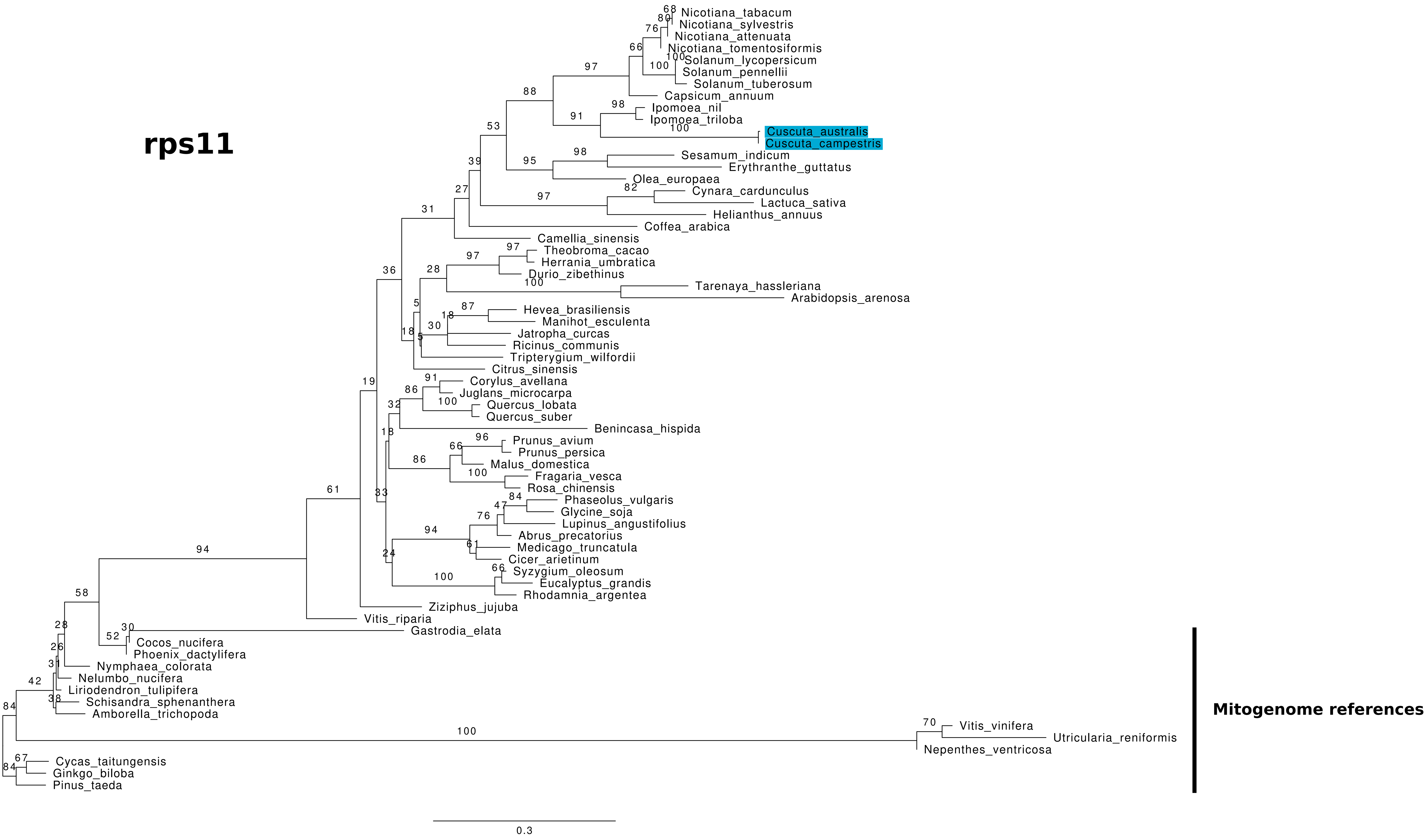
rps11
Nicotiana_tabacum
Nicotiana_sylvestris
Nicotiana_attenuata
Nicotiana_tomentosiformis
Solanum_lycopersicum
Solanum_pennellii
Solanum_tuberosum
Capsicum_annuum
Ipomoea_nil
Ipomoea_triloba
Cuscuta_australis
Cuscuta_campestris
Sesamum_indicum
Erythranthe_guttatus
Olea_europaea
Cynara_cardunculus
Lactuca_sativa
Helianthus_annuus
Coffea_arabica
Camellia_sinensis
Theobroma_cacao
Herrania_umbratica
Durio_zibethinus
Tarenaya_hassleriana
Arabidopsis_arenosa
Hevea_brasiliensis
Manihot_esculenta
Jatropha_curcas
Ricinus_communis
Tripterygium_wilfordii
Citrus_sinensis
Corylus_avellana
Juglans_microcarpa
Quercus_lobata
Quercus_suber
Benincasa_hispida
Prunus_avium
Prunus_persica
Malus_domestica
Fragaria_vesca
Rosa_chinensis
Phaseolus_vulgaris
Glycine_soja
Lupinus_angustifolius
Abrus_precatorius
Medicago_truncatula
Cicer_arietinum
Syzygium_oleosum
Eucalyptus_grandis
Rhodamnia_argentea
Ziziphus_jujuba
Vitis_riparia
Gastrodia_elata
Cocos_nucifera
Phoenix_dactylifera
Nymphaea_colorata
Nelumbo_nucifera
Liriodendron_tulipifera
Schisandra_sphenanthera
Amborella_trichopoda
Vitis_vinifera
Utricularia_reniformis
Nepenthes_ventricosa
Cycas_taitungensis
Ginkgo_biloba
Pinus_taeda
Mitogenome references
0.3

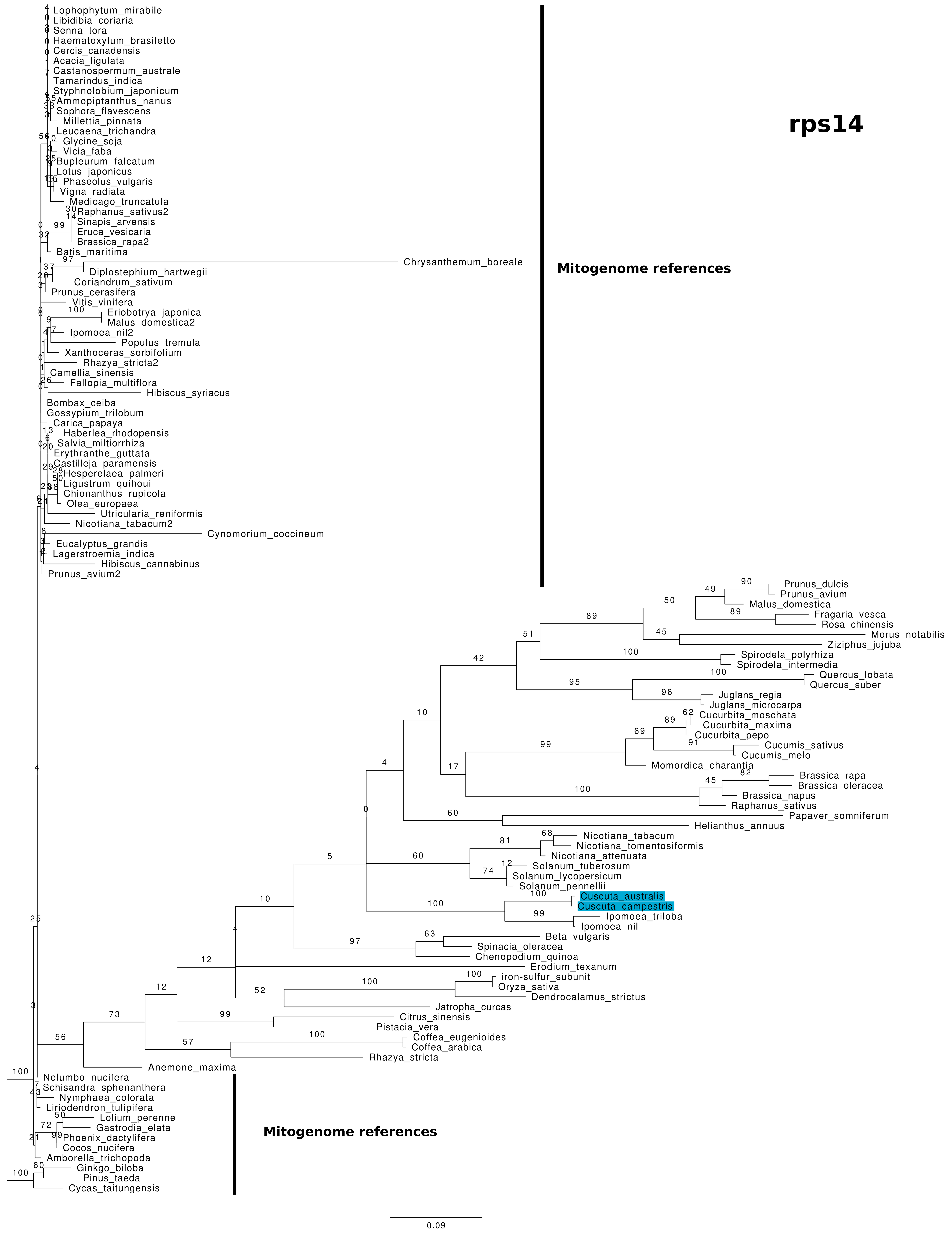
rps14
Lophophytum_mirabile
Libidibia_coriaria
Senna_tora
Haematoxylum_brasiletto
Cercis_canadensis
Acacia_ligulata
Castanospermum_australe
Tamarindus_indica
Styphnolobium_japonicum
Ammopiptanthus_nanus
Sophora_flavescens
Millettia_pinnata
Leucaena_trichandra
Glycine_soja
Vicia_faba
Bupleurum_falcatum
Lotus_japonicus
Phaseolus_vulgaris
Vigna_radiata
Medicago_truncatula
Raphanus_sativus2
Sinapis_arvensis
Eruca_vesicaria
Brassica_rapa2
Batis_maritima
Chrysanthemum_boreale
Diplostephium_hartwegii
Coriandrum_sativum
Prunus_cerasifera
Vitis_vinifera
Eriobotrya_japonica
Malus_domestica2
Ipomoea_nil2
Populus_tremula
Xanthoceras_sorbifolium
Rhazya_stricta2
Camellia_sinensis
Fallopia_multiflora
Hibiscus_syriacus
Bombax_ceiba
Gossypium_trilobum
Carica_papaya
Haberlea_rhodopensis
Salvia_miltiorrhiza
Erythranthe_guttata
Castilleja_paramensis
Hesperelaea_palmeri
Ligustrum_quihoui
Chionanthus_rupicola
Olea_europaea
Utricularia_reniformis
Nicotiana_tabacum2
Cynomorium_coccineum
Eucalyptus_grandis
Lagerstroemia_indica
Hibiscus_cannabinus
Prunus_avium2
Mitogenome references
Prunus_dulcis
Prunus_avium
Malus_domestica
Fragaria_vesca
Rosa_chinensis
Morus_notabilis
Ziziphus_jujuba
Spirodela_polyrhiza
Spirodela_intermedia
Quercus_lobata
Quercus_suber
Juglans_regia
Juglans_microcarpa
Cucurbita_moschata
Cucurbita_maxima
Cucurbita_pepo
Cucumis_sativus
Cucumis_melo
Momordica_charantia
Brassica_rapa
Brassica_oleracea
Brassica_napus
Raphanus_sativus
Papaver_somniferum
Helianthus_annuus
Nicotiana_tabacum
Nicotiana_tomentosiformis
Nicotiana_attenuata
Solanum_tuberosum
Solanum_lycopersicum
Solanum_pennellii
Cuscuta_australis
Cuscuta_campestris
Ipomoea_triloba
Ipomoea_nil
Beta_vulgaris
Spinacia_oleracea
Chenopodium_quinoa
Erodium_texanum
iron-sulfur_subunit
Oryza_sativa
Dendrocalamus_strictus
Jatropha_curcas
Citrus_sinensis
Pistacia_vera
Coffea_eugenioides
Coffea_arabica
Rhazya_stricta
Anemone_maxima
Nelumbo_nucifera
Schisandra_sphenanthera
Nymphaea_colorata
Liriodendron_tulipifera
Lolium_perenne
Gastrodia_elata
Phoenix_dactylifera
Cocos_nucifera
Amborella_trichopoda
Ginkgo_biloba
Pinus_taeda
Cycas_taitungensis
Mitogenome references
0.09

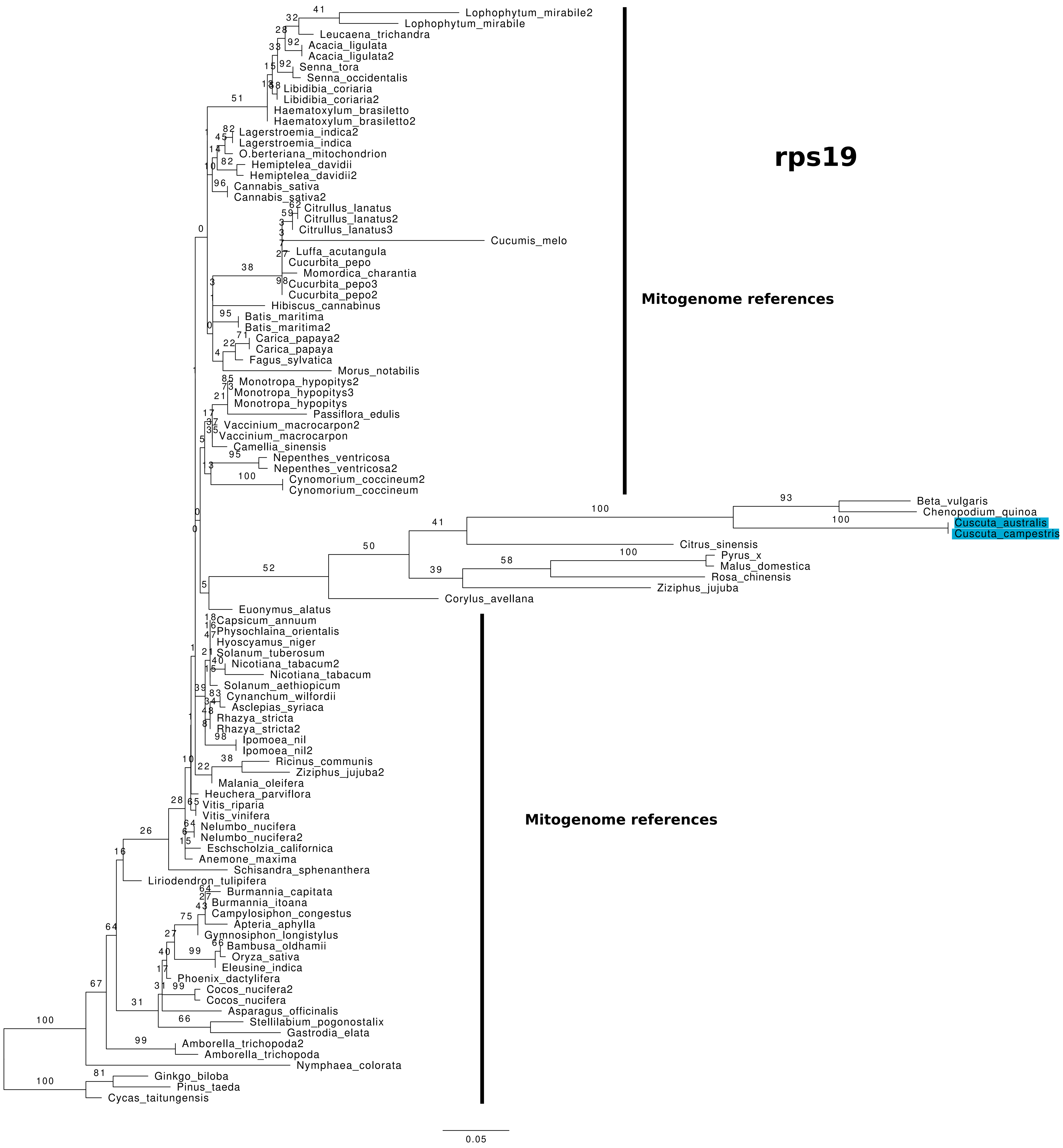
rps19
Mitogenome references
Mitogenome references
Lophophytum_mirabile2
Lophophytum_mirabile
Leucaena_trichandra
Acacia_ligulata
Acacia_ligulata2
Senna_tora
Senna_occidentalis
Libidibia_coriaria
Libidibia_coriaria2
Haematoxylum_brasiletto
Haematoxylum_brasiletto2
Lagerstroemia_indica2
Lagerstroemia_indica
O.berteriana_mitochondrion
Hemiptelea_davidii
Hemiptelea_davidii2
Cannabis_sativa
Cannabis_sativa2
Citrullus_lanatus
Citrullus_lanatus2
Citrullus_lanatus3
Cucumis_melo
Luffa_acutangula
Cucurbita_pepo
Momordica_charantia
Cucurbita_pepo3
Cucurbita_pepo2
Hibiscus_cannabinus
Batis_maritima
Batis_maritima2
Carica_papaya2
Carica_papaya
Fagus_sylvatica
Morus_notabilis
Monotropa_hypopitys2
Monotropa_hypopitys3
Monotropa_hypopitys
Passiflora_edulis
Vaccinium_macrocarpon2
Vaccinium_macrocarpon
Camellia_sinensis
Nepenthes_ventricosa
Nepenthes_ventricosa2
Cynomorium_coccineum2
Cynomorium_coccineum
Beta_vulgaris
Chenopodium_quinoa
Cuscuta_australis
Cuscuta_campestris
Citrus_sinensis
Pyrus_x
Malus_domestica
Rosa_chinensis
Ziziphus_jujuba
Corylus_avellana
Euonymus_alatus
Capsicum_annuum
Physochlaina_orientalis
Hyoscyamus_niger
Solanum_tuberosum
Nicotiana_tabacum2
Nicotiana_tabacum
Solanum_aethiopicum
Cynanchum_wilfordii
Asclepias_syriaca
Rhazya_stricta
Rhazya_stricta2
Ipomoea_nil
Ipomoea_nil2
Ricinus_communis
Ziziphus_jujuba2
Malania_oleifera
Heuchera_parviflora
Vitis_riparia
Vitis_vinifera
Nelumbo_nucifera
Nelumbo_nucifera2
Eschscholzia_californica
Anemone_maxima
Schisandra_sphenanthera
Liriodendron_tulipifera
Burmannia_capitata
Burmannia_itoana
Campylosiphon_congestus
Apteria_aphylla
Gymnosiphon_longistylus
Bambusa_oldhamii
Oryza_sativa
Eleusine_indica
Phoenix_dactylifera
Cocos_nucifera2
Cocos_nucifera
Asparagus_officinalis
Stellilabium_pogonostalix
Gastrodia_elata
Amborella_trichopoda2
Amborella_trichopoda
Nymphaea_colorata
Ginkgo_biloba
Pinus_taeda
Cycas_taitungensis
0.05

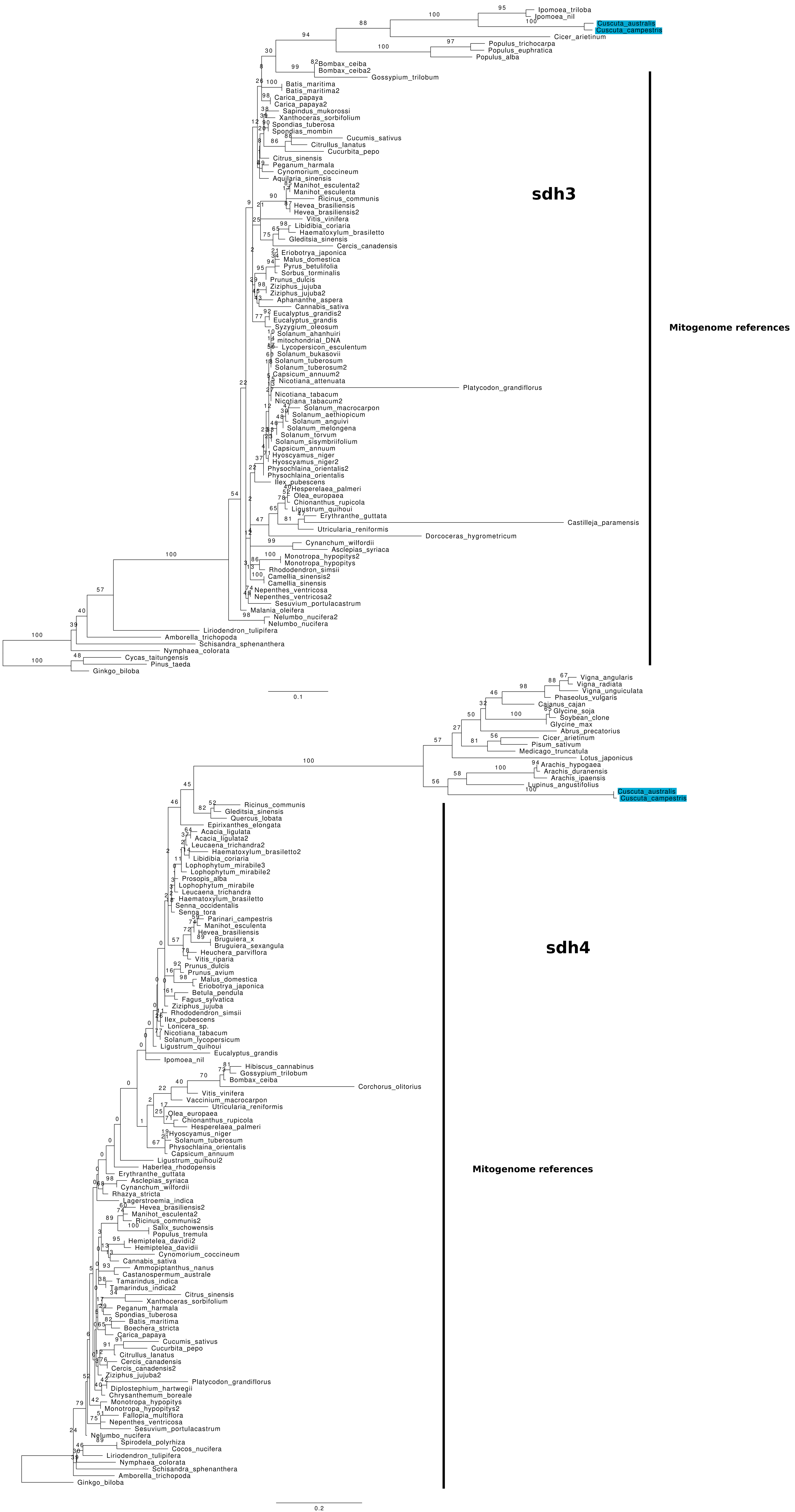

Ipomoea_triloba
Ipomoea_nil
Cuscuta_australis
Cuscuta_campestris
Cicer_arietinum
Populus_trichocarpa
Populus_euphratica
Populus_alba
Bombax_ceiba
Bombax_ceiba2
Gossypium_trilobum
Batis_maritima
Batis_maritima2
Carica_papaya
Carica_papaya2
Sapindus_mukorossi
Xanthoceras_sorbifolium
Spondias_tuberosa
Spondias_mombin
Cucumis_sativus
Citrullus_lanatus
Cucurbita_pepo
Citrus_sinensis
Peganum_harmala
Cynomorium_coccineum
Aquilaria_sinensis
Manihot_esculenta2
Manihot_esculenta
Ricinus_communis
Hevea_brasiliensis
Hevea_brasiliensis2
Vitis_vinifera
Libidibia_coriaria
Haematoxylum_brasiletto
Gleditsia_sinensis
Cercis_canadensis
Eriobotrya_japonica
Malus_domestica
Pyrus_betulifolia
Sorbus_torminalis
Prunus_dulcis
Ziziphus_jujuba
Ziziphus_jujuba2
Aphananthe_aspera
Cannabis_sativa
Eucalyptus_grandis2
Eucalyptus_grandis
Syzygium_oleosum
Solanum_ahanhuiri
mitochondrial_DNA
Lycopersicon_esculentum
Solanum_bukasovii
Solanum_tuberosum
Solanum_tuberosum2
Capsicum_annuum2
Nicotiana_attenuata
Platycodon_grandiflorus
Nicotiana_tabacum
Nicotiana_tabacum2
Solanum_macrocarpon
Solanum_aethiopicum
Solanum_anguivi
Solanum_melongena
Solanum_torvum
Solanum_sisymbriifolium
Capsicum_annuum
Hyoscyamus_niger
Hyoscyamus_niger2
Physochlaina_orientalis2
Physochlaina_orientalis
Ilex_pubescens
Hesperelaea_palmeri
Olea_europaea
Chionanthus_rupicola
Ligustrum_quihoui
Erythranthe_guttata
Castilleja_paramensis
Utricularia_reniformis
Dorcoceras_hygrometricum
Cynanchum_wilfordii
Asclepias_syriaca
Monotropa_hypopitys2
Monotropa_hypopitys
Rhododendron_simsii
Camellia_sinensis2
Camellia_sinensis
Nepenthes_ventricosa
Nepenthes_ventricosa2
Sesuvium_portulacastrum
Malania_oleifera
Nelumbo_nucifera2
Nelumbo_nucifera
Liriodendron_tulipifera
Amborella_trichopoda
Schisandra_sphenanthera
Nymphaea_colorata
Cycas_taitungensis
Pinus_taeda
Ginkgo_biloba
0.1
sdh3
Mitogenome references
Vigna_angularis
Vigna_radiata
Vigna_unguiculata
Phaseolus_vulgaris
Cajanus_cajan
Glycine_soja
Soybean_clone
Glycine_max
Abrus_precatorius
Cicer_arietinum
Pisum_sativum
Medicago_truncatula
Lotus_japonicus
Arachis_hypogaea
Arachis_duranensis
Arachis_ipaensis
Lupinus_angustifolius
Cuscuta_australis
Cuscuta_campestris
Ricinus_communis
Gleditsia_sinensis
Quercus_lobata
Epirixanthes_elongata
Acacia_ligulata
Acacia_ligulata2
Leucaena_trichandra2
Haematoxylum_brasiletto2
Libidibia_coriaria
Lophophytum_mirabile3
Lophophytum_mirabile2
Prosopis_alba
Lophophytum_mirabile
Leucaena_trichandra
Haematoxylum_brasiletto
Senna_occidentalis
Senna_tora
Parinari_campestris
Manihot_esculenta
Hevea_brasiliensis
Bruguiera_x
Bruguiera_sexangula
Heuchera_parviflora
Vitis_riparia
Prunus_dulcis
Prunus_avium
Malus_domestica
Eriobotrya_japonica
Betula_pendula
Fagus_sylvatica
Ziziphus_jujuba
Rhododendron_simsii
Ilex_pubescens
Lonicera_sp.
Nicotiana_tabacum
Solanum_lycopersicum
Ligustrum_quihoui
Eucalyptus_grandis
Ipomoea_nil
Hibiscus_cannabinus
Gossypium_trilobum
Bombax_ceiba
Corchorus_olitorius
Vitis_vinifera
Vaccinium_macrocarpon
Utricularia_reniformis
Olea_europaea
Chionanthus_rupicola
Hesperelaea_palmeri
Hyoscyamus_niger
Solanum_tuberosum
Physochlaina_orientalis
Capsicum_annuum
Ligustrum_quihoui2
Haberlea_rhodopensis
Erythranthe_guttata
Asclepias_syriaca
Cynanchum_wilfordii
Rhazya_stricta
Lagerstroemia_indica
Hevea_brasiliensis2
Manihot_esculenta2
Ricinus_communis2
Salix_suchowensis
Populus_tremula
Hemiptelea_davidii2
Hemiptelea_davidii
Cynomorium_coccineum
Cannabis_sativa
Ammopiptanthus_nanus
Castanospermum_australe
Tamarindus_indica
Tamarindus_indica2
Citrus_sinensis
Xanthoceras_sorbifolium
Peganum_harmala
Spondias_tuberosa
Batis_maritima
Boechera_stricta
Carica_papaya
Cucumis_sativus
Cucurbita_pepo
Citrullus_lanatus
Cercis_canadensis
Cercis_canadensis2
Ziziphus_jujuba2
Platycodon_grandiflorus
Diplostephium_hartwegii
Chrysanthemum_boreale
Monotropa_hypopitys
Monotropa_hypopitys2
Fallopia_multiflora
Nepenthes_ventricosa
Sesuvium_portulacastrum
Nelumbo_nucifera
Spirodela_polyrhiza
Cocos_nucifera
Liriodendron_tulipifera
Nymphaea_colorata
Schisandra_sphenanthera
Amborella_trichopoda
Ginkgo_biloba
0.2
sdh4
Mitogenome references

Figs. S10–13 (previous pages). Phylogenetic trees for nuclear candidate copies of mitochondrial genes in *Cuscuta australis* and *C. campestris* in relation to known mitochondrial copies in other plants and top BLASTN hits from NCBI. Bootstrap support values from 1000 rapid bootstrap replicates are shown above branches. The name of each candidate gene copy is in bold adjacent to its respective tree. Mitogenomic reference sequences (and BLASTN hits to mitogenomes) are indicated. Sequences from *Cuscuta* are highlighted in blue. Scale bars indicate inferred nucleotide substitutions per site.
