## Additional file 4 for "Mitochondrial genomes of two parasitic *Cuscuta* species lack clear evidence of horizontal gene transfer and retain unusually fragmented *ccmF_C_* genes"

### Additional file 4: supplementary Fig. S14

Fig. S14 (subsequent pages). Phylogenetic trees for mitochondrial genes across angiosperms, with particular focus (red) on *Cuscuta* species. Bootstrap support >60% is shown adjacent to nodes. Scale bars indicate inferred nucleotide substitutions per site. Groups are coloured according to the legend below, with the exception of *Cuscuta* (red). The names of the genes are indicated above their respective trees.

- non-angiosperms
- basal + magnoliids
- monocots
- basal eudicots
- rosids
- rosids / fabids
- rosids / malvids
- asterids
- asterids / campanulids
- asterids / lamiids

**atp1**

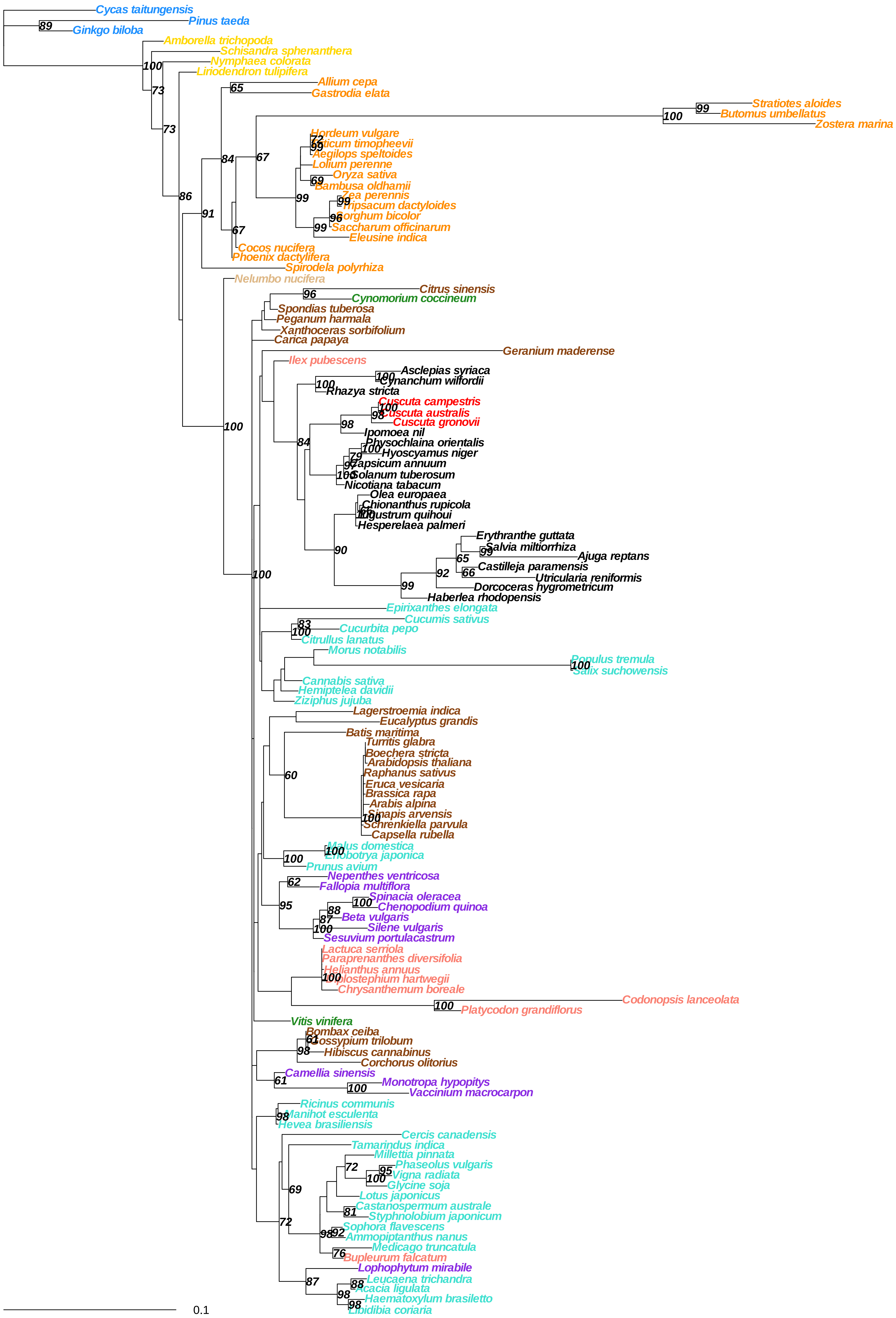

Cycas taitungensis
Pinus taeda
Ginkgo biloba
Amborella trichopoda
Schisandra sphenanthera
Nymphaea colorata
Liriodendron tulipifera
Allium cepa
Gastrodia elata
Stratiotes aloides
Butomus umbellatus
Zostera marina
Hordeum vulgare
Aegilops speltoides
Lolium perenne
Oryza sativa
Bambusa oldhamii
Zea perennis
Tripsacum dactyloides
Sorghum bicolor
Saccharum officinarum
Eleusine indica
Cocos nucifera
Phoenix dactylifera
Spirodela polyrhiza
Nelumbo nucifera
Citrus sinensis
Cynomorium coccineum
Spondias tuberosa
Peganum harmala
Xanthoceras sorbifolium
Carica papaya
Geranium maderense
Ilex pubescens
Asclepias syriaca
Cynanchum wilfordii
Rhazya stricta
Cuscuta campestris
Cuscuta australis
Cuscuta gronovii
Ipomoea nil
Physochlaina orientalis
Hyoscyamus niger
Capsicum annuum
Solanum tuberosum
Nicotiana tabacum
Olea europaea
Chionanthus rupicola
Hesperelaea palmeri
Erythranthe guttata
Salvia miltiorrhiza
Ajuga reptans
Castilleja paramensis
Utricularia reniformis
Dorcoceras hygrometricum
Haberlea rhodopensis
Epirixanthes elongata
Cucumis sativus
Cucurbita pepo
Citrullus lanatus
Morus notabilis
Populus tremula
Salix suchowensis
Cannabis sativa
Hemiptelea davidii
Ziziphus jujuba
Lagerstroemia indica
Eucalyptus grandis
Batis maritima
Turritis glabra
Boechera stricta
Arabidopsis thaliana
Raphanus sativus
Eruca vesicaria
Brassica rapa
Arabis alpina
Sinapis arvensis
Schrenkiella parvula
Capsella rubella
Malus domestica
Prunus avium
Nepenthes ventricosa
Fallopia multiflora
Spinacia oleracea
Chenopodium quinoa
Beta vulgaris
Silene vulgaris
Sesuvium portulacastrum
Lactuca serriola
Paraprenanthes diversifolia
Helianthus annuus
Chrysanthemum boreale
Codonopsis lanceolata
Platycodon grandiflorus
Vitis vinifera
Bombax ceiba
Gossypium trilobum
Hibiscus cannabinus
Corchorus olitorius
Camellia sinensis
Monotropa hypopitys
Vaccinium macrocarpon
Ricinus communis
Manihot esculenta
Hevea brasiliensis
Cercis canadensis
Tamarindus indica
Millettia pinnata
Phaseolus vulgaris
Vigna radiata
Glycine soja
Lotus japonicus
Castanospermum australe
Styphnolobium japonicum
Sophora flavescens
Ammopiptanthus nanus
Medicago truncatula
Bupleurum falcatum
Lophophytum mirabile
Leucaena trichandra
Acacia ligulata
Haematoxylum brasiletto
0.1

### atp4

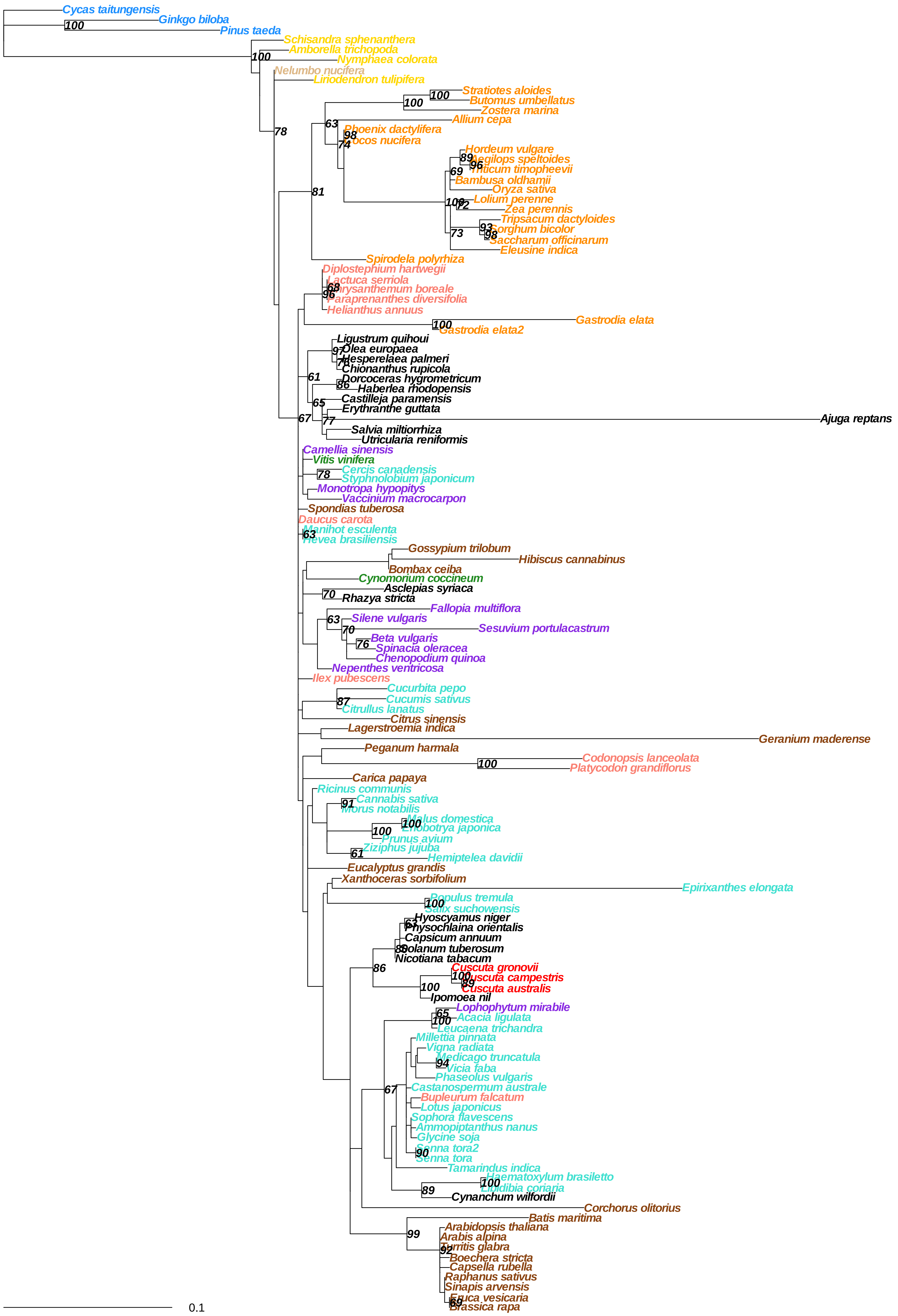

Cycas taitungensis
Ginkgo biloba
Pinus taeda
Schisandra sphenanthera
Amborella trichopoda
Nymphaea colorata
Nelumbo nucifera
Liriodendron tulipifera
Stratiotes aloides
Butomus umbellatus
Zostera marina
Allium cepa
Phoenix dactylifera
Cocos nucifera
Hordeum vulgare
Aegilops speltoides
Triticum timopheevii
Bambusa oldhamii
Oryza sativa
Lolium perenne
Zea perennis
Tripsacum dactyloides
Sorghum bicolor
Saccharum officinarum
Eleusine indica
Spirodela polyrhiza
Diplostephium hartwegii
Lactuca serriola
Chrysanthemum boreale
Paraprenanthes diversifolia
Helianthus annuus
Gastrodia elata
Gastrodia elata2
Ligustrum quihoui
Olea europaea
Hesperelaea palmeri
Chionanthus rupicola
Dorcoceras hygrometricum
Haberlea rhodopensis
Castilleja paramensis
Erythranthe guttata
Ajuga reptans
Salvia miltiorrhiza
Utricularia reniformis
Camellia sinensis
Vitis vinifera
Cercis canadensis
Styphnolobium japonicum
Monotropa hypopitys
Vaccinium macrocarpon
Spondias tuberosa
Daucus carota
Manihot esculenta
Hevea brasiliensis
Gossypium trilobum
Hibiscus cannabinus
Bombax ceiba
Cynomorium coccineum
Asclepias syriaca
Rhazya stricta
Fallopia multiflora
Silene vulgaris
Sesuvium portulacastrum
Beta vulgaris
Spinacia oleracea
Chenopodium quinoa
Nepenthes ventricosa
Ilex pubescens
Cucurbita pepo
Cucumis sativus
Citrullus lanatus
Citrus sinensis
Lagerstroemia indica
Geranium maderense
Peganum harmala
Codonopsis lanceolata
Platycodon grandiflorus
Carica papaya
Ricinus communis
Cannabis sativa
Morus notabilis
Malus domestica
Eriobotrya japonica
Prunus avium
Ziziphus jujuba
Hemiptelea davidii
Eucalyptus grandis
Xanthoceras sorbifolium
Epirixanthes elongata
Populus tremula
Salix suchowensis
Hyoscyamus niger
Physochlaina orientalis
Capsicum annuum
Solanum tuberosum
Nicotiana tabacum
Cuscuta gronovii
Cuscuta campestris
Cuscuta australis
Ipomoea nil
Lophophytum mirabile
Acacia ligulata
Leucaena trichandra
Millettia pinnata
Vigna radiata
Medicago truncatula
Vicia faba
Phaseolus vulgaris
Castanospermum australe
Bupleurum falcatum
Lotus japonicus
Sophora flavescens
Ammopiptanthus nanus
Glycine soja
Senna tora2
Senna tora
Tamarindus indica
Haematoxylum brasiletto
Libidibia coriaria
Cynanchum wilfordii
Corchorus olitorius
Batis maritima
Arabidopsis thaliana
Arabis alpina
Turritis glabra
Boechera stricta
Capsella rubella
Raphanus sativus
Sinapis arvensis
Eruca vesicaria
Brassica rapa
0.1

**atp6**

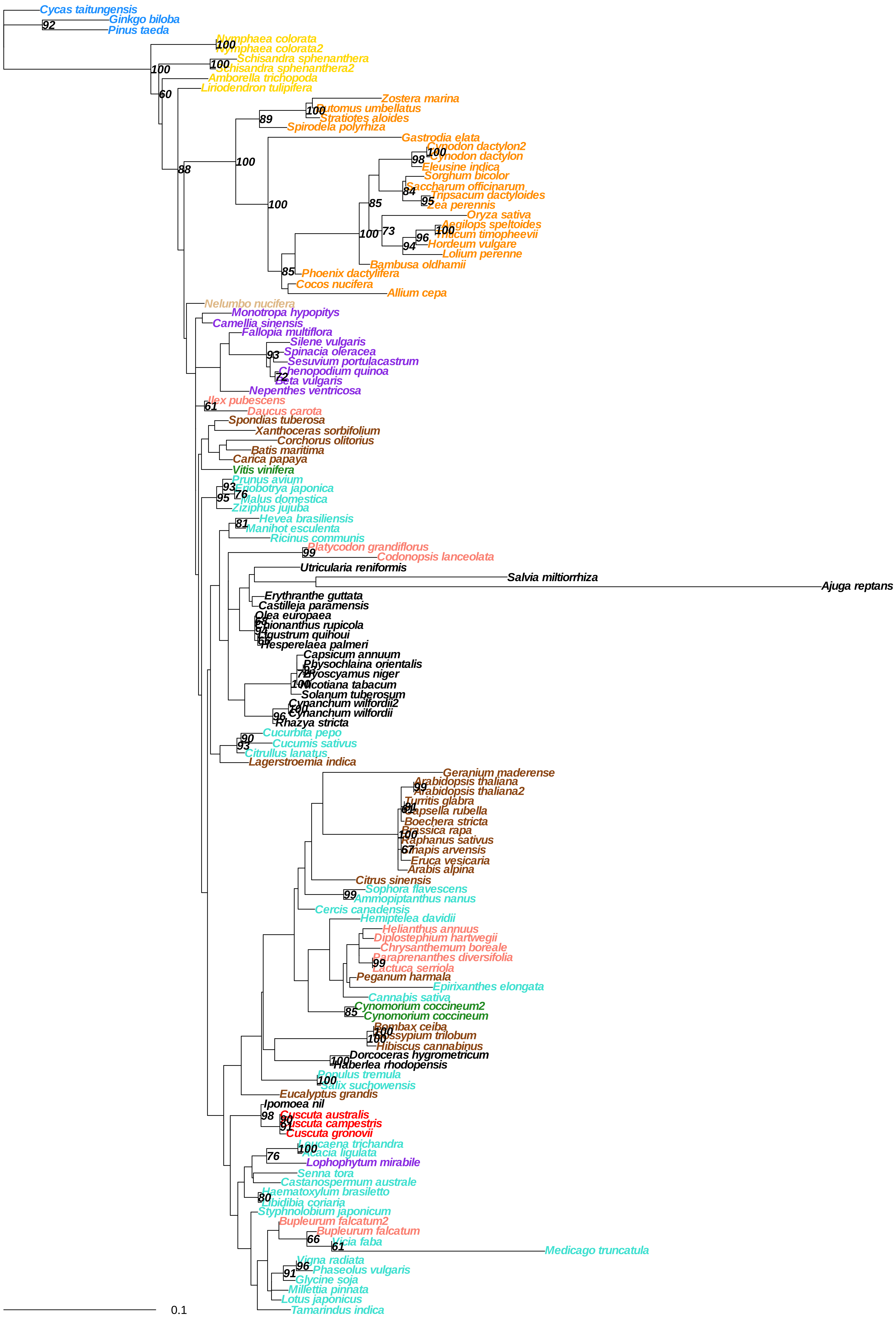
Cycas taitungensis
Ginkgo biloba
Pinus taeda
Nymphaea colorata
Nymphaea colorata2
Schisandra sphenanthera
Schisandra sphenanthera2
Amborella trichopoda
Liriodendron tulipifera
Zostera marina
Butomus umbellatus
Stratiotes aloides
Spirodela polyrhiza
Gastrodia elata
Cynodon dactylon2
Cynodon dactylon
Eleusine indica
Sorghum bicolor
Saccharum officinarum
Tripsacum dactyloides
Zea perennis
Oryza sativa
Aegilops speltoides
Triticum timopheevii
Hordeum vulgare
Lolium perenne
Bambusa oldhamii
Phoenix dactylifera
Cocos nucifera
Allium cepa
Nelumbo nucifera
Monotropa hypopitys
Camellia sinensis
Fallopia multiflora
Silene vulgaris
Spinacia oleracea
Sesuvium portulacastrum
Chenopodium quinoa
Beta vulgaris
Nepenthes ventricosa
Ilex pubescens
Daucus carota
Spondias tuberosa
Xanthoceras sorbifolium
Corchorus olitorius
Batis maritima
Carica papaya
Vitis vinifera
Prunus avium
Eriobotrya japonica
Malus domestica
Ziziphus jujuba
Hevea brasiliensis
Manihot esculenta
Ricinus communis
Platycodon grandiflorus
Codonopsis lanceolata
Utricularia reniformis
Salvia miltiorrhiza
Ajuga reptans
Erythranthe guttata
Castilleja paramensis
Olea europaea
Chionanthus rupicola
Ligustrum quihoui
Hesperelaea palmeri
Capsicum annuum
Physochlaina orientalis
Hyoscyamus niger
Nicotiana tabacum
Solanum tuberosum
Cynanchum wilfordii2
Cynanchum wilfordii
Rhazya stricta
Cucurbita pepo
Cucumis sativus
Citrullus lanatus
Lagerstroemia indica
Geranium maderense
Arabidopsis thaliana
Arabidopsis thaliana2
Turritis glabra
Capsella rubella
Boechera stricta
Brassica rapa
Raphanus sativus
Sinapis arvensis
Eruca vesicaria
Arabis alpina
Citrus sinensis
Sophora flavescens
Ammopiptanthus nanus
Cercis canadensis
Hemiptelea davidii
Helianthus annuus
Diplostephium hartwegii
Chrysanthemum boreale
Paraprenanthes diversifolia
Lactuca serriola
Peganum harmala
Epirixanthes elongata
Cannabis sativa
Cynomorium coccineum2
Cynomorium coccineum
Bombax ceiba
Gossypium trilobum
Hibiscus cannabinus
Dorcoceras hygrometricum
Haberlea rhodopensis
Populus tremula
Salix suchowensis
Eucalyptus grandis
Ipomoea nil
Cuscuta australis
Cuscuta campestris
Cuscuta gronovii
Leucaena trichandra
Acacia ligulata
Lophophytum mirabile
Senna tora
Castanospermum australe
Haematoxylum brasiletto
Libidibia coriaria
Styphnolobium japonicum
Bupleurum falcatum2
Bupleurum falcatum
Vicia faba
Medicago truncatula
Vigna radiata
Phaseolus vulgaris
Glycine soja
Millettia pinnata
Lotus japonicus
Tamarindus indica
0.1

**atp8**

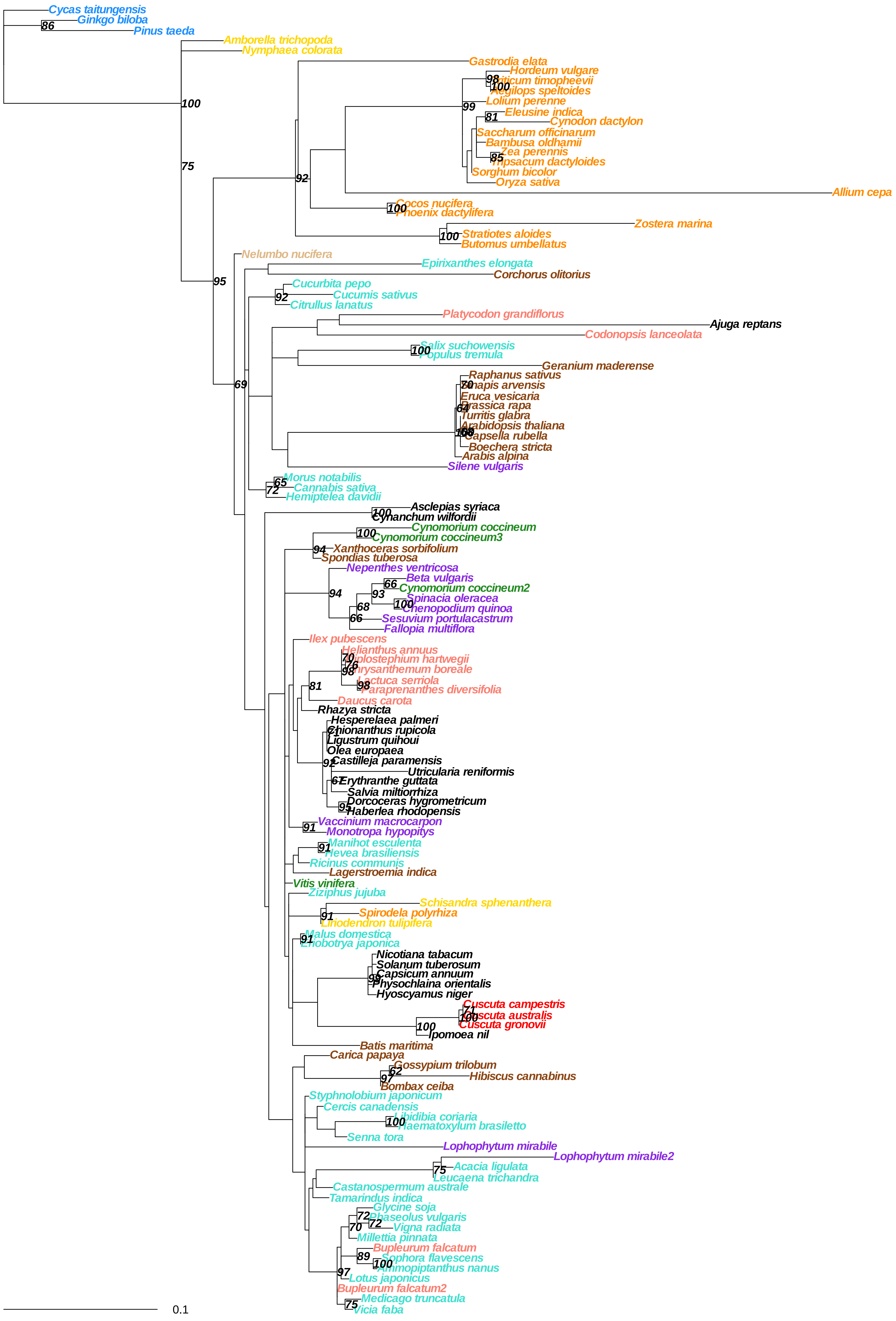

#### atp9

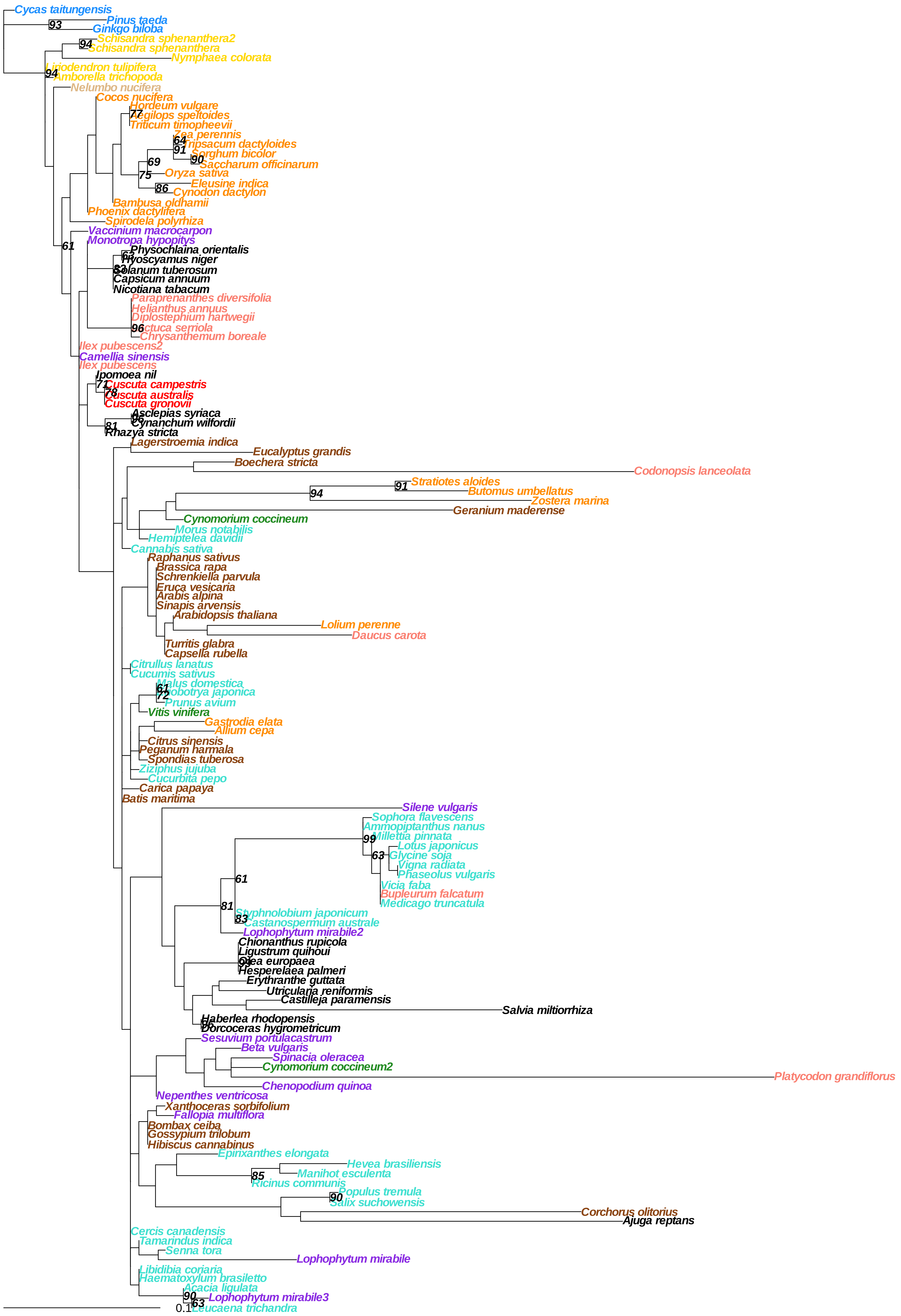
Cycas taitungensis
Pinus taeda
Ginkgo biloba
Schisandra sphenanthera2
Schisandra sphenanthera
Nymphaea colorata
Liriodendron tulipifera
Amborella trichopoda
Nelumbo nucifera
Cocos nucifera
Hordeum vulgare
Aegilops speltoides
Triticum timopheevii
Zea perennis
Tripsacum dactyloides
Sorghum bicolor
Saccharum officinarum
Oryza sativa
Eleusine indica
Cynodon dactylon
Bambusa oldhamii
Phoenix dactylifera
Spirodela polyrhiza
Vaccinium macrocarpon
Monotropa hypopitys
Physochlaina orientalis
Hyoscyamus niger
Solanum tuberosum
Capsicum annuum
Nicotiana tabacum
Paraprenanthes diversifolia
Helianthus annuus
Diplostephium hartwegii
Lactuca serriola
Chrysanthemum boreale
Ilex pubescens2
Camellia sinensis
Ilex pubescens
Ipomoea nil
Cuscuta campestris
Cuscuta australis
Cuscuta gronovii
Asclepias syriaca
Cynanchum wilfordii
Rhazya stricta
Lagerstroemia indica
Eucalyptus grandis
Boechera stricta
Codonopsis lanceolata
Stratiotes aloides
Butomus umbellatus
Zostera marina
Geranium maderense
Cynomorium coccineum
Morus notabilis
Hemiptelea davidii
Cannabis sativa
Raphanus sativus
Brassica rapa
Schrenkiella parvula
Eruca vesicaria
Arabis alpina
Sinapis arvensis
Arabidopsis thaliana
Lolium perenne
Daucus carota
Turritis glabra
Capsella rubella
Citrullus lanatus
Cucumis sativus
Malus domestica
Eriobotrya japonica
Prunus avium
Vitis vinifera
Gastrodia elata
Allium cepa
Citrus sinensis
Peganum harmala
Spondias tuberosa
Ziziphus jujuba
Cucurbita pepo
Carica papaya
Batis maritima
Silene vulgaris
Sophora flavescens
Ammopiptanthus nanus
Millettia pinnata
Lotus japonicus
Glycine soja
Vigna radiata
Phaseolus vulgaris
Vicia faba
Bupleurum falcatum
Medicago truncatula
Styphnolobium japonicum
Castanospermum australe
Lophophytum mirabile2
Chionanthus rupicola
Ligustrum quihoui
Olea europaea
Hesperelaea palmeri
Erythranthe guttata
Utricularia reniformis
Castilleja paramensis
Salvia miltiorrhiza
Haberlea rhodopensis
Dorcoceras hygrometricum
Sesuvium portulacastrum
Beta vulgaris
Spinacia oleracea
Cynomorium coccineum2
Platycodon grandiflorus
Chenopodium quinoa
Nepenthes ventricosa
Xanthoceras sorbifolium
Fallopia multiflora
Bombax ceiba
Gossypium trilobum
Hibiscus cannabinus
Epirixanthes elongata
Hevea brasiliensis
Manihot esculenta
Ricinus communis
Populus tremula
Salix suchowensis
Corchorus olitorius
Ajuga reptans
Cercis canadensis
Tamarindus indica
Senna tora
Lophophytum mirabile
Libidibia coriaria
Haematoxylum brasiletto
Acacia ligulata
Lophophytum mirabile3
Leucaena trichandra
0.1

### ccmb

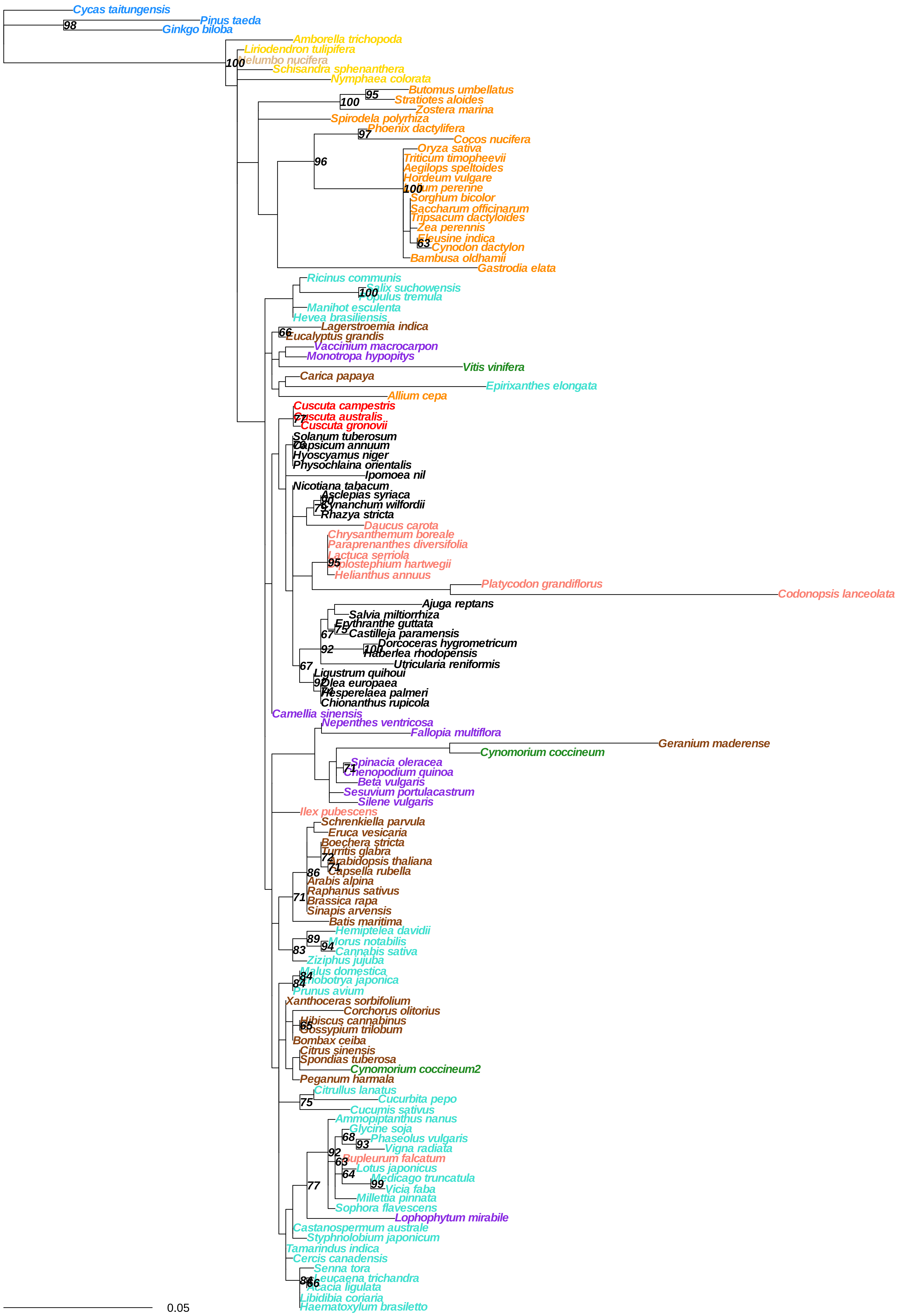

Cycas taitungensis
Pinus taeda
Ginkgo biloba
Amborella trichopoda
Liriodendron tulipifera
Nelumbo nucifera
Schisandra sphenanthera
Nymphaea colorata
Butomus umbellatus
Stratiotes aloides
Zostera marina
Spirodela polyrhiza
Phoenix dactylifera
Cocos nucifera
Oryza sativa
Triticum timopheevii
Aegilops speltoides
Hordeum vulgare
Lolium perenne
Sorghum bicolor
Saccharum officinarum
Tripsacum dactyloides
Zea perennis
Eleusine indica
Cynodon dactylon
Bambusa oldhamii
Gastrodia elata
Ricinus communis
Salix suchowensis
Populus tremula
Manihot esculenta
Hevea brasiliensis
Lagerstroemia indica
Eucalyptus grandis
Vaccinium macrocarpon
Monotropa hypopitys
Vitis vinifera
Carica papaya
Epirixanthes elongata
Allium cepa
Cuscuta campestris
Cuscuta australis
Cuscuta gronovii
Solanum tuberosum
Capsicum annuum
Hyoscyamus niger
Physochlaina orientalis
Ipomoea nil
Nicotiana tabacum
Asclepias syriaca
Cynanchum wilfordii
Rhazya stricta
Daucus carota
Chrysanthemum boreale
Paraprenanthes diversifolia
Lactuca serriola
Diplostephium hartwegii
Helianthus annuus
Platycodon grandiflorus
Codonopsis lanceolata
Ajuga reptans
Salvia miltiorrhiza
Erythranthe guttata
Castilleja paramensis
Dorcoceras hygrometricum
Haberlea rhodopensis
Utricularia reniformis
Ligustrum quihoui
Olea europaea
Hesperelaea palmeri
Chionanthus rupicola
Camellia sinensis
Nepenthes ventricosa
Fallopia multiflora
Geranium maderense
Cynomorium coccineum
Spinacia oleracea
Chenopodium quinoa
Beta vulgaris
Sesuvium portulacastrum
Silene vulgaris
Ilex pubescens
Schrenkiella parvula
Eruca vesicaria
Boechera stricta
Turritis glabra
Arabidopsis thaliana
Capsella rubella
Arabis alpina
Raphanus sativus
Brassica rapa
Sinapis arvensis
Batis maritima
Hemiptelea davidii
Morus notabilis
Cannabis sativa
Ziziphus jujuba
Malus domestica
Eriobotrya japonica
Prunus avium
Xanthoceras sorbifolium
Corchorus olitorius
Hibiscus cannabinus
Gossypium trilobum
Bombax ceiba
Citrus sinensis
Spondias tuberosa
Cynomorium coccineum2
Peganum harmala
Citrullus lanatus
Cucurbita pepo
Cucumis sativus
Ammopiptanthus nanus
Glycine soja
Phaseolus vulgaris
Vigna radiata
Bupleurum falcatum
Lotus japonicus
Medicago truncatula
Vicia faba
Millettia pinnata
Sophora flavescens
Lophophytum mirabile
Castanospermum australe
Styphnolobium japonicum
Tamarindus indica
Cercis canadensis
Senna tora
Leucaena trichandra
Acacia ligulata
Libidibia coriaria
Haematoxylum brasiletto
0.05

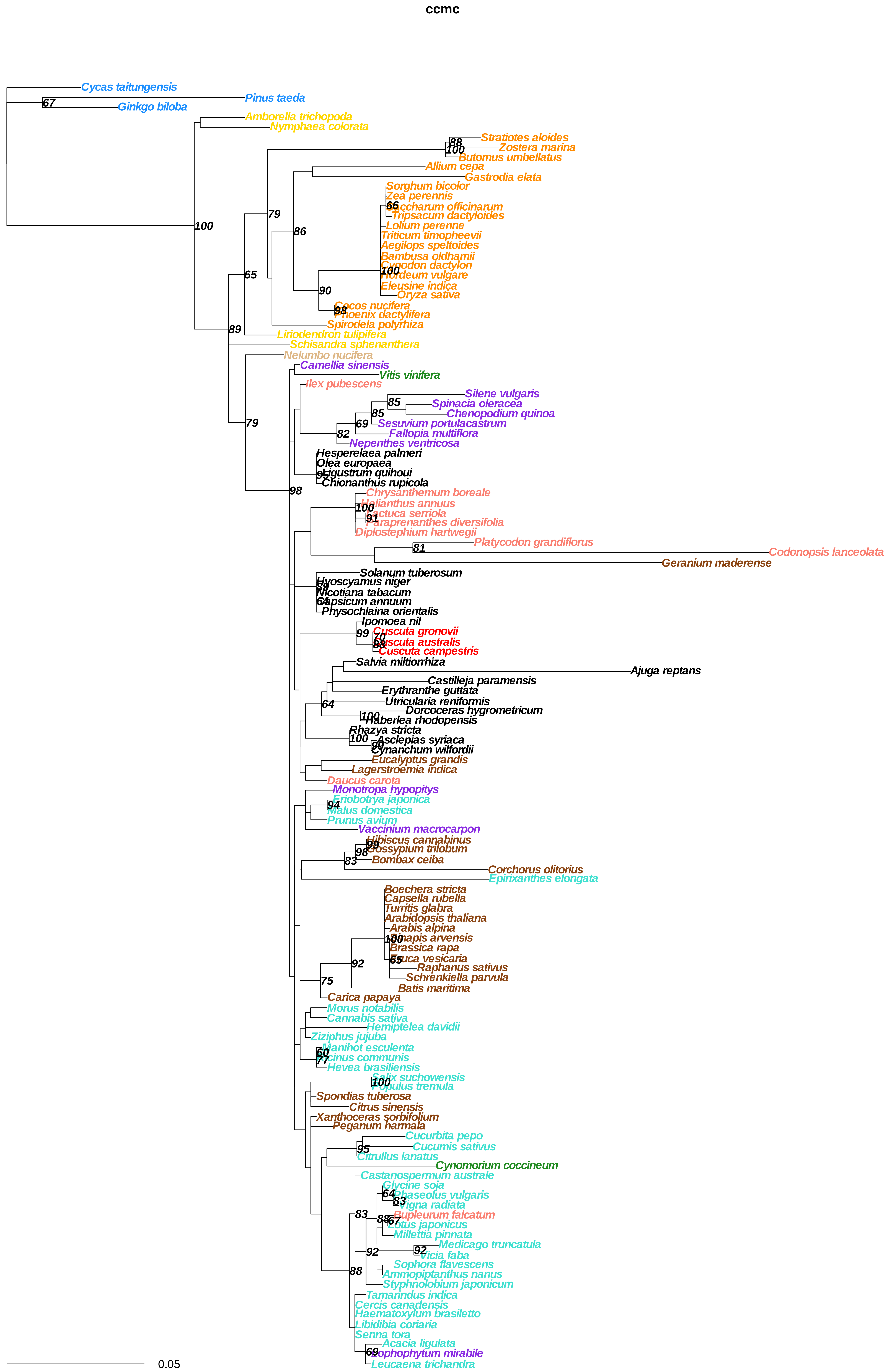
ccmc
Cycas taitungensis
Pinus taeda
Ginkgo biloba
Amborella trichopoda
Nymphaea colorata
Stratiotes aloides
Zostera marina
Butomus umbellatus
Allium cepa
Gastrodia elata
Sorghum bicolor
Zea perennis
Saccharum officinarum
Tripsacum dactyloides
Lolium perenne
Triticum timopheevii
Aegilops speltoides
Bambusa oldhamii
Cynodon dactylon
Hordeum vulgare
Eleusine indica
Oryza sativa
Cocos nucifera
Phoenix dactylifera
Spirodela polyrhiza
Liriodendron tulipifera
Schisandra sphenanthera
Nelumbo nucifera
Camellia sinensis
Vitis vinifera
Ilex pubescens
Silene vulgaris
Spinacia oleracea
Chenopodium quinoa
Sesuvium portulacastrum
Fallopia multiflora
Nepenthes ventricosa
Hesperelaea palmeri
Olea europaea
Ligustrum quihoui
Chionanthus rupicola
Chrysanthemum boreale
Helianthus annuus
Lactuca serriola
Paraprenanthes diversifolia
Diplostephium hartwegii
Platycodon grandiflorus
Codonopsis lanceolata
Geranium maderense
Solanum tuberosum
Hyoscyamus niger
Nicotiana tabacum
Capsicum annuum
Physochlaina orientalis
Ipomoea nil
Cuscuta gronovii
Cuscuta australis
Cuscuta campestris
Salvia miltiorrhiza
Ajuga reptans
Castilleja paramensis
Erythranthe guttata
Utricularia reniformis
Dorcoceras hygrometricum
Haberlea rhodopensis
Rhazya stricta
Asclepias syriaca
Cynanchum wilfordii
Eucalyptus grandis
Lagerstroemia indica
Daucus carota
Monotropa hypopitys
Eriobotrya japonica
Malus domestica
Prunus avium
Vaccinium macrocarpon
Hibiscus cannabinus
Gossypium trilobum
Bombax ceiba
Corchorus olitorius
Epirixanthes elongata
Boechera stricta
Capsella rubella
Turritis glabra
Arabidopsis thaliana
Arabis alpina
Sinapis arvensis
Brassica rapa
Eruca vesicaria
Raphanus sativus
Schrenkiella parvula
Batis maritima
Carica papaya
Morus notabilis
Cannabis sativa
Hemiptelea davidii
Ziziphus jujuba
Manihot esculenta
Ricinus communis
Hevea brasiliensis
Salix suchowensis
Populus tremula
Spondias tuberosa
Citrus sinensis
Xanthoceras sorbifolium
Peganum harmala
Cucurbita pepo
Cucumis sativus
Citrullus lanatus
Cynomorium coccineum
Castanospermum australe
Glycine soja
Phaseolus vulgaris
Vigna radiata
Bupleurum falcatum
Lotus japonicus
Millettia pinnata
Medicago truncatula
Vicia faba
Sophora flavescens
Ammopiptanthus nanus
Styphnolobium japonicum
Tamarindus indica
Cercis canadensis
Haematoxylum brasiletto
Libidibia coriaria
Senna tora
Acacia ligulata
Lophophytum mirabile
Leucaena trichandra
0.05

ccmfc

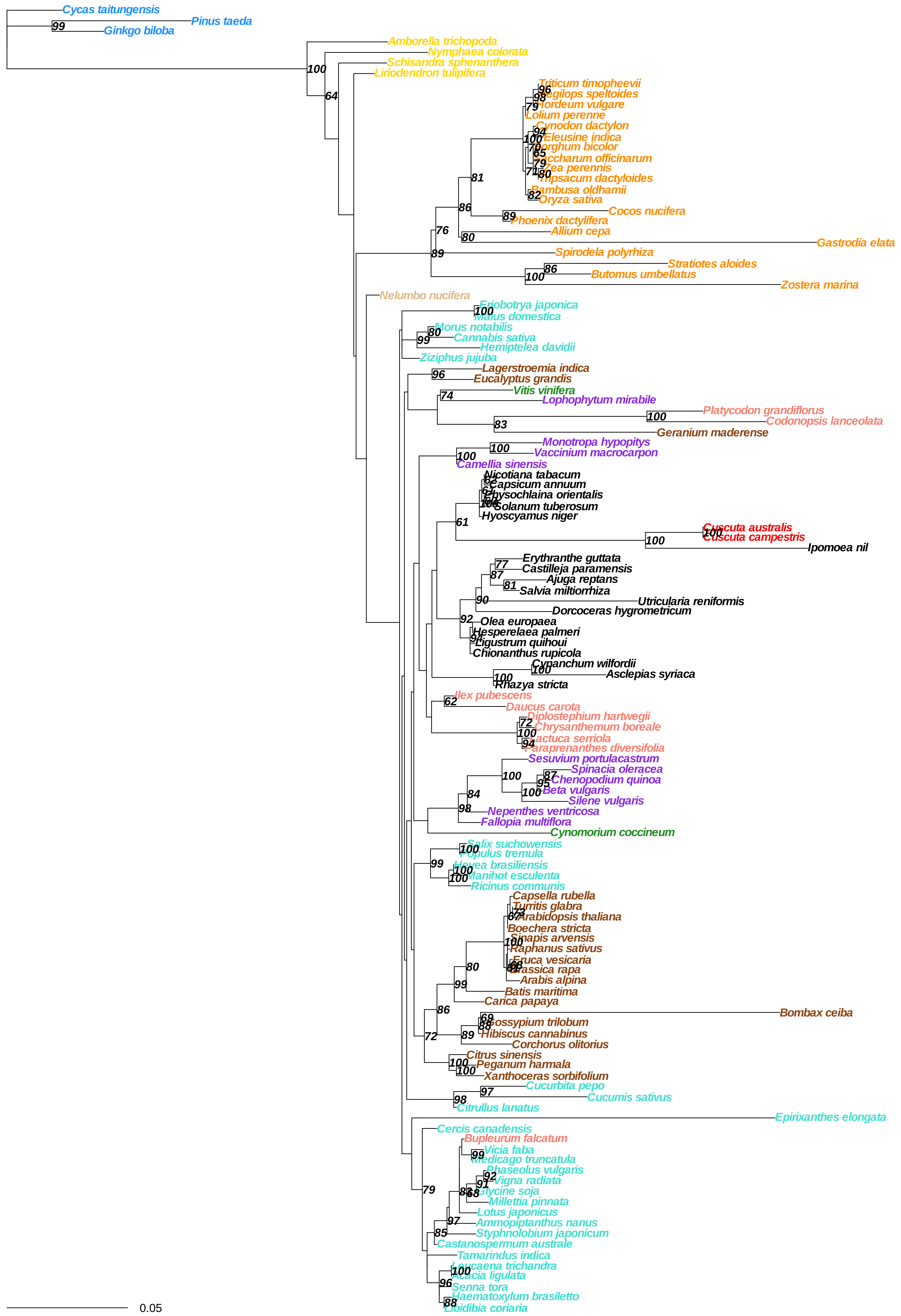

Cycas taitungensis
Pinus taeda
Ginkgo biloba
Amborella trichopoda
Nymphaea colorata
Schisandra sphenanthera
Liriodendron tulipifera
Triticum timopheevii
Aegilops speltoides
Hordeum vulgare
Lolium perenne
Cynodon dactylon
Eleusine indica
Sorghum bicolor
Saccharum officinarum
Zea perennis
Tripsacum dactyloides
Bambusa oldhamii
Oryza sativa
Cocos nucifera
Phoenix dactylifera
Allium cepa
Gastrodia elata
Spirodela polyrhiza
Stratiotes aloides
Butomus umbellatus
Zostera marina
Nelumbo nucifera
Eriobotrya japonica
Malus domestica
Morus notabilis
Cannabis sativa
Hemiptelea davidii
Ziziphus jujuba
Lagerstroemia indica
Eucalyptus grandis
Vitis vinifera
Lophophytum mirabile
Platycodon grandiflorus
Codonopsis lanceolata
Geranium maderense
Monotropa hypopitys
Vaccinium macrocarpon
Camellia sinensis
Nicotiana tabacum
Capsicum annuum
Physochlaina orientalis
Solanum tuberosum
Hyoscyamus niger
Cuscuta australis
Cuscuta campestris
Ipomoea nil
Erythranthe guttata
Castilleja paramensis
Ajuga reptans
Salvia miltiorrhiza
Utricularia reniformis
Dorcoceras hygrometricum
Olea europaea
Hesperelaea palmeri
Ligustrum quihoui
Chionanthus rupicola
Cynanchum wilfordii
Asclepias syriaca
Rhazya stricta
Ilex pubescens
Daucus carota
Diplostephium hartwegii
Chrysanthemum boreale
Lactuca serriola
Paraprenanthes diversifolia
Sesuvium portulacastrum
Spinacia oleracea
Chenopodium quinoa
Beta vulgaris
Silene vulgaris
Nepenthes ventricosa
Fallopia multiflora
Cynomorium coccineum
Salix suchowensis
Populus tremula
Hevea brasiliensis
Manihot esculenta
Ricinus communis
Capsella rubella
Turritis glabra
Arabidopsis thaliana
Boechera stricta
Sinapis arvensis
Raphanus sativus
Eruca vesicaria
Brassica rapa
Arabis alpina
Batis maritima
Carica papaya
Bombax ceiba
Gossypium trilobum
Hibiscus cannabinus
Corchorus olitorius
Citrus sinensis
Peganum harmala
Xanthoceras sorbifolium
Cucurbita pepo
Cucumis sativus
Citrullus lanatus
Epirixanthes elongata
Cercis canadensis
Bupleurum falcatum
Vicia faba
Medicago truncatula
Phaseolus vulgaris
Vigna radiata
Glycine soja
Millettia pinnata
Lotus japonicus
Ammopiptanthus nanus
Styphnolobium japonicum
Castanospermum australe
Tamarindus indica
Leucaena trichandra
Acacia ligulata
Senna tora
Haematoxylum brasiletto
Libidibia coriaria
0.05

### ccmfn

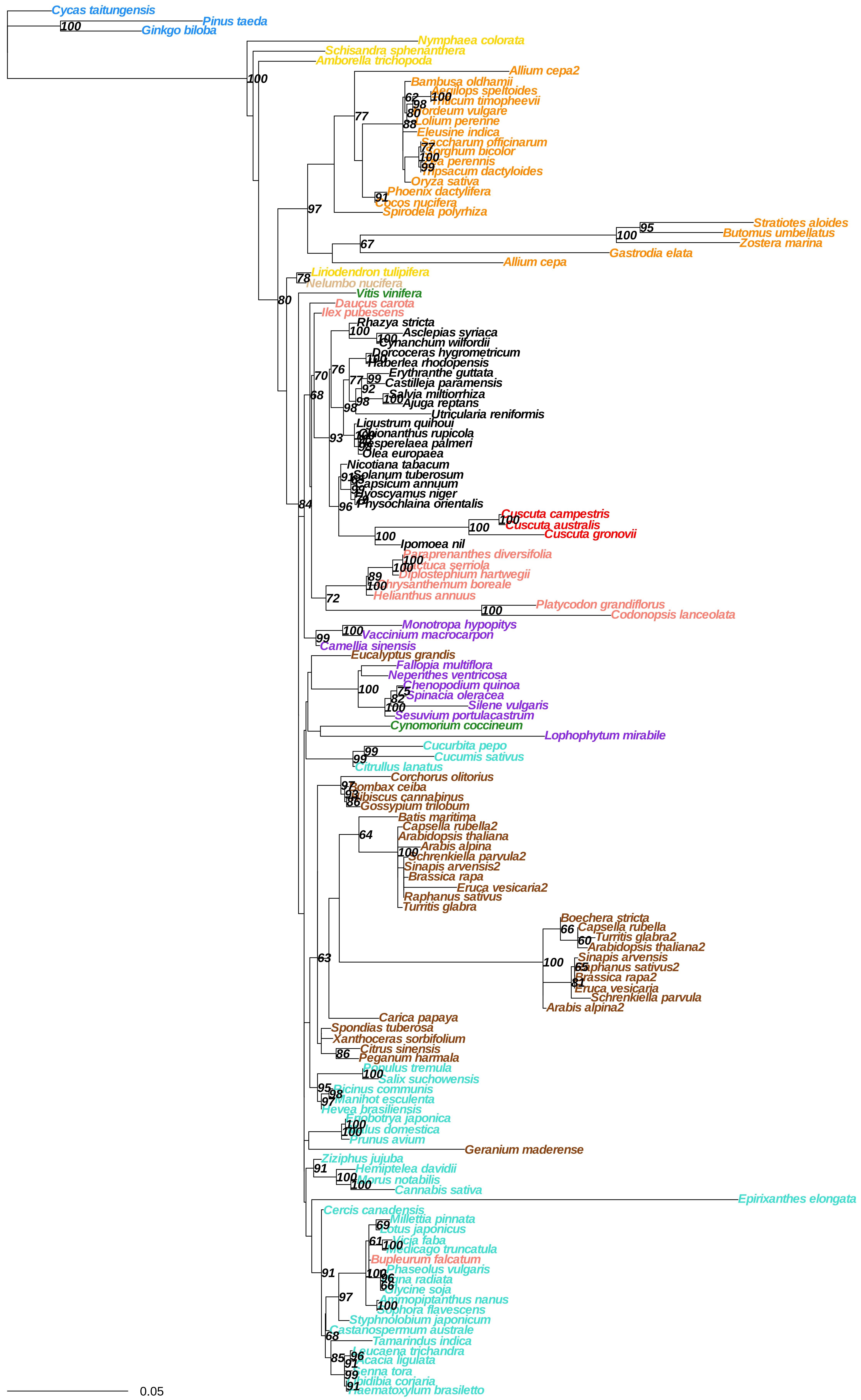

Cycas taitungensis
Pinus taeda
Ginkgo biloba
Nymphaea colorata
Schisandra sphenanthera
Amborella trichopoda
Allium cepa2
Bambusa oldhamii
Aegilops speltoides
Triticum timopheevii
Hordeum vulgare
Lolium perenne
Eleusine indica
Saccharum officinarum
Sorghum bicolor
Zea perennis
Tripsacum dactyloides
Oryza sativa
Phoenix dactylifera
Cocos nucifera
Spirodela polyrhiza
Stratiotes aloides
Butomus umbellatus
Zostera marina
Gastrodia elata
Allium cepa
Liriodendron tulipifera
Nelumbo nucifera
Vitis vinifera
Daucus carota
Ilex pubescens
Rhazya stricta
Asclepias syriaca
Cynanchum wilfordii
Dorcoceras hygrometricum
Haberlea rhodopensis
Erythranthe guttata
Castilleja paramensis
Salvia miltiorrhiza
Ajuga reptans
Utricularia reniformis
Ligustrum quihoui
Chionanthus rupicola
Hesperelaea palmeri
Olea europaea
Nicotiana tabacum
Solanum tuberosum
Capsicum annuum
Hyoscyamus niger
Physochlaina orientalis
Cuscuta campestris
Cuscuta australis
Cuscuta gronovii
Ipomoea nil
Paraprenanthes diversifolia
Lactuca serriola
Diplostephium hartwegii
Chrysanthemum boreale
Helianthus annuus
Platycodon grandiflorus
Codonopsis lanceolata
Monotropa hypopitys
Vaccinium macrocarpon
Camellia sinensis
Eucalyptus grandis
Fallopia multiflora
Nepenthes ventricosa
Chenopodium quinoa
Spinacia oleracea
Silene vulgaris
Sesuvium portulacastrum
Cynomorium coccineum
Lophophytum mirabile
Cucurbita pepo
Cucumis sativus
Citrullus lanatus
Corchorus olitorius
Bombax ceiba
Hibiscus cannabinus
Gossypium trilobum
Batis maritima
Capsella rubella2
Arabidopsis thaliana
Arabis alpina
Schrenkiella parvula2
Sinapis arvensis2
Brassica rapa
Eruca vesicaria2
Raphanus sativus
Turritis glabra
Boechera stricta
Capsella rubella
Turritis glabra2
Arabidopsis thaliana2
Sinapis arvensis
Raphanus sativus2
Brassica rapa2
Eruca vesicaria
Schrenkiella parvula
Arabis alpina2
Carica papaya
Spondias tuberosa
Xanthoceras sorbifolium
Citrus sinensis
Peganum harmala
Populus tremula
Salix suchowensis
Ricinus communis
Manihot esculenta
Hevea brasiliensis
Eriobotrya japonica
Malus domestica
Prunus avium
Geranium maderense
Ziziphus jujuba
Hemiptelea davidii
Morus notabilis
Cannabis sativa
Epirixanthes elongata
Cercis canadensis
Millettia pinnata
Lotus japonicus
Vicia faba
Medicago truncatula
Bupleurum falcatum
Phaseolus vulgaris
Vigna radiata
Glycine soja
Ammopiptanthus nanus
Sophora flavescens
Styphnolobium japonicum
Castanospermum australe
Tamarindus indica
Leucaena trichandra
Acacia ligulata
Senna tora
Libidibia coriaria
Haematoxylum brasiletto
0.05

**cob**

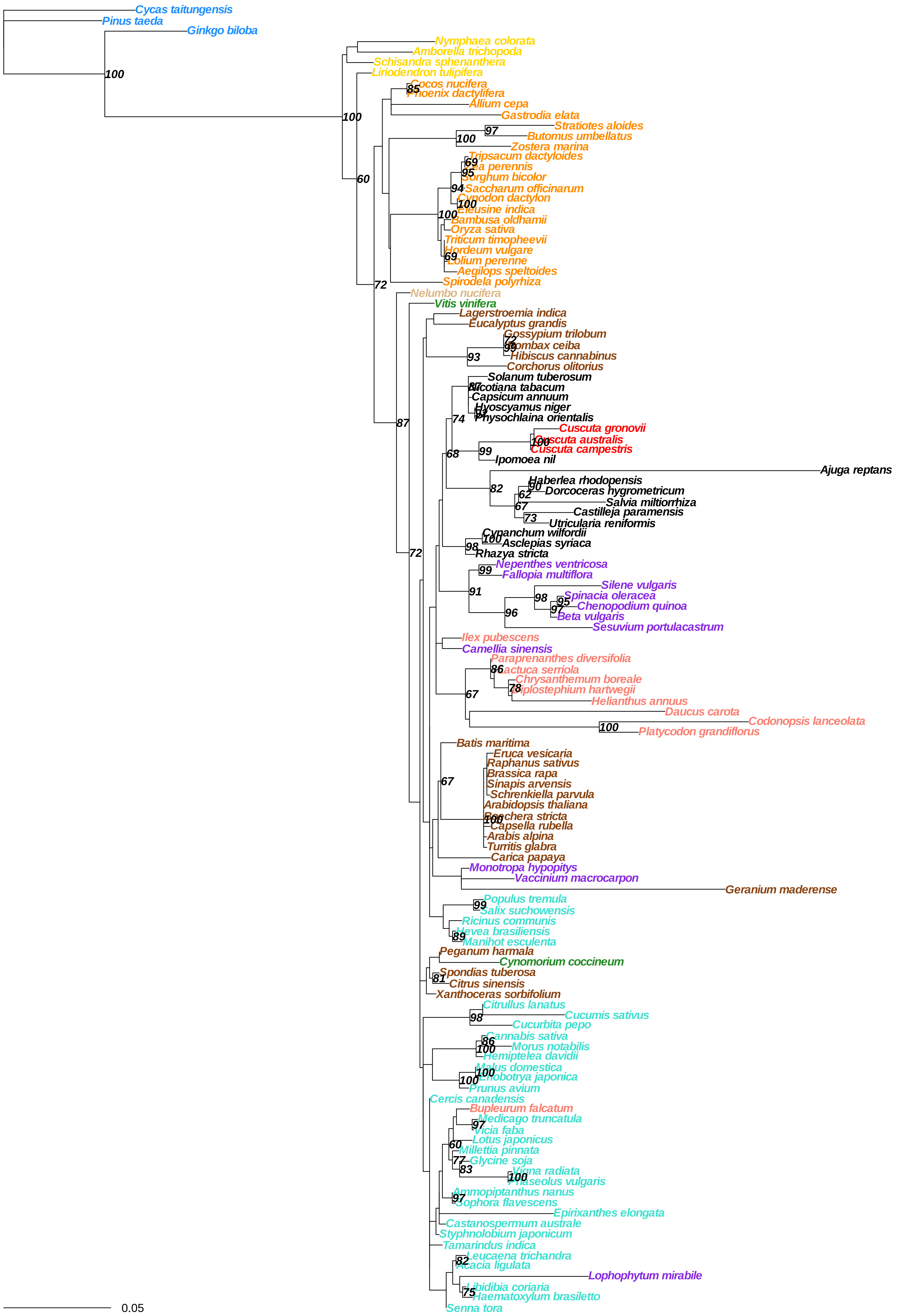

Cycas taitungensis
Pinus taeda
Ginkgo biloba
100
Nymphaea colorata
Amborella trichopoda
Schisandra sphenanthera
Liriodendron tulipifera
100
60
Cocos nucifera
85
Phoenix dactylifera
Allium cepa
Gastrodia elata
Stratiotes aloides
97
Butomus umbellatus
100
Zostera marina
Tripsacum dactyloides
69
Zea perennis
95
Sorghum bicolor
94
Saccharum officinarum
Cynodon dactylon
100
Eleusine indica
100
Bambusa oldhamii
Oryza sativa
Triticum timopheevii
Hordeum vulgare
69
Lolium perenne
Aegilops speltoides
Spirodela polyrhiza
72
Nelumbo nucifera
Vitis vinifera
Lagerstroemia indica
Eucalyptus grandis
Gossypium trilobum
72
Bombax ceiba
95
93
Hibiscus cannabinus
Corchorus olitorius
Solanum tuberosum
87
Nicotiana tabacum
Capsicum annuum
Hyoscyamus niger
94
Physochlaina orientalis
74
87
Cuscuta gronovii
100
Cuscuta australis
Cuscuta campestris
68
99
Ipomoea nil
Ajuga reptans
82
Haberlea rhodopensis
90
Dorcoceras hygrometricum
62
Salvia miltiorrhiza
67
Castilleja paramensis
73
Utricularia reniformis
Cynanchum wilfordii
100
Asclepias syriaca
98
Rhazya stricta
72
Nepenthes ventricosa
99
Fallopia multiflora
91
Silene vulgaris
98
Spinacia oleracea
95
Chenopodium quinoa
97
96
Beta vulgaris
Sesuvium portulacastrum
Ilex pubescens
Camellia sinensis
Paraprenanthes diversifolia
86
Lactuca serriola
Chrysanthemum boreale
78
Diplostephium hartwegii
67
Helianthus annuus
Daucus carota
Codonopsis lanceolata
100
Platycodon grandiflorus
Batis maritima
Eruca vesicaria
Raphanus sativus
Brassica rapa
67
Sinapis arvensis
Schrenkiella parvula
Arabidopsis thaliana
Boechera stricta
100
Capsella rubella
Arabis alpina
Turritis glabra
Carica papaya
Monotropa hypopitys
Vaccinium macrocarpon
Geranium maderense
Populus tremula
99
Salix suchowensis
Ricinus communis
Hevea brasiliensis
89
Manihot esculenta
Peganum harmala
Cynomorium coccineum
Spondias tuberosa
81
Citrus sinensis
Xanthoceras sorbifolium
Citrullus lanatus
98
Cucumis sativus
Cucurbita pepo
86
Cannabis sativa
100
Morus notabilis
Hemiptelea davidii
Malus domestica
100
Eriobotrya japonica
100
Prunus avium
Cercis canadensis
Bupleurum falcatum
97
Medicago truncatula
Vicia faba
Lotus japonicus
60
Millettia pinnata
77
Glycine soja
83
Vigna radiata
100
Phaseolus vulgaris
Ammopiptanthus nanus
97
Sophora flavescens
Epirixanthes elongata
Castanospermum australe
Styphnolobium japonicum
Tamarindus indica
82
Leucaena trichandra
Acacia ligulata
Lophophytum mirabile
Libidibia coriaria
75
Haematoxylum brasiletto
Senna tora
0.05

**cox1**

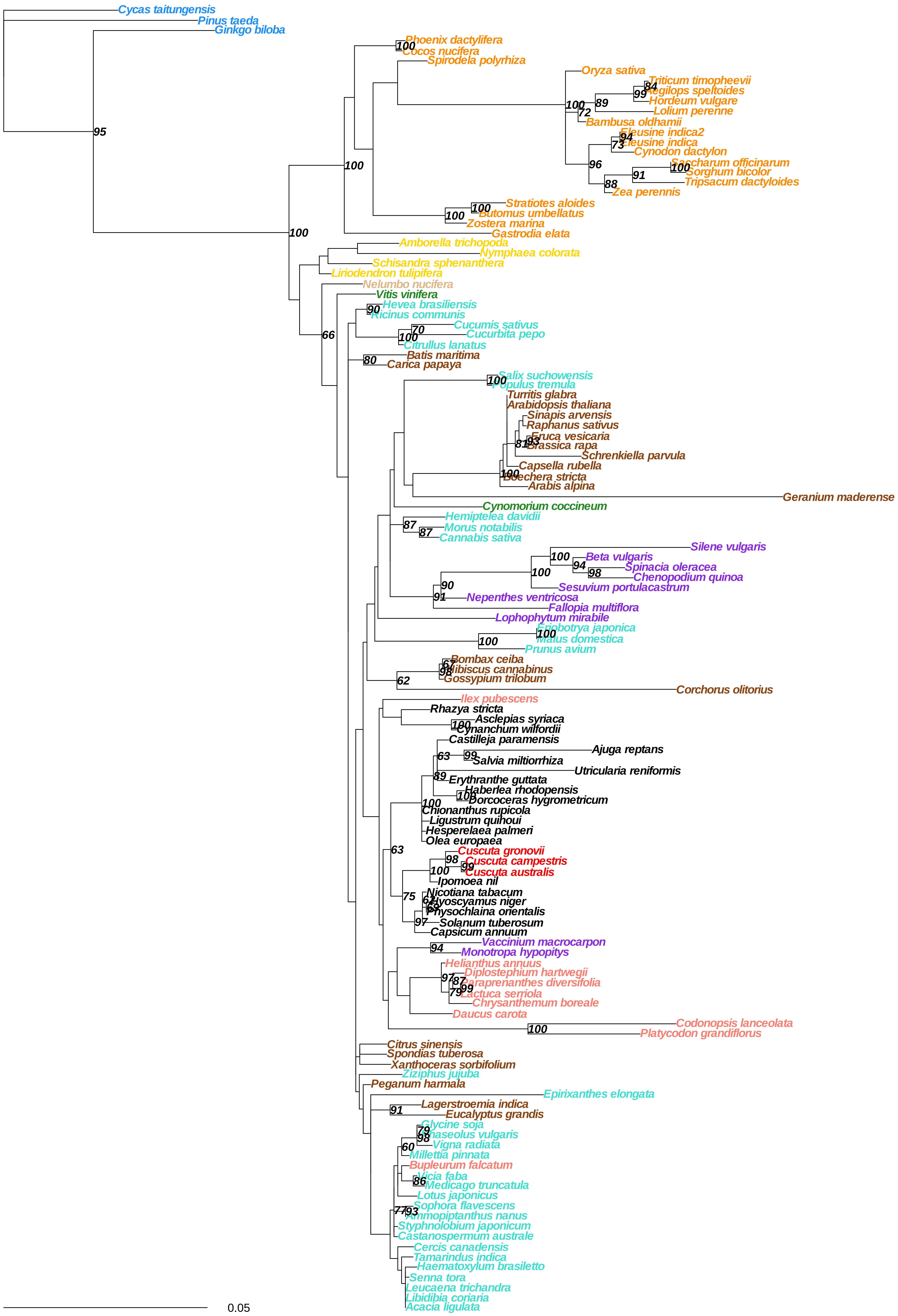
Cycas taitungensis
Pinus taeda
Ginkgo biloba
Phoenix dactylifera
Cocos nucifera
Spirodela polyrhiza
Oryza sativa
Triticum timopheevii
Aegilops speltoides
Hordeum vulgare
Lolium perenne
Bambusa oldhamii
Eleusine indica2
Eleusine indica
Cynodon dactylon
Saccharum officinarum
Sorghum bicolor
Tripsacum dactyloides
Zea perennis
Stratiotes aloides
Butomus umbellatus
Zostera marina
Gastrodia elata
Amborella trichopoda
Nymphaea colorata
Schisandra sphenanthera
Liriodendron tulipifera
Nelumbo nucifera
Vitis vinifera
Hevea brasiliensis
Ricinus communis
Cucumis sativus
Cucurbita pepo
Citrullus lanatus
Batis maritima
Carica papaya
Salix suchowensis
Populus tremula
Turritis glabra
Arabidopsis thaliana
Sinapis arvensis
Raphanus sativus
Eruca vesicaria
Brassica rapa
Schrenkiella parvula
Capsella rubella
Boechera stricta
Arabis alpina
Geranium maderense
Cynomorium coccineum
Hemiptelea davidii
Morus notabilis
Cannabis sativa
Silene vulgaris
Beta vulgaris
Spinacia oleracea
Chenopodium quinoa
Sesuvium portulacastrum
Nepenthes ventricosa
Fallopia multiflora
Lophophytum mirabile
Eriobotrya japonica
Malus domestica
Prunus avium
Bombax ceiba
Hibiscus cannabinus
Gossypium trilobum
Corchorus olitorius
Ilex pubescens
Rhazya stricta
Asclepias syriaca
Cynanchum wilfordii
Castilleja paramensis
Ajuga reptans
Salvia miltiorrhiza
Utricularia reniformis
Erythranthe guttata
Haberlea rhodopensis
Dorcoceras hygrometricum
Chionanthus rupicola
Ligustrum quihoui
Hesperelaea palmeri
Olea europaea
Cuscuta gronovii
Cuscuta campestris
Cuscuta australis
Ipomoea nil
Nicotiana tabacum
Hyoscyamus niger
Physochlaina orientalis
Solanum tuberosum
Capsicum annuum
Vaccinium macrocarpon
Monotropa hypopitys
Helianthus annuus
Diplostephium hartwegii
Paraprenanthes diversifolia
Lactuca serriola
Chrysanthemum boreale
Daucus carota
Codonopsis lanceolata
Platycodon grandiflorus
Citrus sinensis
Spondias tuberosa
Xanthoceras sorbifolium
Ziziphus jujuba
Peganum harmala
Epirixanthes elongata
Lagerstroemia indica
Eucalyptus grandis
Glycine soja
Phaseolus vulgaris
Vigna radiata
Millettia pinnata
Bupleurum falcatum
Vicia faba
Medicago truncatula
Lotus japonicus
Sophora flavescens
Ammopiptanthus nanus
Styphnolobium japonicum
Castanospermum australe
Cercis canadensis
Tamarindus indica
Haematoxylum brasiletto
Senna tora
Leucaena trichandra
Libidibia coriaria
Acacia ligulata
0.05

cox2

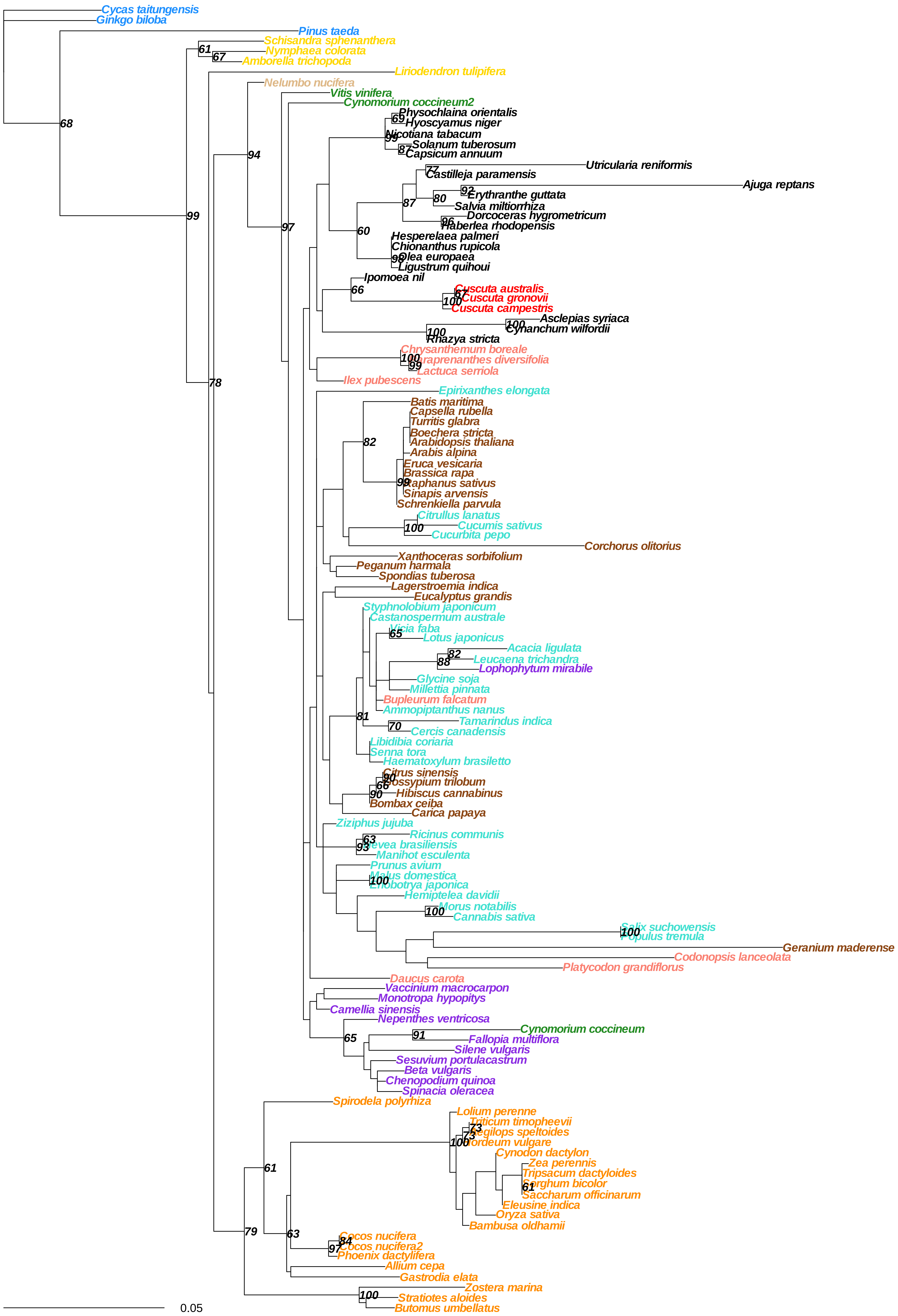

Cycas taitungensis
Ginkgo biloba
Pinus taeda
Schisandra sphenanthera
Nymphaea colorata
Amborella trichopoda
Liriodendron tulipifera
Nelumbo nucifera
Vitis vinifera
Cynomorium coccineum2
Physochlaina orientalis
Hyoscyamus niger
Nicotiana tabacum
Solanum tuberosum
Capsicum annuum
Utricularia reniformis
Castilleja paramensis
Ajuga reptans
Erythranthe guttata
Salvia miltiorrhiza
Dorcoceras hygrometricum
Haberlea rhodopensis
Hesperelaea palmeri
Chionanthus rupicola
Olea europaea
Ligustrum quihoui
Ipomoea nil
Cuscuta australis
Cuscuta gronovii
Cuscuta campestris
Asclepias syriaca
Cynanchum wilfordii
Rhazya stricta
Chrysanthemum boreale
Paraprenanthes diversifolia
Lactuca serriola
Ilex pubescens
Epirixanthes elongata
Batis maritima
Capsella rubella
Turritis glabra
Boechera stricta
Arabidopsis thaliana
Arabis alpina
Eruca vesicaria
Brassica rapa
Raphanus sativus
Sinapis arvensis
Schrenkiella parvula
Citrullus lanatus
Cucumis sativus
Cucurbita pepo
Corchorus olitorius
Xanthoceras sorbifolium
Peganum harmala
Spondias tuberosa
Lagerstroemia indica
Eucalyptus grandis
Styphnolobium japonicum
Castanospermum australe
Vicia faba
Lotus japonicus
Acacia ligulata
Leucaena trichandra
Lophophytum mirabile
Glycine soja
Millettia pinnata
Bupleurum falcatum
Ammopiptanthus nanus
Tamarindus indica
Cercis canadensis
Libidibia coriaria
Senna tora
Haematoxylum brasiletto
Citrus sinensis
Gossypium trilobum
Hibiscus cannabinus
Bombax ceiba
Carica papaya
Ziziphus jujuba
Ricinus communis
Hevea brasiliensis
Manihot esculenta
Prunus avium
Malus domestica
Eriobotrya japonica
Hemiptelea davidii
Morus notabilis
Cannabis sativa
Salix suchowensis
Populus tremula
Geranium maderense
Codonopsis lanceolata
Platycodon grandiflorus
Daucus carota
Vaccinium macrocarpon
Monotropa hypopitys
Camellia sinensis
Nepenthes ventricosa
Cynomorium coccineum
Fallopia multiflora
Silene vulgaris
Sesuvium portulacastrum
Beta vulgaris
Chenopodium quinoa
Spinacia oleracea
Spirodela polyrhiza
Lolium perenne
Triticum timopheevii
Aegilops speltoides
Hordeum vulgare
Cynodon dactylon
Zea perennis
Tripsacum dactyloides
Sorghum bicolor
Saccharum officinarum
Eleusine indica
Oryza sativa
Bambusa oldhamii
Cocos nucifera
Cocos nucifera2
Phoenix dactylifera
Allium cepa
Gastrodia elata
Zostera marina
Stratiotes aloides
Butomus umbellatus
0.05

**cox3**

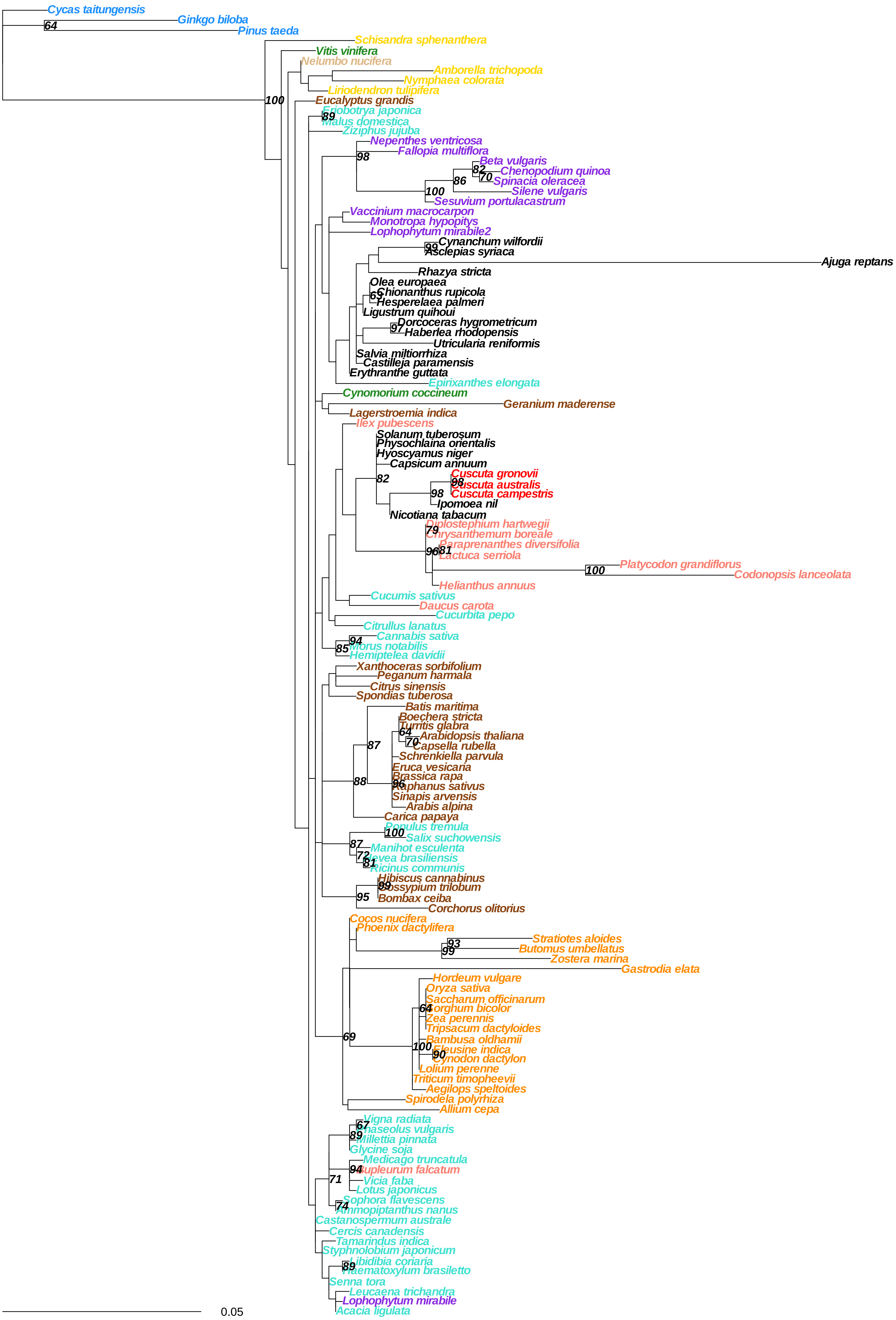
Cycas taitungensis
Ginkgo biloba
Pinus taeda
Schisandra sphenanthera
Vitis vinifera
Nelumbo nucifera
Amborella trichopoda
Nymphaea colorata
Liriodendron tulipifera
Eucalyptus grandis
Eriobotrya japonica
Malus domestica
Ziziphus jujuba
Nepenthes ventricosa
Fallopia multiflora
Beta vulgaris
Chenopodium quinoa
Spinacia oleracea
Silene vulgaris
Sesuvium portulacastrum
Vaccinium macrocarpon
Monotropa hypopitys
Lophophytum mirabile2
Cynanchum wilfordii
Asclepias syriaca
Ajuga reptans
Rhazya stricta
Olea europaea
Chionanthus rupicola
Hesperelaea palmeri
Ligustrum quihoui
Dorcoceras hygrometricum
Haberlea rhodopensis
Utricularia reniformis
Salvia miltiorrhiza
Castilleja paramensis
Erythranthe guttata
Epirixanthes elongata
Cynomorium coccineum
Geranium maderense
Lagerstroemia indica
Ilex pubescens
Solanum tuberosum
Physochlaina orientalis
Hyoscyamus niger
Capsicum annuum
Cuscuta gronovii
Cuscuta australis
Cuscuta campestris
Ipomoea nil
Nicotiana tabacum
Diplostephium hartwegii
Chrysanthemum boreale
Paraprenanthes diversifolia
Lactuca serriola
Platycodon grandiflorus
Codonopsis lanceolata
Helianthus annuus
Cucumis sativus
Daucus carota
Cucurbita pepo
Citrullus lanatus
Cannabis sativa
Morus notabilis
Hemiptelea davidii
Xanthoceras sorbifolium
Peganum harmala
Citrus sinensis
Spondias tuberosa
Batis maritima
Boechera stricta
Turritis glabra
Arabidopsis thaliana
Capsella rubella
Schrenkiella parvula
Eruca vesicaria
Brassica rapa
Raphanus sativus
Sinapis arvensis
Arabis alpina
Carica papaya
Populus tremula
Salix suchowensis
Manihot esculenta
Hevea brasiliensis
Ricinus communis
Hibiscus cannabinus
Gossypium trilobum
Bombax ceiba
Corchorus olitorius
Cocos nucifera
Phoenix dactylifera
Stratiotes aloides
Butomus umbellatus
Zostera marina
Gastrodia elata
Hordeum vulgare
Oryza sativa
Saccharum officinarum
Sorghum bicolor
Zea perennis
Tripsacum dactyloides
Bambusa oldhamii
Eleusine indica
Cynodon dactylon
Lolium perenne
Triticum timopheevii
Aegilops speltoides
Spirodela polyrhiza
Allium cepa
Vigna radiata
Phaseolus vulgaris
Millettia pinnata
Glycine soja
Medicago truncatula
Bupleurum falcatum
Vicia faba
Lotus japonicus
Sophora flavescens
Ammopiptanthus nanus
Castanospermum australe
Cercis canadensis
Tamarindus indica
Styphnolobium japonicum
Libidibia coriaria
Haematoxylum brasiletto
Senna tora
Leucaena trichandra
Lophophytum mirabile
Acacia ligulata
0.05

### matr

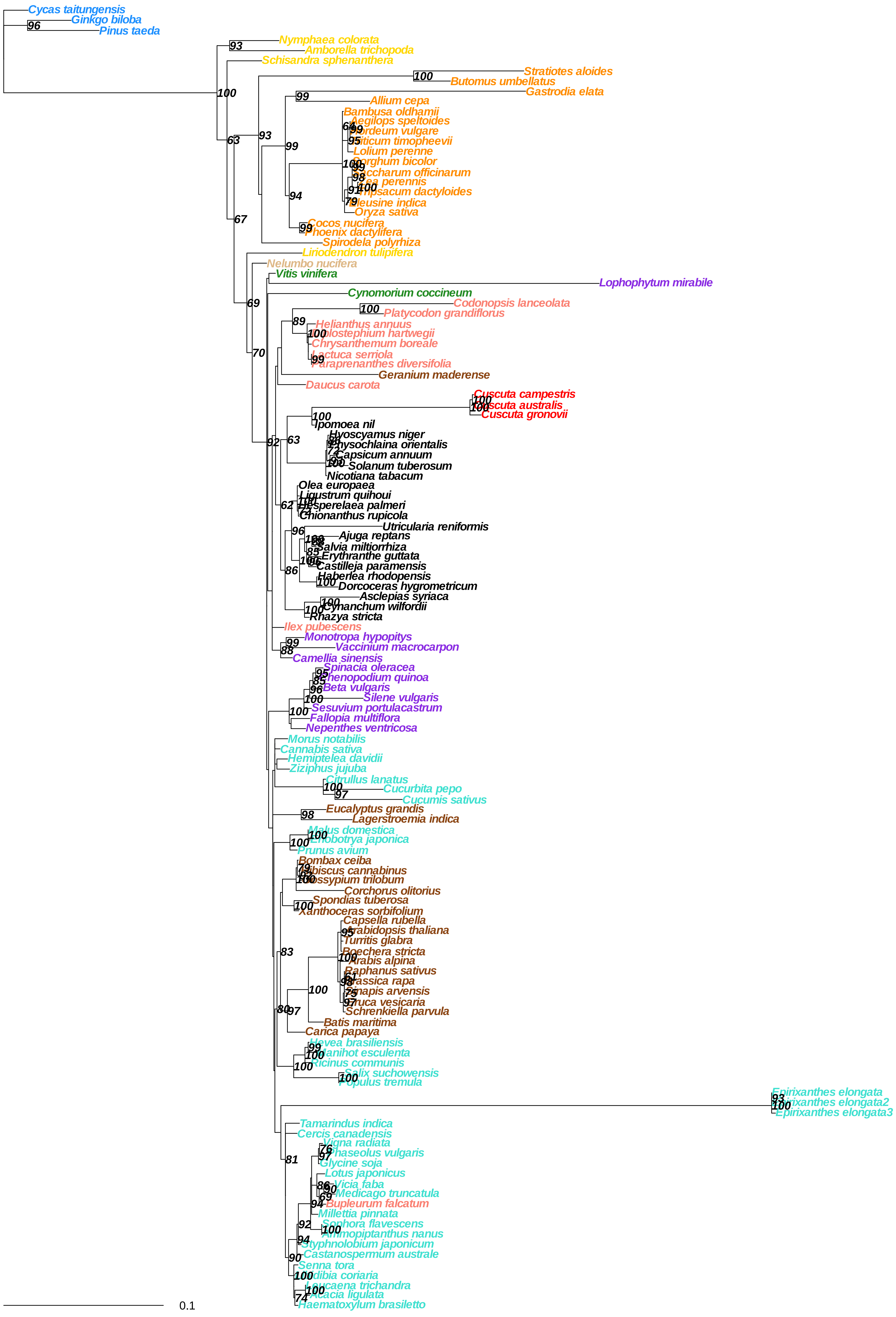
Cycas taitungensis
Ginkgo biloba
Pinus taeda
Nymphaea colorata
Amborella trichopoda
Schisandra sphenanthera
Stratiotes aloides
Butomus umbellatus
Gastrodia elata
Allium cepa
Bambusa oldhamii
Aegilops speltoides
ordeum vulgare
riticum timopheevii
Lolium perenne
Sorghum bicolor
ccharum officinarum
ea perennis
ipsacum dactyloides
leusine indica
Oryza sativa
Cocos nucifera
hoenix dactylifera
Spirodela polyrhiza
Liriodendron tulipifera
Nelumbo nucifera
Vitis vinifera
Lophophytum mirabile
Cynomorium coccineum
Codonopsis lanceolata
Platycodon grandiflorus
Helianthus annuus
olostephium hartwegii
Chrysanthemum boreale
ctuca serriola
araprenanthes diversifolia
Geranium maderense
Daucus carota
Cuscuta campestris
scuta australis
Cuscuta gronovii
Ipomoea nil
Hyoscyamus niger
hysochlaina orientalis
Capsicum annuum
Solanum tuberosum
Nicotiana tabacum
Olea europaea
Ligustrum quihoui
sperelaea palmeri
Chionanthus rupicola
Utricularia reniformis
Ajuga reptans
Salvia miltiorrhiza
Erythranthe guttata
Castilleja paramensis
Haberlea rhodopensis
Dorcoceras hygrometricum
Asclepias syriaca
Cynanchum wilfordii
Rhazya stricta
Ilex pubescens
Monotropa hypopitys
Vaccinium macrocarpon
Camellia sinensis
Spinacia oleracea
henopodium quinoa
Beta vulgaris
Silene vulgaris
Sesuvium portulacastrum
Fallopia multiflora
Nepenthes ventricosa
Morus notabilis
Cannabis sativa
Hemiptelea davidii
Ziziphus jujuba
Citrullus lanatus
Cucurbita pepo
Cucumis sativus
Eucalyptus grandis
Lagerstroemia indica
Malus domestica
obotrya japonica
Prunus avium
Bombax ceiba
biscus cannabinus
ossypium trilobum
Corchorus olitorius
Spondias tuberosa
Xanthoceras sorbifolium
Capsella rubella
rabidopsis thaliana
Turritis glabra
Boechera stricta
rabis alpina
Raphanus sativus
assica rapa
napis arvensis
ruca vesicaria
Schrenkiella parvula
Batis maritima
Carica papaya
Hevea brasiliensis
anihot esculenta
Ricinus communis
Salix suchowensis
opulus tremula
Epirixanthes elongata
rixanthes elongata2
Epirixanthes elongata3
Tamarindus indica
Cercis canadensis
Vigna radiata
haseolus vulgaris
Glycine soja
Lotus japonicus
Vicia faba
Medicago truncatula
Bupleurum falcatum
Millettia pinnata
Sophora flavescens
mmopiptanthus nanus
Styphnolobium japonicum
Castanospermum australe
Senna tora
dibia coriaria
Leucaena trichandra
Acacia ligulata
Haematoxylum brasiletto
0.1

**mttb**

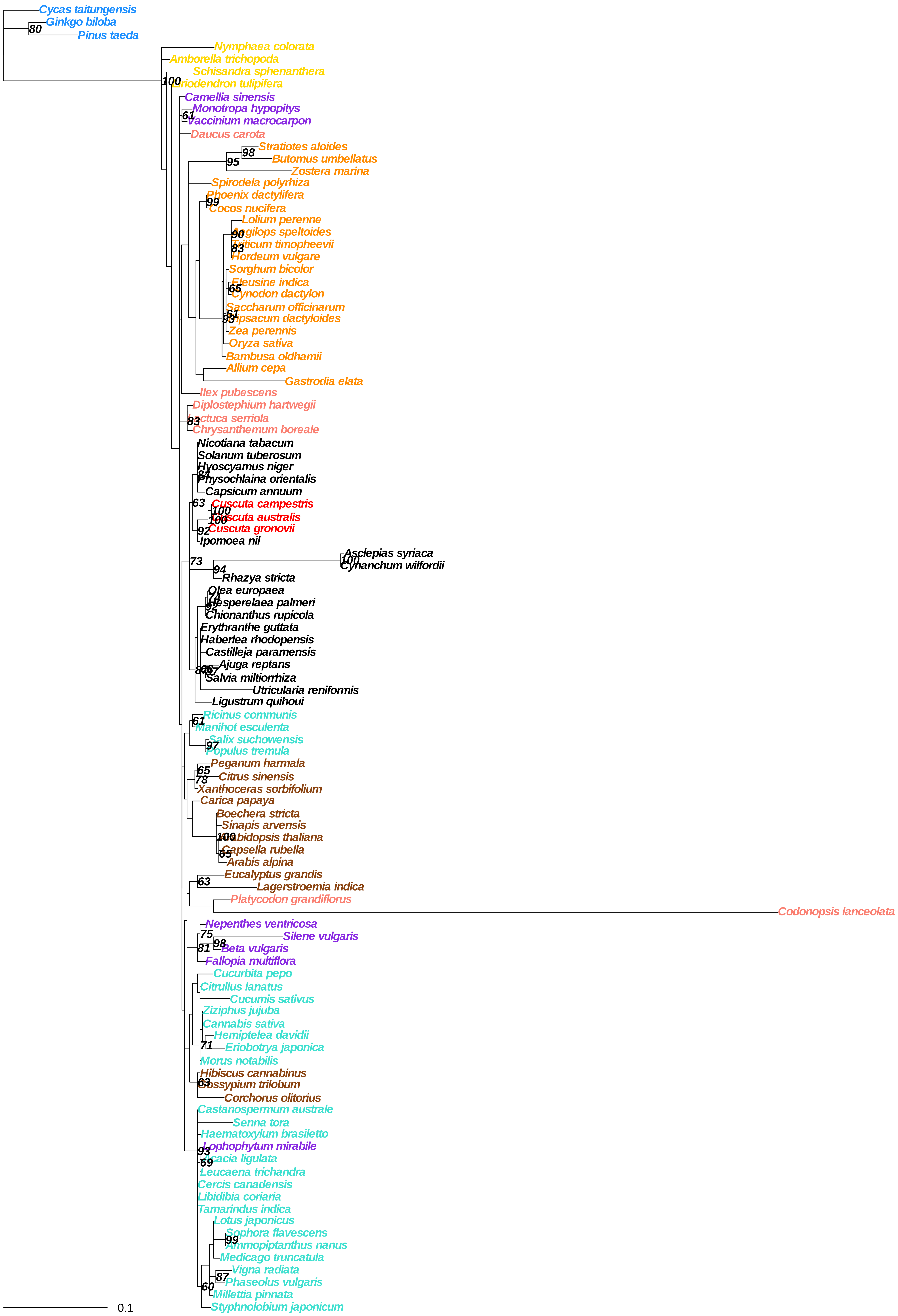
Cycas taitungensis
Ginkgo biloba
Pinus taeda
Nymphaea colorata
Amborella trichopoda
Schisandra sphenanthera
Liriodendron tulipifera
Camellia sinensis
Monotropa hypopitys
Vaccinium macrocarpon
Daucus carota
Stratiotes aloides
Butomus umbellatus
Zostera marina
Spirodela polyrhiza
Phoenix dactylifera
Cocos nucifera
Lolium perenne
Aegilops speltoides
Triticum timopheevii
Hordeum vulgare
Sorghum bicolor
Eleusine indica
Cynodon dactylon
Saccharum officinarum
Tripsacum dactyloides
Zea perennis
Oryza sativa
Bambusa oldhamii
Allium cepa
Gastrodia elata
Ilex pubescens
Diplostephium hartwegii
Lactuca serriola
Chrysanthemum boreale
Nicotiana tabacum
Solanum tuberosum
Hyoscyamus niger
Physochlaina orientalis
Capsicum annuum
Cuscuta campestris
Cuscuta australis
Cuscuta gronovii
Ipomoea nil
Asclepias syriaca
Cynanchum wilfordii
Rhazya stricta
Olea europaea
Hesperelaea palmeri
Chionanthus rupicola
Erythranthe guttata
Haberlea rhodopensis
Castilleja paramensis
Ajuga reptans
Salvia miltiorrhiza
Utricularia reniformis
Ligustrum quihoui
Ricinus communis
Manihot esculenta
Salix suchowensis
Populus tremula
Peganum harmala
Citrus sinensis
Xanthoceras sorbifolium
Carica papaya
Boechera stricta
Sinapis arvensis
Arabidopsis thaliana
Capsella rubella
Arabis alpina
Eucalyptus grandis
Lagerstroemia indica
Platycodon grandiflorus
Codonopsis lanceolata
Nepenthes ventricosa
Silene vulgaris
Beta vulgaris
Fallopia multiflora
Cucurbita pepo
Citrullus lanatus
Cucumis sativus
Ziziphus jujuba
Cannabis sativa
Hemiptelea davidii
Eriobotrya japonica
Morus notabilis
Hibiscus cannabinus
Gossypium trilobum
Corchorus olitorius
Castanospermum australe
Senna tora
Haematoxylum brasiletto
Lophophytum mirabile
Acacia ligulata
Leucaena trichandra
Cercis canadensis
Libidibia coriaria
Tamarindus indica
Lotus japonicus
Sophora flavescens
Ammopiptanthus nanus
Medicago truncatula
Vigna radiata
Phaseolus vulgaris
Millettia pinnata
Styphnolobium japonicum
0.1

### nad1

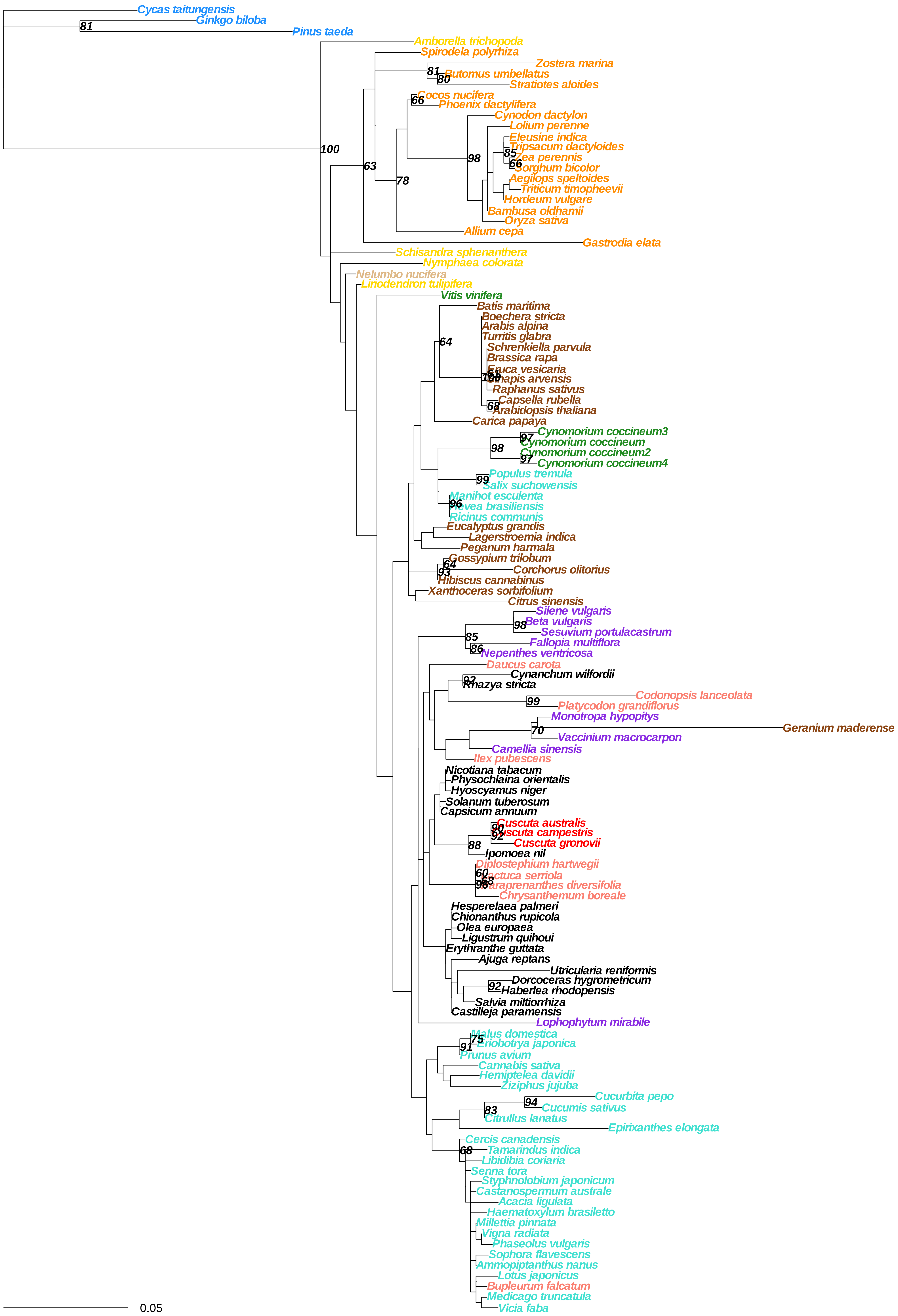

Cycas taitungensis
Ginkgo biloba
Pinus taeda
Amborella trichopoda
Spirodela polyrhiza
Zostera marina
Butomus umbellatus
Stratiotes aloides
Cocos nucifera
Phoenix dactylifera
Cynodon dactylon
Lolium perenne
Eleusine indica
Tripsacum dactyloides
Zea perennis
Sorghum bicolor
Aegilops speltoides
Triticum timopheevii
Hordeum vulgare
Bambusa oldhamii
Oryza sativa
Allium cepa
Gastrodia elata
Schisandra sphenanthera
Nymphaea colorata
Nelumbo nucifera
Liriodendron tulipifera
Vitis vinifera
Batis maritima
Boechera stricta
Arabis alpina
Turritis glabra
Schrenkiella parvula
Brassica rapa
Raphanus sativus
Capsella rubella
Arabidopsis thaliana
Carica papaya
Cynomorium coccineum3
Cynomorium coccineum
Cynomorium coccineum2
Cynomorium coccineum4
Populus tremula
Salix suchowensis
Manihot esculenta
Ricinus communis
Eucalyptus grandis
Lagerstroemia indica
Peganum harmala
Gossypium trilobum
Corchorus olitorius
Hibiscus cannabinus
Xanthoceras sorbifolium
Citrus sinensis
Silene vulgaris
Beta vulgaris
Sesuvium portulacastrum
Fallopia multiflora
Nepenthes ventricosa
Daucus carota
Cynanchum wilfordii
Rhazya stricta
Codonopsis lanceolata
Platycodon grandiflorus
Monotropa hypopitys
Geranium maderense
Vaccinium macrocarpon
Camellia sinensis
Ilex pubescens
Nicotiana tabacum
Physochlaina orientalis
Hyoscyamus niger
Solanum tuberosum
Capsicum annuum
Cuscuta australis
Cuscuta gronovii
Ipomoea nil
Diplostephium hartwegii
Chrysanthemum boreale
Hesperelaea palmeri
Chionanthus rupicola
Olea europaea
Ligustrum quihoui
Erythranthe guttata
Ajuga reptans
Utricularia reniformis
Dorcoceras hygrometricum
Haberlea rhodopensis
Salvia miltiorrhiza
Castilleja paramensis
Lophophytum mirabile
Malus domestica
Eriobotrya japonica
Prunus avium
Cannabis sativa
Hemiptelea davidii
Ziziphus jujuba
Cucurbita pepo
Cucumis sativus
Citrullus lanatus
Epirixanthes elongata
Cercis canadensis
Tamarindus indica
Libidibia coriaria
Senna tora
Styphnolobium japonicum
Castanospermum australe
Acacia ligulata
Haematoxylum brasiletto
Millettia pinnata
Vigna radiata
Phaseolus vulgaris
Sophora flavescens
Ammopiptanthus nanus
Lotus japonicus
Bupleurum falcatum
Medicago truncatula
Vicia faba
0.05

**nad2**

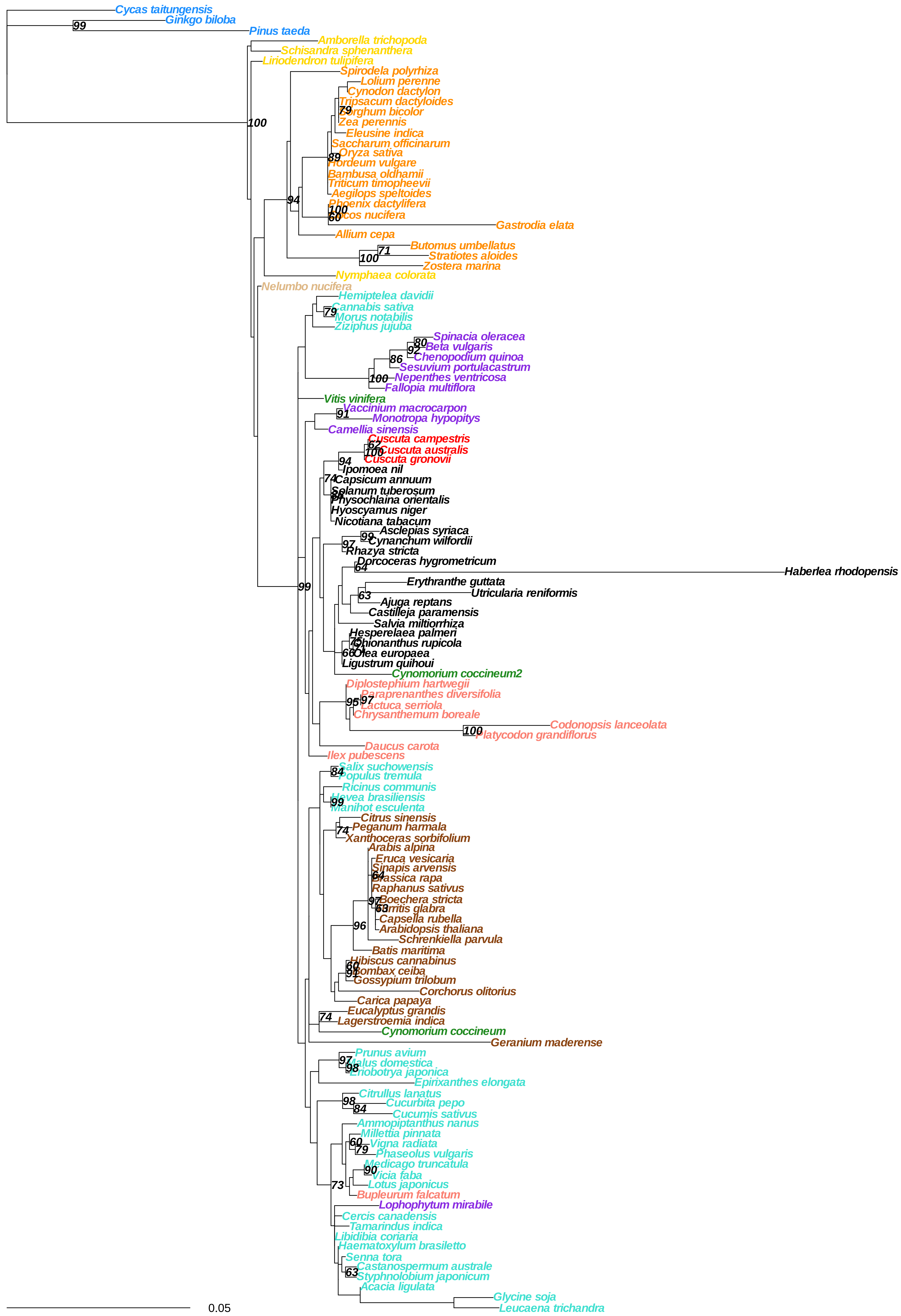
Cycas taitungensis
Ginkgo biloba
Pinus taeda
Amborella trichopoda
Schisandra sphenanthera
Liriodendron tulipifera
Spirodela polyrhiza
Lolium perenne
Cynodon dactylon
Tripsacum dactyloides
Sorghum bicolor
Zea perennis
Eleusine indica
Saccharum officinarum
Oryza sativa
Hordeum vulgare
Bambusa oldhamii
Triticum timopheevii
Aegilops speltoides
Phoenix dactylifera
Cocos nucifera
Gastrodia elata
Allium cepa
Butomus umbellatus
Stratiotes aloides
Zostera marina
Nymphaea colorata
Nelumbo nucifera
Hemiptelea davidii
Cannabis sativa
Morus notabilis
Ziziphus jujuba
Spinacia oleracea
Beta vulgaris
Chenopodium quinoa
Sesuvium portulacastrum
Nepenthes ventricosa
Fallopia multiflora
Vitis vinifera
Vaccinium macrocarpon
Monotropa hypopitys
Camellia sinensis
Cuscuta campestris
Cuscuta australis
Cuscuta gronovii
Ipomoea nil
Capsicum annuum
Solanum tuberosum
Physochlaina orientalis
Hyoscyamus niger
Nicotiana tabacum
Asclepias syriaca
Cynanchum wilfordii
Rhazya stricta
Dorcoceras hygrometricum
Haberlea rhodopensis
Erythranthe guttata
Utricularia reniformis
Ajuga reptans
Castilleja paramensis
Salvia miltiorrhiza
Hesperelaea palmeri
Chionanthus rupicola
Olea europaea
Ligustrum quihoui
Cynomorium coccineum2
Diplostephium hartwegii
Paraprenanthes diversifolia
Lactuca serriola
Chrysanthemum boreale
Codonopsis lanceolata
Platycodon grandiflorus
Daucus carota
Ilex pubescens
Salix suchowensis
Populus tremula
Ricinus communis
Hevea brasiliensis
Manihot esculenta
Citrus sinensis
Peganum harmala
Xanthoceras sorbifolium
Arabis alpina
Eruca vesicaria
Sinapis arvensis
Brassica rapa
Raphanus sativus
Boechera stricta
Turritis glabra
Capsella rubella
Arabidopsis thaliana
Schrenkiella parvula
Batis maritima
Hibiscus cannabinus
Bombax ceiba
Gossypium trilobum
Corchorus olitorius
Carica papaya
Eucalyptus grandis
Lagerstroemia indica
Cynomorium coccineum
Geranium maderense
Prunus avium
Malus domestica
Eriobotrya japonica
Epirixanthes elongata
Citrullus lanatus
Cucurbita pepo
Cucumis sativus
Ammopiptanthus nanus
Millettia pinnata
Vigna radiata
Phaseolus vulgaris
Medicago truncatula
Vicia faba
Lotus japonicus
Bupleurum falcatum
Lophophytum mirabile
Cercis canadensis
Tamarindus indica
Libidibia coriaria
Haematoxylum brasiletto
Senna tora
Castanospermum australe
Styphnolobium japonicum
Acacia ligulata
Glycine soja
Leucaena trichandra
0.05

**nad3**

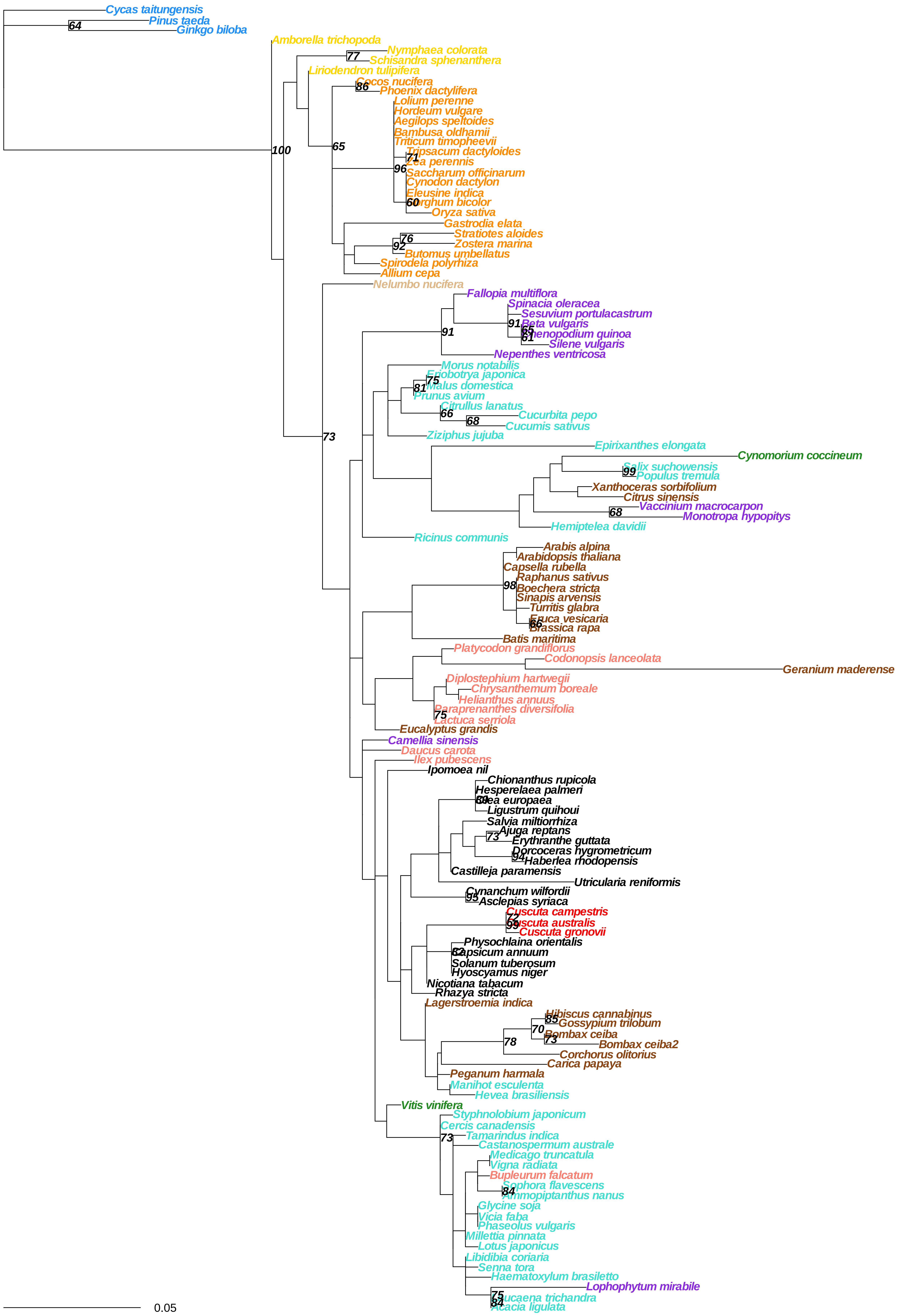
Cycas taitungensis
Pinus taeda
64
Ginkgo biloba
Amborella trichopoda
Nymphaea colorata
77
Schisandra sphenanthera
Liriodendron tulipifera
Cocos nucifera
86
Phoenix dactylifera
Lolium perenne
Hordeum vulgare
Aegilops speltoides
Bambusa oldhamii
Triticum timopheevii
Tripsacum dactyloides
71
100
65
96
Saccharum officinarum
Cynodon dactylon
Eleusine indica
60
Oryza sativa
Gastrodia elata
Stratiotes aloides
76
92
Zostera marina
Butomus umbellatus
Spirodela polyrhiza
Allium cepa
Nelumbo nucifera
Fallopia multiflora
Spinacia oleracea
Sesuvium portulacastrum
91
Beta vulgaris
65
61
Silene vulgaris
Nepenthes ventricosa
Morus notabilis
75
Malus domestica
81
Prunus avium
Citrullus lanatus
66
Cucurbita pepo
68
Cucumis sativus
Ziziphus jujuba
73
Epirixanthes elongata
Cynomorium coccineum
99
Populus tremula
Xanthoceras sorbifolium
Citrus sinensis
Vaccinium macrocarpon
68
Monotropa hypopitys
Hemiptelea davidii
Ricinus communis
Arabis alpina
Arabidopsis thaliana
Capsella rubella
Raphanus sativus
98
Boechera stricta
Sinapis arvensis
Turritis glabra
66
Batis maritima
Platycodon grandiflorus
Codonopsis lanceolata
Geranium maderense
Diplostephium hartwegii
Chrysanthemum boreale
Helianthus annuus
75
Eucalyptus grandis
Camellia sinensis
Daucus carota
Ilex pubescens
Ipomoea nil
Chionanthus rupicola
Hesperelaea palmeri
Ligustrum quihoui
Salvia miltiorrhiza
Ajuga reptans
73
Erythranthe guttata
Dorcoceras hygrometricum
94
Haberlea rhodopensis
Castilleja paramensis
Utricularia reniformis
Cynanchum wilfordii
95
Asclepias syriaca
Cuscuta campestris
99
Cuscuta gronovii
Physochlaina orientalis
Solanum tuberosum
Hyoscyamus niger
Nicotiana tabacum
Rhazya stricta
Lagerstroemia indica
Hibiscus cannabinus
85
Gossypium trilobum
70
Bombax ceiba
73
Bombax ceiba2
78
Corchorus olitorius
Carica papaya
Peganum harmala
Manihot esculenta
Hevea brasiliensis
Vitis vinifera
Styphnolobium japonicum
Cercis canadensis
Tamarindus indica
73
Castanospermum australe
Medicago truncatula
Vigna radiata
Bupleurum falcatum
Sophora flavescens
84
Glycine soja
Vicia faba
Phaseolus vulgaris
Millettia pinnata
Lotus japonicus
Libidibia coriaria
Senna tora
Haematoxylum brasiletto
Lophophytum mirabile
75
84
Acacia ligulata
0.05

#### nad4

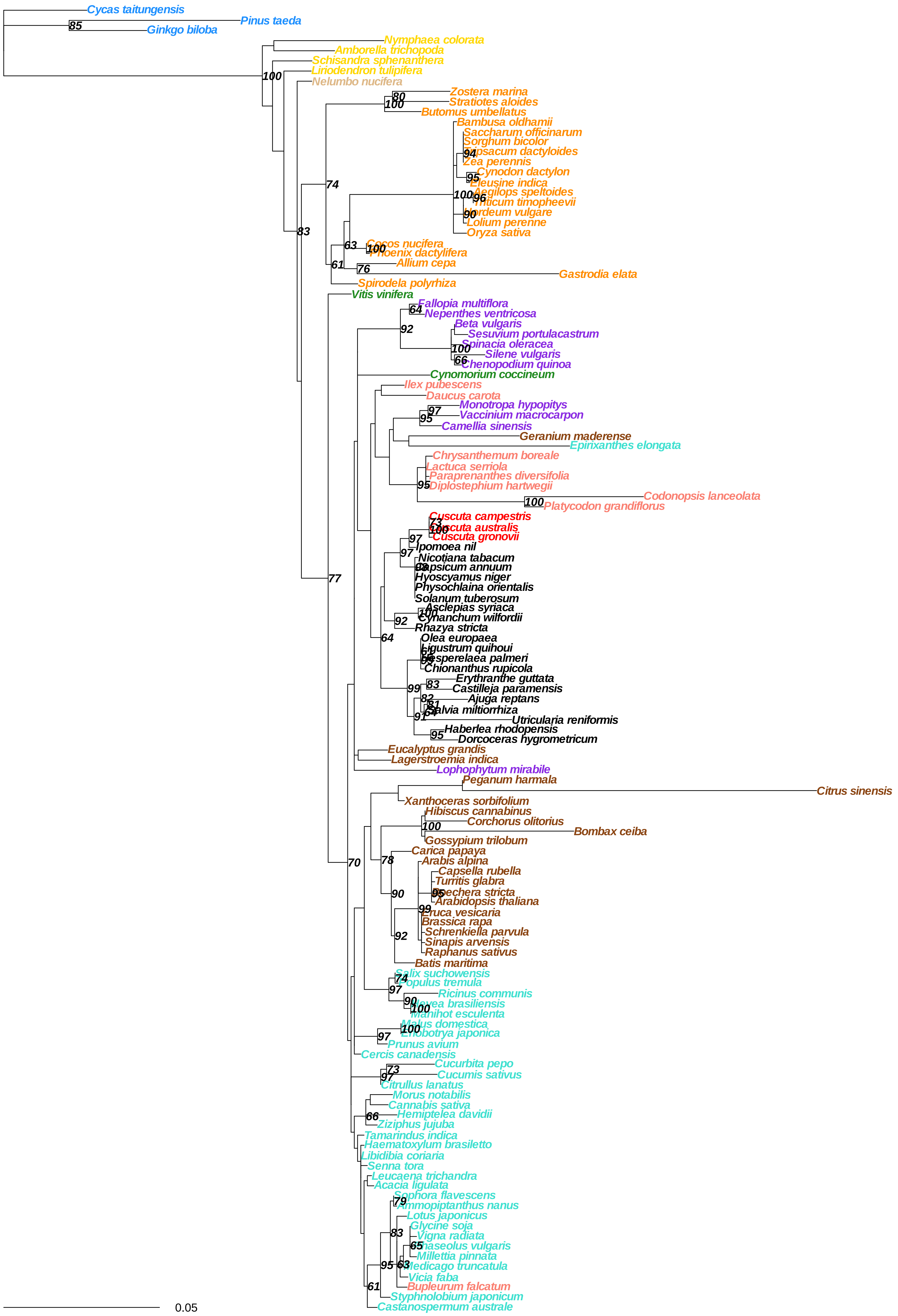

Cycas taitungensis
Pinus taeda
Ginkgo biloba
Nymphaea colorata
Amborella trichopoda
Schisandra sphenanthera
Liriodendron tulipifera
Nelumbo nucifera
Zostera marina
Stratiotes aloides
Butomus umbellatus
Bambusa oldhamii
Saccharum officinarum
Sorghum bicolor
Tripsacum dactyloides
Zea perennis
Cynodon dactylon
Eleusine indica
Aegilops speltoides
Triticum timopheevii
Hordeum vulgare
Lolium perenne
Oryza sativa
Cocos nucifera
Phoenix dactylifera
Allium cepa
Gastrodia elata
Spirodela polyrhiza
Vitis vinifera
Fallopia multiflora
Nepenthes ventricosa
Beta vulgaris
Sesuvium portulacastrum
Spinacia oleracea
Silene vulgaris
Chenopodium quinoa
Cynomorium coccineum
Ilex pubescens
Daucus carota
Monotropa hypopitys
Vaccinium macrocarpon
Camellia sinensis
Geranium maderense
Epirixanthes elongata
Chrysanthemum boreale
Lactuca serriola
Paraprenanthes diversifolia
Diplostephium hartwegii
Codonopsis lanceolata
Platycodon grandiflorus
Cuscuta campestris
Cuscuta australis
Cuscuta gronovii
Ipomoea nil
Nicotiana tabacum
Capsicum annuum
Hyoscyamus niger
Physochlaina orientalis
Solanum tuberosum
Asclepias syriaca
Cynanchum wilfordii
Rhazya stricta
Olea europaea
Ligustrum quihoui
Hesperelaea palmeri
Chionanthus rupicola
Erythranthe guttata
Castilleja paramensis
Ajuga reptans
Salvia miltiorrhiza
Utricularia reniformis
Haberlea rhodopensis
Dorcoceras hygrometricum
Eucalyptus grandis
Lagerstroemia indica
Lophophytum mirabile
Peganum harmala
Citrus sinensis
Xanthoceras sorbifolium
Hibiscus cannabinus
Corchorus olitorius
Bombax ceiba
Gossypium trilobum
Carica papaya
Arabis alpina
Capsella rubella
Turritis glabra
Boechera stricta
Arabidopsis thaliana
Eruca vesicaria
Brassica rapa
Schrenkiella parvula
Sinapis arvensis
Raphanus sativus
Batis maritima
Salix suchowensis
Populus tremula
Ricinus communis
Hevea brasiliensis
Manihot esculenta
Malus domestica
Eriobotrya japonica
Prunus avium
Cercis canadensis
Cucurbita pepo
Cucumis sativus
Citrullus lanatus
Morus notabilis
Cannabis sativa
Hemiptelea davidii
Ziziphus jujuba
Tamarindus indica
Haematoxylum brasiletto
Libidibia coriaria
Senna tora
Leucaena trichandra
Acacia ligulata
Sophora flavescens
Ammopiptanthus nanus
Lotus japonicus
Glycine soja
Vigna radiata
Phaseolus vulgaris
Millettia pinnata
Medicago truncatula
Vicia faba
Bupleurum falcatum
Styphnolobium japonicum
Castanospermum australe
0.05

### nad4l

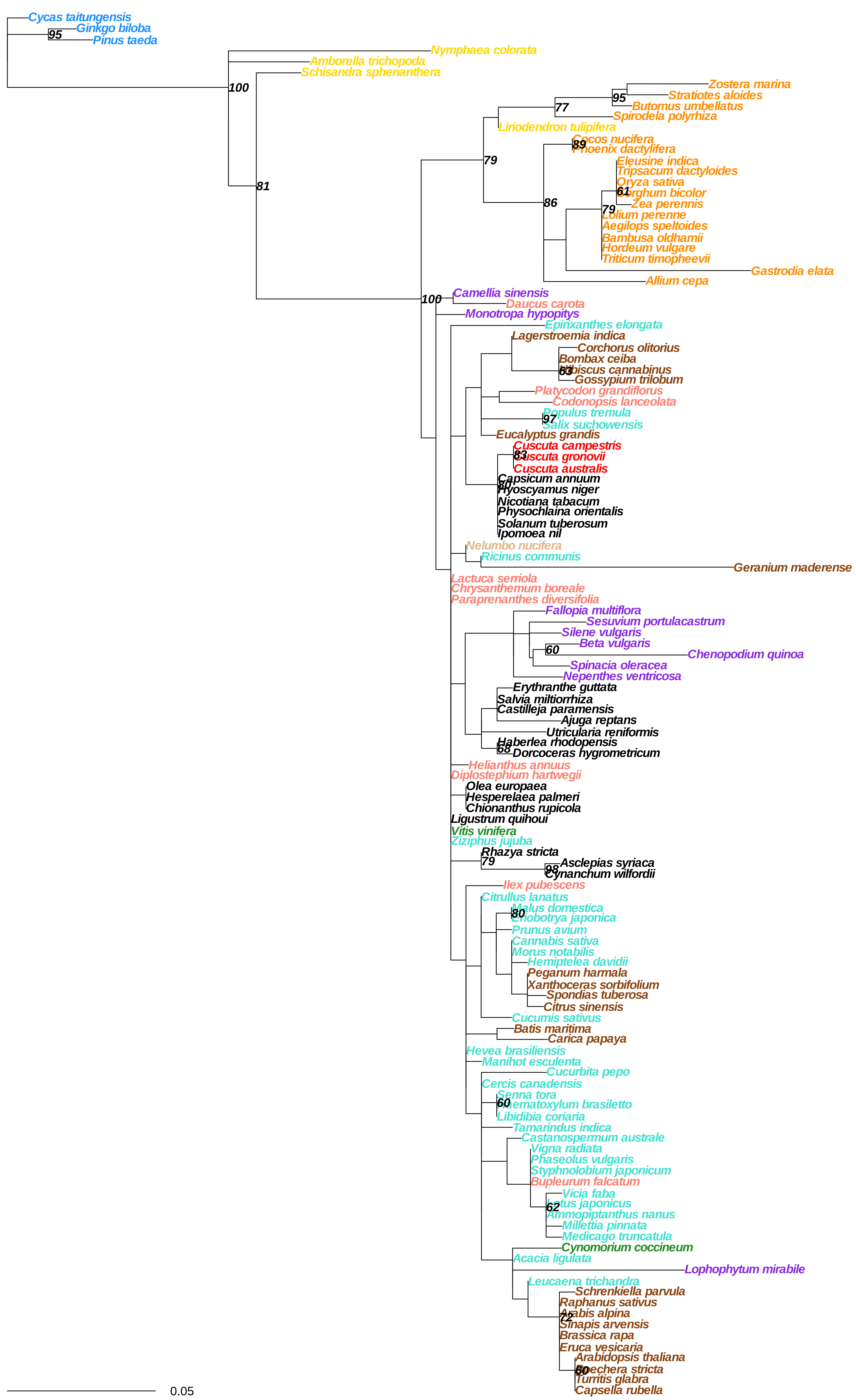
Cycas taitungensis
Ginkgo biloba
Pinus taeda
95
Nymphaea colorata
Amborella trichopoda
Schisandra sphenanthera
100
81
Zostera marina
Stratiotes aloides
Butomus umbellatus
Spirodela polyrhiza
95
77
Liriodendron tulipifera
Cocos nucifera
Phoenix dactylifera
89
79
Eleusine indica
Tripsacum dactyloides
Oryza sativa
Sorghum bicolor
61
Zea perennis
Lolium perenne
Aegilops speltoides
Bambusa oldhamii
Hordeum vulgare
Triticum timopheevii
86
79
Gastrodia elata
Allium cepa
Camellia sinensis
Daucus carota
100
Monotropa hypopitys
Epirixanthes elongata
Lagerstroemia indica
Corchorus olitorius
Bombax ceiba
Hibiscus cannabinus
63
Gossypium trilobum
Platycodon grandiflorus
Codonopsis lanceolata
Populus tremula
Salix suchowensis
97
Eucalyptus grandis
Cuscuta campestris
Cuscuta gronovii
83
Cuscuta australis
Capsicum annuum
80
Hyoscyamus niger
Nicotiana tabacum
Physochlaina orientalis
Solanum tuberosum
Ipomoea nil
Nelumbo nucifera
Ricinus communis
Geranium maderense
Lactuca serriola
Chrysanthemum boreale
Paraprenanthes diversifolia
Fallopia multiflora
Sesuvium portulacastrum
Silene vulgaris
Beta vulgaris
60
Chenopodium quinoa
Spinacia oleracea
Nepenthes ventricosa
Erythranthe guttata
Salvia miltiorrhiza
Castilleja paramensis
Ajuga reptans
Utricularia reniformis
Haberlea rhodopensis
68
Dorcoceras hygrometricum
Helianthus annuus
Diplostephium hartwegii
Olea europaea
Hesperelaea palmeri
Chionanthus rupicola
Ligustrum quihoui
Vitis vinifera
Ziziphus jujuba
Rhazya stricta
79
Asclepias syriaca
98
Cynanchum wilfordii
Ilex pubescens
Citrullus lanatus
Malus domestica
80
Eriobotrya japonica
Prunus avium
Cannabis sativa
Morus notabilis
Hemiptelea davidii
Peganum harmala
Xanthoceras sorbifolium
Spondias tuberosa
Citrus sinensis
Cucumis sativus
Batis maritima
Carica papaya
Hevea brasiliensis
Manihot esculenta
Cucurbita pepo
Cercis canadensis
Senna tora
60
Haematoxylum brasiletto
Libidibia coriaria
Tamarindus indica
Castanospermum australe
Vigna radiata
Phaseolus vulgaris
Styphnolobium japonicum
Bupleurum falcatum
Vicia faba
62
Lotus japonicus
Ammopiptanthus nanus
Millettia pinnata
Medicago truncatula
Cynomorium coccineum
Acacia ligulata
Lophophytum mirabile
Leucaena trichandra
Schrenkiella parvula
Raphanus sativus
72
Arabis alpina
Sinapis arvensis
Brassica rapa
Eruca vesicaria
Arabidopsis thaliana
60
Boechera stricta
Turritis glabra
Capsella rubella
0.05

**nad5**

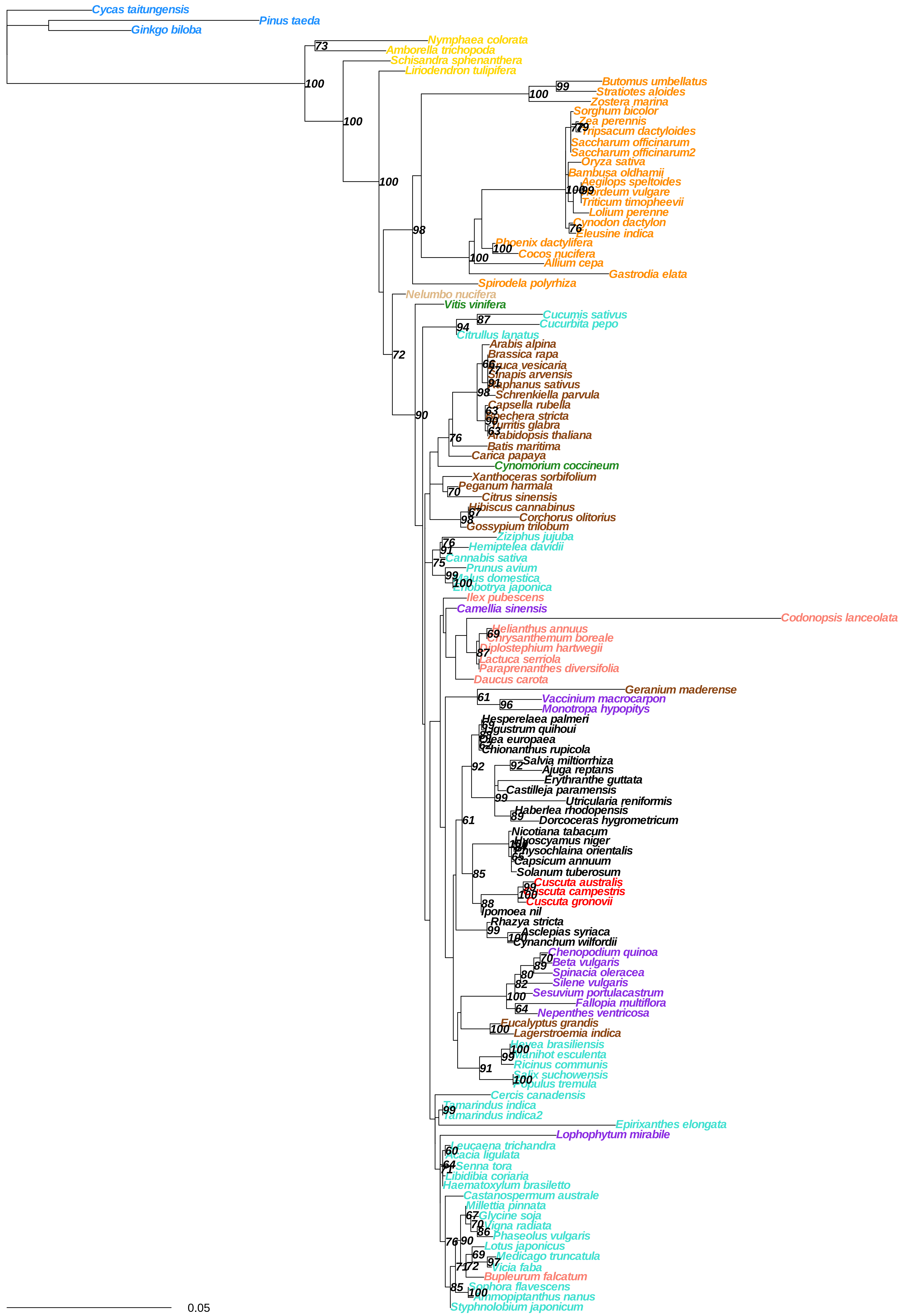

Cycas taitungensis
Pinus taeda
Ginkgo biloba
Nymphaea colorata
Amborella trichopoda
Schisandra sphenanthera
Liriodendron tulipifera
Butomus umbellatus
Stratiotes aloides
Zostera marina
Sorghum bicolor
Zea perennis
Tripsacum dactyloides
Saccharum officinarum
Saccharum officinarum2
Oryza sativa
Bambusa oldhamii
Aegilops speltoides
Hordeum vulgare
Triticum timopheevii
Lolium perenne
Cynodon dactylon
Eleusine indica
Phoenix dactylifera
Cocos nucifera
Allium cepa
Gastrodia elata
Spirodela polyrhiza
Nelumbo nucifera
Vitis vinifera
Cucumis sativus
Cucurbita pepo
Citrullus lanatus
Arabis alpina
Brassica rapa
Eruca vesicaria
Sinapis arvensis
Raphanus sativus
Schrenkiella parvula
Capsella rubella
Boechera stricta
Turritis glabra
Arabidopsis thaliana
Batis maritima
Carica papaya
Cynomorium coccineum
Xanthoceras sorbifolium
Peganum harmala
Citrus sinensis
Hibiscus cannabinus
Corchorus olitorius
Gossypium trilobum
Ziziphus jujuba
Hemiptelea davidii
Cannabis sativa
Prunus avium
Malus domestica
Eriobotrya japonica
Ilex pubescens
Camellia sinensis
Codonopsis lanceolata
Helianthus annuus
Chrysanthemum boreale
Diplostephium hartwegii
Lactuca serriola
Paraprenanthes diversifolia
Daucus carota
Geranium maderense
Vaccinium macrocarpon
Monotropa hypopitys
Hesperelaea palmeri
Ligustrum quihoui
Olea europaea
Chionanthus rupicola
Salvia miltiorrhiza
Ajuga reptans
Erythranthe guttata
Castilleja paramensis
Utricularia reniformis
Haberlea rhodopensis
Dorcoceras hygrometricum
Nicotiana tabacum
Hyoscyamus niger
Physochlaina orientalis
Capsicum annuum
Solanum tuberosum
Cuscuta australis
Cuscuta campestris
Cuscuta gronovii
Ipomoea nil
Rhazya stricta
Asclepias syriaca
Cynanchum wilfordii
Chenopodium quinoa
Beta vulgaris
Spinacia oleracea
Silene vulgaris
Sesuvium portulacastrum
Fallopia multiflora
Nepenthes ventricosa
Eucalyptus grandis
Lagerstroemia indica
Hevea brasiliensis
Manihot esculenta
Ricinus communis
Salix suchowensis
Populus tremula
Cercis canadensis
Tamarindus indica
Tamarindus indica2
Epirixanthes elongata
Lophophytum mirabile
Leucaena trichandra
Acacia ligulata
Senna tora
Libidibia coriaria
Haematoxylum brasiletto
Castanospermum australe
Millettia pinnata
Glycine soja
Vigna radiata
Phaseolus vulgaris
Lotus japonicus
Medicago truncatula
Vicia faba
Bupleurum falcatum
Sophora flavescens
Ammopiptanthus nanus
Styphnolobium japonicum
0.05

**nad6**

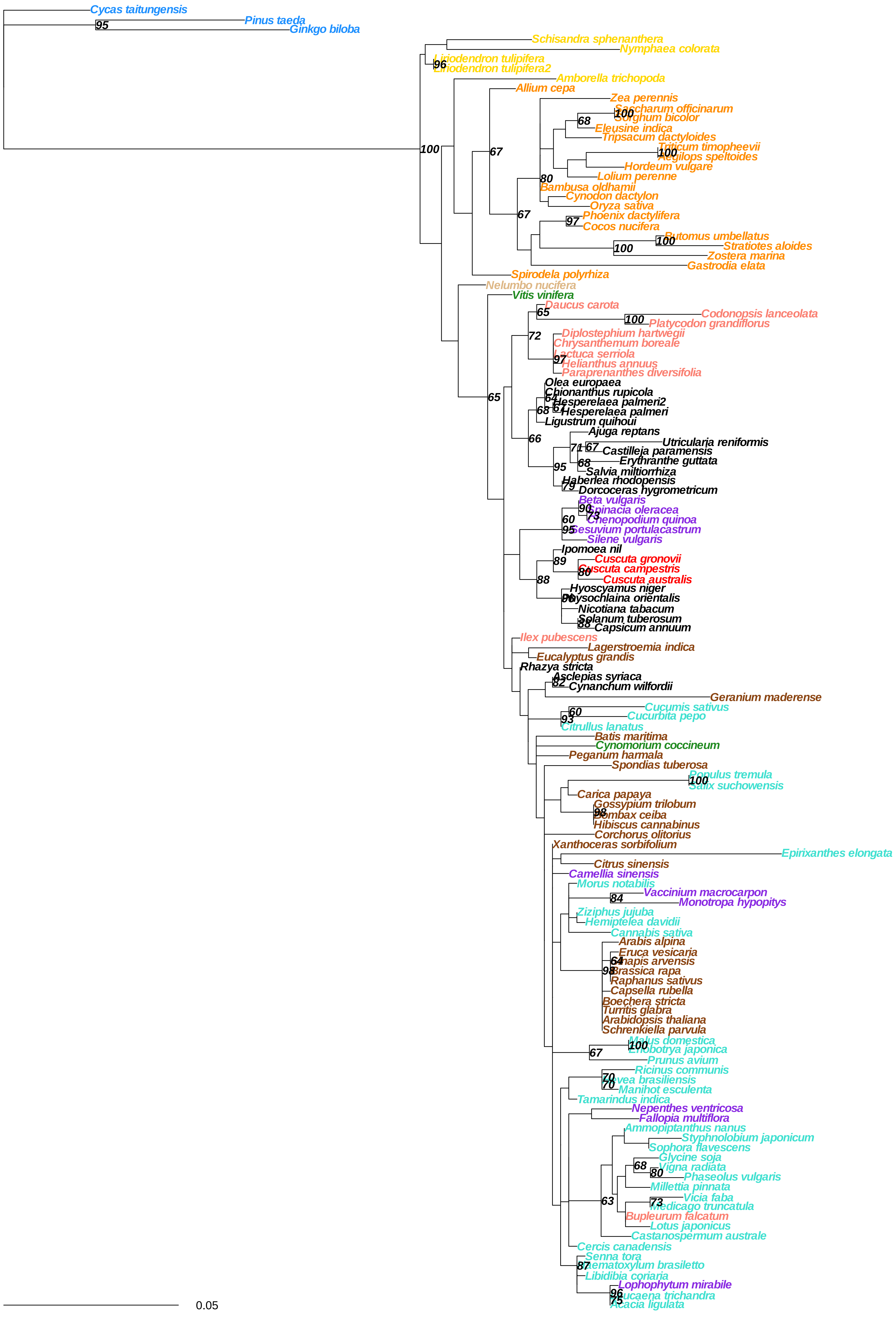
Cycas taitungensis
Pinus taeda
Ginkgo biloba
Schisandra sphenanthera
Nymphaea colorata
Liriodendron tulipifera
Liriodendron tulipifera2
Amborella trichopoda
Allium cepa
Zea perennis
Saccharum officinarum
Sorghum bicolor
Eleusine indica
Tripsacum dactyloides
Triticum timopheevii
Aegilops speltoides
Hordeum vulgare
Lolium perenne
Bambusa oldhamii
Cynodon dactylon
Oryza sativa
Phoenix dactylifera
Cocos nucifera
Butomus umbellatus
Stratiotes aloides
Zostera marina
Gastrodia elata
Spirodela polyrhiza
Nelumbo nucifera
Vitis vinifera
Daucus carota
Codonopsis lanceolata
Platycodon grandiflorus
Diplostephium hartwegii
Chrysanthemum boreale
Lactuca serriola
Helianthus annuus
Paraprenanthes diversifolia
Olea europaea
Chionanthus rupicola
Hesperelaea palmeri2
Hesperelaea palmeri
Ligustrum quihoui
Ajuga reptans
Utricularia reniformis
Castilleja paramensis
Erythranthe guttata
Salvia miltiorrhiza
Haberlea rhodopensis
Dorcoceras hygrometricum
Beta vulgaris
Spinacia oleracea
Chenopodium quinoa
Sesuvium portulacastrum
Silene vulgaris
Ipomoea nil
Cuscuta gronovii
Cuscuta campestris
Cuscuta australis
Hyoscyamus niger
Physochlaina orientalis
Nicotiana tabacum
Solanum tuberosum
Capsicum annuum
Ilex pubescens
Lagerstroemia indica
Eucalyptus grandis
Rhazya stricta
Asclepias syriaca
Cynanchum wilfordii
Geranium maderense
Cucumis sativus
Cucurbita pepo
Citrullus lanatus
Batis maritima
Cynomorium coccineum
Peganum harmala
Spondias tuberosa
Populus tremula
Salix suchowensis
Carica papaya
Gossypium trilobum
Bombax ceiba
Hibiscus cannabinus
Corchorus olitorius
Xanthoceras sorbifolium
Epirixanthes elongata
Citrus sinensis
Camellia sinensis
Morus notabilis
Vaccinium macrocarpon
Monotropa hypopitys
Ziziphus jujuba
Hemiptelea davidii
Cannabis sativa
Arabis alpina
Eruca vesicaria
Sinapis arvensis
Brassica rapa
Raphanus sativus
Capsella rubella
Boechera stricta
Turritis glabra
Arabidopsis thaliana
Schrenkiella parvula
Malus domestica
Eriobotrya japonica
Prunus avium
Ricinus communis
Hevea brasiliensis
Manihot esculenta
Tamarindus indica
Nepenthes ventricosa
Fallopia multiflora
Ammopiptanthus nanus
Styphnolobium japonicum
Sophora flavescens
Glycine soja
Vigna radiata
Phaseolus vulgaris
Millettia pinnata
Vicia faba
Medicago truncatula
Bupleurum falcatum
Lotus japonicus
Castanospermum australe
Cercis canadensis
Senna tora
Haematoxylum brasiletto
Libidibia coriaria
Lophophytum mirabile
Leucaena trichandra
Acacia ligulata
0.05

nad7

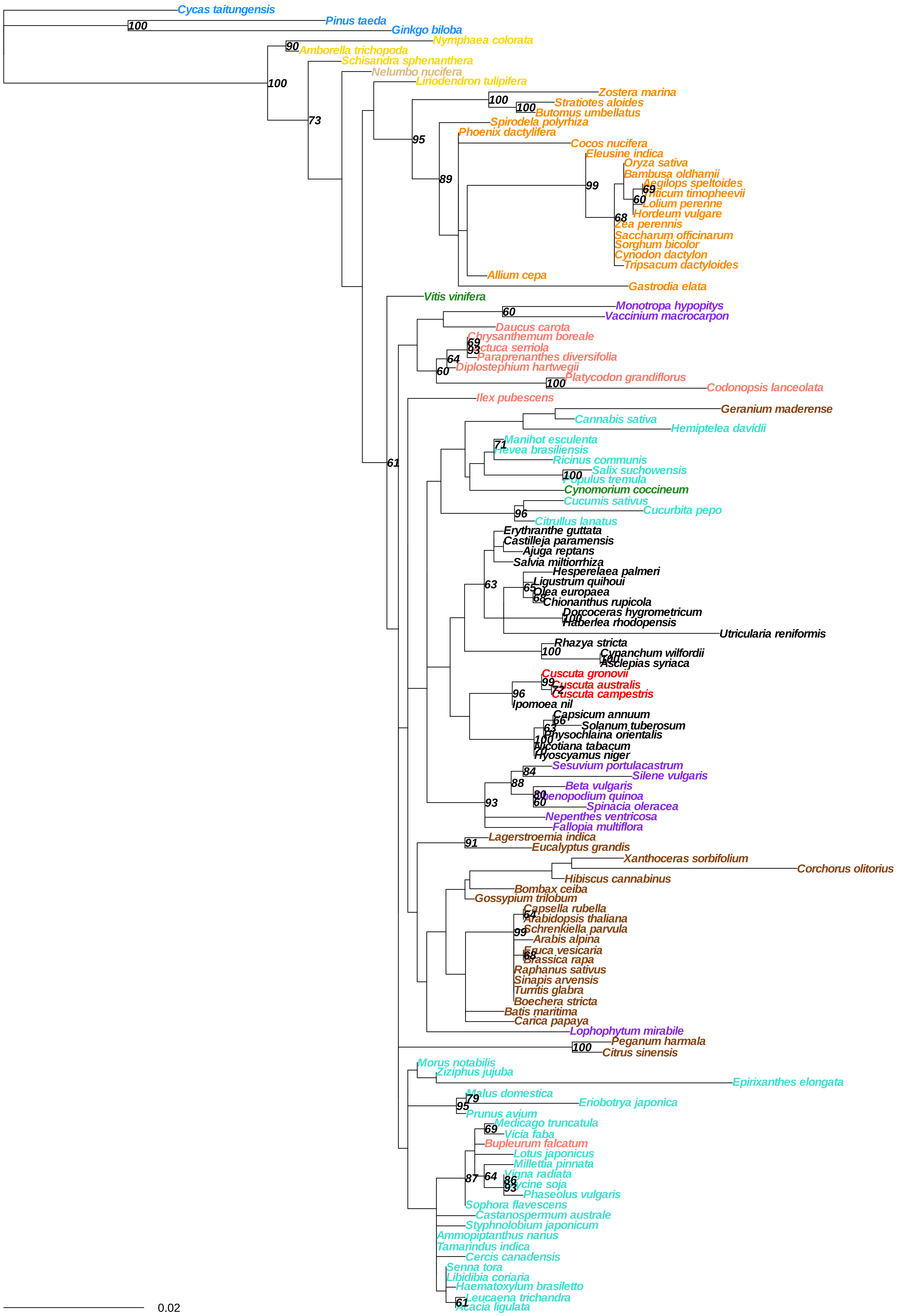

Cycas taitungensis
Pinus taeda
Ginkgo biloba
Nymphaea colorata
Amborella trichopoda
Schisandra sphenanthera
Nelumbo nucifera
Liriodendron tulipifera
Zostera marina
Stratiotes aloides
Butomus umbellatus
Spirodela polyrhiza
Phoenix dactylifera
Cocos nucifera
Eleusine indica
Oryza sativa
Bambusa oldhamii
Aegilops speltoides
Triticum timopheevii
Lolium perenne
Hordeum vulgare
Zea perennis
Saccharum officinarum
Sorghum bicolor
Cynodon dactylon
Tripsacum dactyloides
Allium cepa
Gastrodia elata
Vitis vinifera
Monotropa hypopitys
Vaccinium macrocarpon
Daucus carota
Chrysanthemum boreale
Lactuca serriola
Paraprenanthes diversifolia
Diplostephium hartwegii
Platycodon grandiflorus
Codonopsis lanceolata
Ilex pubescens
Geranium maderense
Cannabis sativa
Hemiptelea davidii
Manihot esculenta
Hevea brasiliensis
Ricinus communis
Salix suchowensis
Populus tremula
Cynomorium coccineum
Cucumis sativus
Cucurbita pepo
Citrullus lanatus
Erythranthe guttata
Castilleja paramensis
Ajuga reptans
Salvia miltiorrhiza
Hesperelaea palmeri
Ligustrum quihoui
Olea europaea
Chionanthus rupicola
Dorcoceras hygrometricum
Haberlea rhodopensis
Utricularia reniformis
Rhazya stricta
Cynanchum wilfordii
Asclepias syriaca
Cuscuta gronovii
Cuscuta australis
Cuscuta campestris
Ipomoea nil
Capsicum annuum
Solanum tuberosum
Physochlaina orientalis
Nicotiana tabacum
Hyoscyamus niger
Sesuvium portulacastrum
Silene vulgaris
Beta vulgaris
Chenopodium quinoa
Spinacia oleracea
Nepenthes ventricosa
Fallopia multiflora
Lagerstroemia indica
Eucalyptus grandis
Xanthoceras sorbifolium
Corchorus olitorius
Hibiscus cannabinus
Bombax ceiba
Gossypium trilobum
Capsella rubella
Arabidopsis thaliana
Schrenkiella parvula
Arabis alpina
Eruca vesicaria
Brassica rapa
Raphanus sativus
Sinapis arvensis
Turritis glabra
Boechera stricta
Batis maritima
Carica papaya
Lophophytum mirabile
Peganum harmala
Citrus sinensis
Morus notabilis
Ziziphus jujuba
Epirixanthes elongata
Malus domestica
Eriobotrya japonica
Prunus avium
Medicago truncatula
Vicia faba
Bupleurum falcatum
Lotus japonicus
Millettia pinnata
Vigna radiata
Glycine soja
Phaseolus vulgaris
Sophora flavescens
Castanospermum australe
Styphnolobium japonicum
Ammopiptanthus nanus
Tamarindus indica
Cercis canadensis
Senna tora
Libidibia coriaria
Haematoxylum brasiletto
Leucaena trichandra
Acacia ligulata
0.02

**nad9**

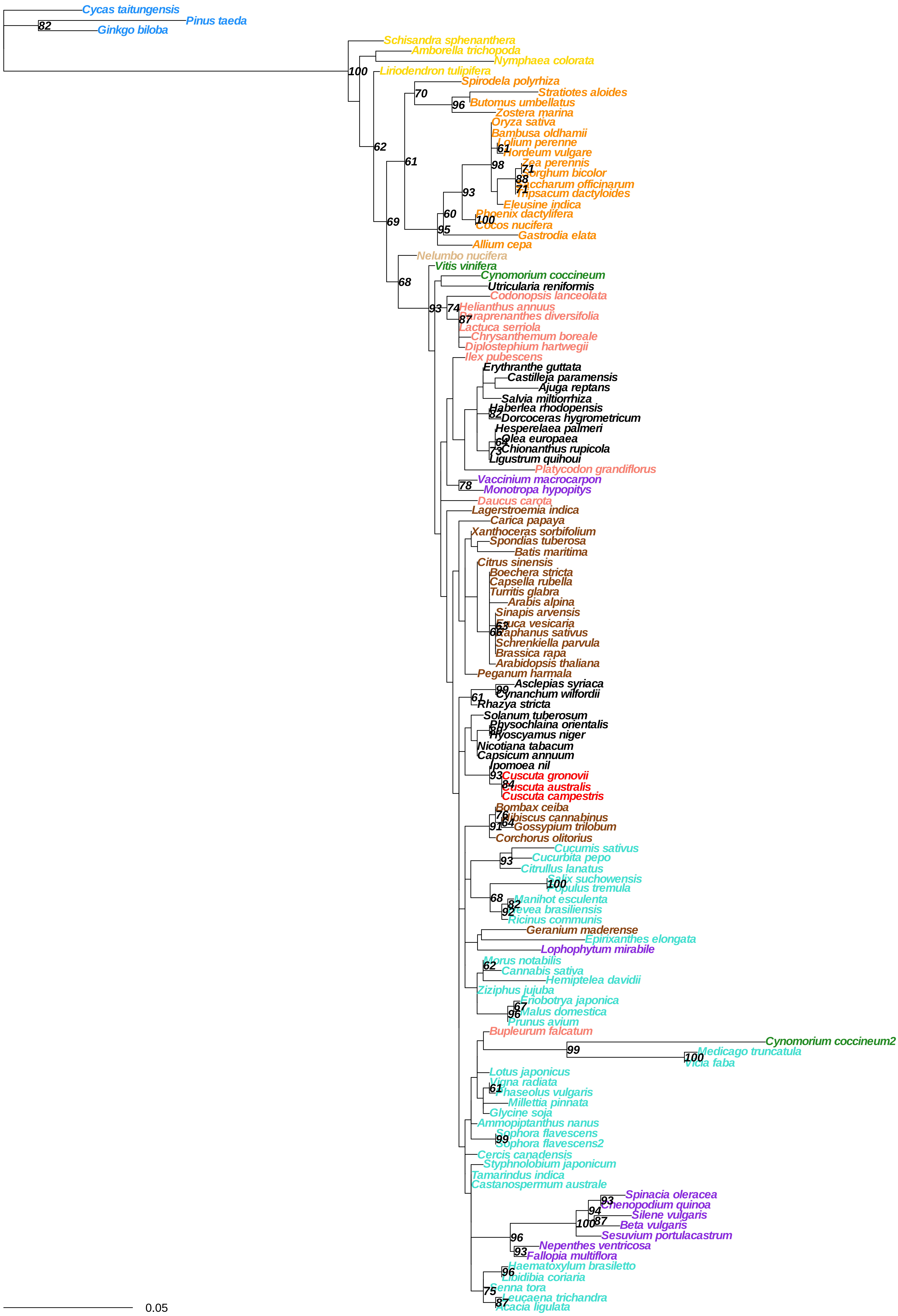

Cycas taitungensis
Pinus taeda
Ginkgo biloba
Schisandra sphenanthera
Amborella trichopoda
Nymphaea colorata
Liriodendron tulipifera
Spirodela polyrhiza
Stratiotes aloides
Butomus umbellatus
Zostera marina
Oryza sativa
Bambusa oldhamii
Lolium perenne
Hordeum vulgare
Zea perennis
Sorghum bicolor
Saccharum officinarum
Tripsacum dactyloides
Eleusine indica
Phoenix dactylifera
Cocos nucifera
Gastrodia elata
Allium cepa
Nelumbo nucifera
Vitis vinifera
Cynomorium coccineum
Utricularia reniformis
Codonopsis lanceolata
Helianthus annuus
Paraprenanthes diversifolia
Lactuca serriola
Chrysanthemum boreale
Diplostephium hartwegii
Ilex pubescens
Erythranthe guttata
Castilleja paramensis
Ajuga reptans
Salvia miltiorrhiza
Haberlea rhodopensis
Dorcoceras hygrometricum
Hesperelaea palmeri
Olea europaea
Chionanthus rupicola
Ligustrum quihoui
Platycodon grandiflorus
Vaccinium macrocarpon
Monotropa hypopitys
Daucus carota
Lagerstroemia indica
Carica papaya
Xanthoceras sorbifolium
Spondias tuberosa
Batis maritima
Citrus sinensis
Boechera stricta
Capsella rubella
Turritis glabra
Arabis alpina
Sinapis arvensis
Eruca vesicaria
Raphanus sativus
Schrenkiella parvula
Brassica rapa
Arabidopsis thaliana
Peganum harmala
Asclepias syriaca
Cynanchum wilfordii
Rhazya stricta
Solanum tuberosum
Physochlaina orientalis
Hyoscyamus niger
Nicotiana tabacum
Capsicum annuum
Ipomoea nil
Cuscuta gronovii
Cuscuta australis
Cuscuta campestris
Bombax ceiba
Hibiscus cannabinus
Gossypium trilobum
Corchorus olitorius
Cucumis sativus
Cucurbita pepo
Citrullus lanatus
Salix suchowensis
Populus tremula
Manihot esculenta
Hevea brasiliensis
Ricinus communis
Geranium maderense
Epirixanthes elongata
Lophophytum mirabile
Morus notabilis
Cannabis sativa
Hemiptelea davidii
Ziziphus jujuba
Eriobotrya japonica
Malus domestica
Prunus avium
Bupleurum falcatum
Cynomorium coccineum2
Medicago truncatula
Vicia faba
Lotus japonicus
Vigna radiata
Phaseolus vulgaris
Millettia pinnata
Glycine soja
Ammopiptanthus nanus
Sophora flavescens
Sophora flavescens2
Cercis canadensis
Styphnolobium japonicum
Tamarindus indica
Castanospermum australe
Spinacia oleracea
Chenopodium quinoa
Silene vulgaris
Beta vulgaris
Sesuvium portulacastrum
Nepenthes ventricosa
Fallopia multiflora
Haematoxylum brasiletto
Libidibia coriaria
Senna tora
Leucaena trichandra
Acacia ligulata
0.05

### rpl16

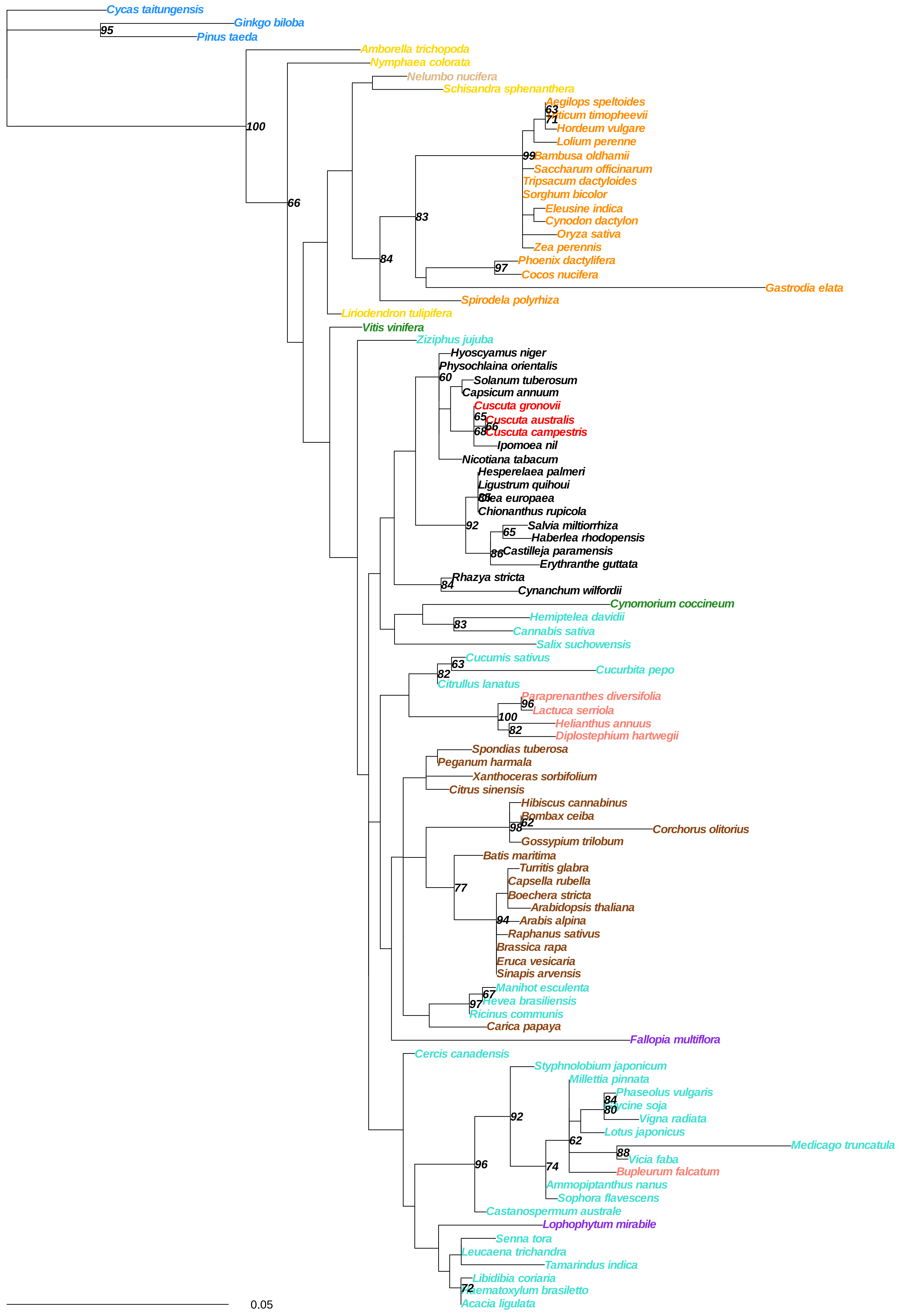

Cycas taitungensis
Ginkgo biloba
95
Pinus taeda
Amborella trichopoda
Nymphaea colorata
Nelumbo nucifera
Schisandra sphenanthera
Aegilops speltoides
63
ticum timopheevii
71
Hordeum vulgare
Lolium perenne
99
Bambusa oldhamii
Saccharum officinarum
Tripsacum dactyloides
Sorghum bicolor
Eleusine indica
Cynodon dactylon
Oryza sativa
Zea perennis
Phoenix dactylifera
97
Cocos nucifera
Gastrodia elata
Spirodela polyrhiza
Liriodendron tulipifera
100
66
83
84
Vitis vinifera
Ziziphus jujuba
Hyoscyamus niger
Physochlaina orientalis
60
Solanum tuberosum
Capsicum annuum
Cuscuta gronovii
65
Cuscuta australis
66
68
Cuscuta campestris
Ipomoea nil
Nicotiana tabacum
Hesperelaea palmeri
Ligustrum quihoui
Chionanthus rupicola
92
Salvia miltiorrhiza
65
Haberlea rhodopensis
86
Castilleja paramensis
Erythranthe guttata
Rhazya stricta
84
Cynanchum wilfordii
Cynomorium coccineum
Hemiptelea davidii
83
Cannabis sativa
Salix suchowensis
Cucumis sativus
63
82
Cucurbita pepo
Citrullus lanatus
Paraprenanthes diversifolia
96
Lactuca serriola
100
Helianthus annuus
82
Diplostephium hartwegii
Spondias tuberosa
Peganum harmala
Xanthoceras sorbifolium
Citrus sinensis
Hibiscus cannabinus
Bombax ceiba
62
98
Corchorus olitorius
Gossypium trilobum
Batis maritima
77
Turritis glabra
Capsella rubella
Boechera stricta
Arabidopsis thaliana
94
Arabis alpina
Raphanus sativus
Brassica rapa
Eruca vesicaria
Sinapis arvensis
Manihot esculenta
67
97
Hevea brasiliensis
Ricinus communis
Carica papaya
Fallopia multiflora
Cercis canadensis
Styphnolobium japonicum
Millettia pinnata
84
Phaseolus vulgaris
80
ycine soja
Vigna radiata
92
Lotus japonicus
62
Medicago truncatula
88
Vicia faba
96
74
Bupleurum falcatum
Ammopiptanthus nanus
Sophora flavescens
Castanospermum australe
Lophophytum mirabile
Senna tora
Leucaena trichandra
Tamarindus indica
Libidibia coriaria
72
aematoxylum brasiletto
Acacia ligulata
0.05

**rpl5**

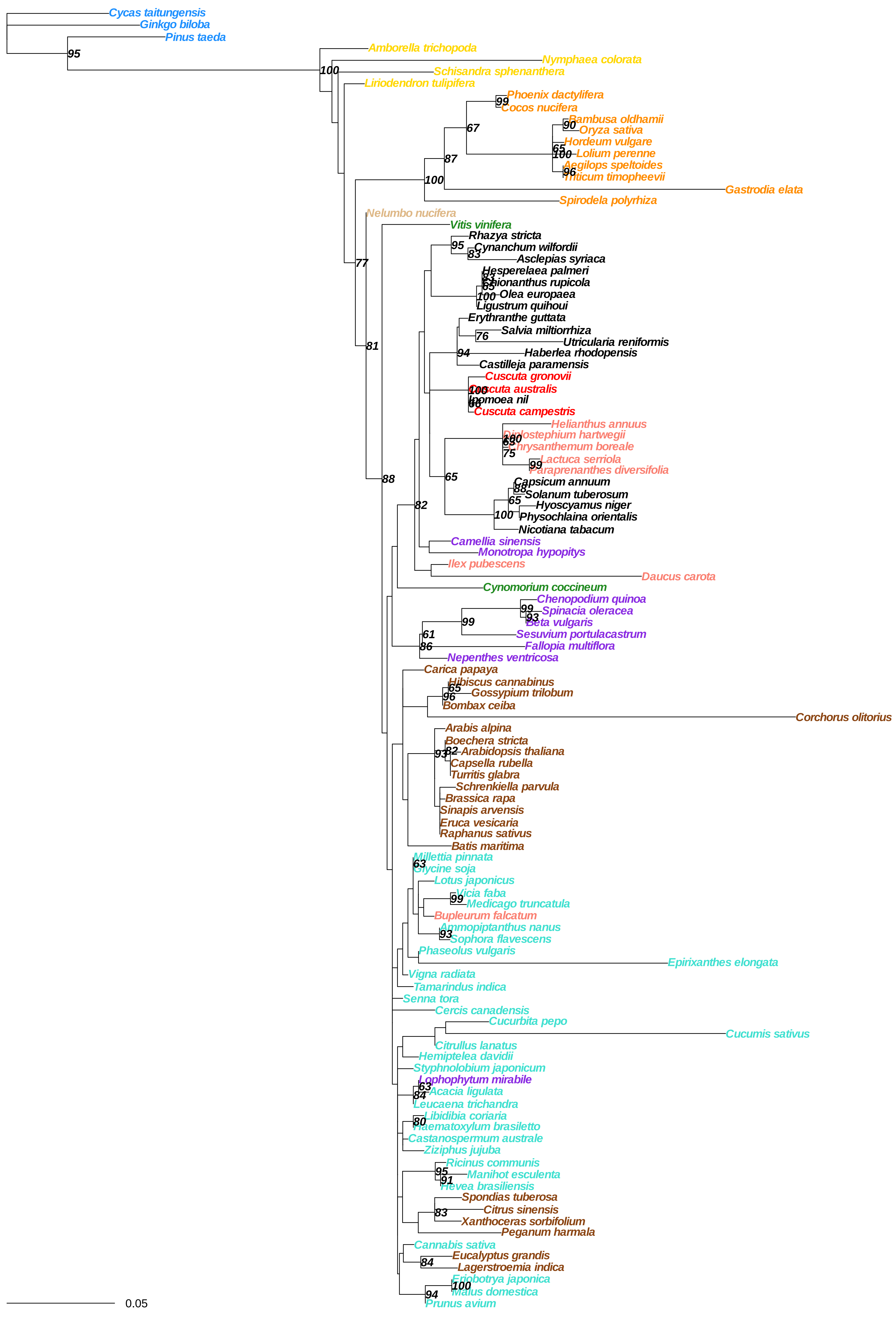
Cycas taitungensis
Ginkgo biloba
Pinus taeda
Amborella trichopoda
Nymphaea colorata
Schisandra sphenanthera
Liriodendron tulipifera
Phoenix dactylifera
Cocos nucifera
Bambusa oldhamii
Oryza sativa
Hordeum vulgare
Lolium perenne
Aegilops speltoides
Triticum timopheevii
Gastrodia elata
Spirodela polyrhiza
Nelumbo nucifera
Vitis vinifera
Rhazya stricta
Cynanchum wilfordii
Asclepias syriaca
Hesperelaea palmeri
Chionanthus rupicola
Olea europaea
Ligustrum quihoui
Erythranthe guttata
Salvia miltiorrhiza
Utricularia reniformis
Haberlea rhodopensis
Castilleja paramensis
Cuscuta gronovii
Cuscuta australis
Ipomoea nil
Cuscuta campestris
Helianthus annuus
Diplostephium hartwegii
Chrysanthemum boreale
Lactuca serriola
Paraprenanthes diversifolia
Capsicum annuum
Solanum tuberosum
Hyoscyamus niger
Physochlaina orientalis
Nicotiana tabacum
Camellia sinensis
Monotropa hypopitys
Ilex pubescens
Daucus carota
Cynomorium coccineum
Chenopodium quinoa
Spinacia oleracea
Beta vulgaris
Sesuvium portulacastrum
Fallopia multiflora
Nepenthes ventricosa
Carica papaya
Hibiscus cannabinus
Gossypium trilobum
Bombax ceiba
Corchorus olitorius
Arabis alpina
Boechera stricta
Arabidopsis thaliana
Capsella rubella
Turritis glabra
Schrenkiella parvula
Brassica rapa
Sinapis arvensis
Eruca vesicaria
Raphanus sativus
Batis maritima
Millettia pinnata
Glycine soja
Lotus japonicus
Vicia faba
Medicago truncatula
Bupleurum falcatum
Ammopiptanthus nanus
Sophora flavescens
Phaseolus vulgaris
Epirixanthes elongata
Vigna radiata
Tamarindus indica
Senna tora
Cercis canadensis
Cucurbita pepo
Cucumis sativus
Citrullus lanatus
Hemiptelea davidii
Styphnolobium japonicum
Lophophytum mirabile
Acacia ligulata
Leucaena trichandra
Libidibia coriaria
Haematoxylum brasiletto
Castanospermum australe
Ziziphus jujuba
Ricinus communis
Manihot esculenta
Hevea brasiliensis
Spondias tuberosa
Citrus sinensis
Xanthoceras sorbifolium
Peganum harmala
Cannabis sativa
Eucalyptus grandis
Lagerstroemia indica
Eriobotrya japonica
Malus domestica
Prunus avium
0.05

**rps1**

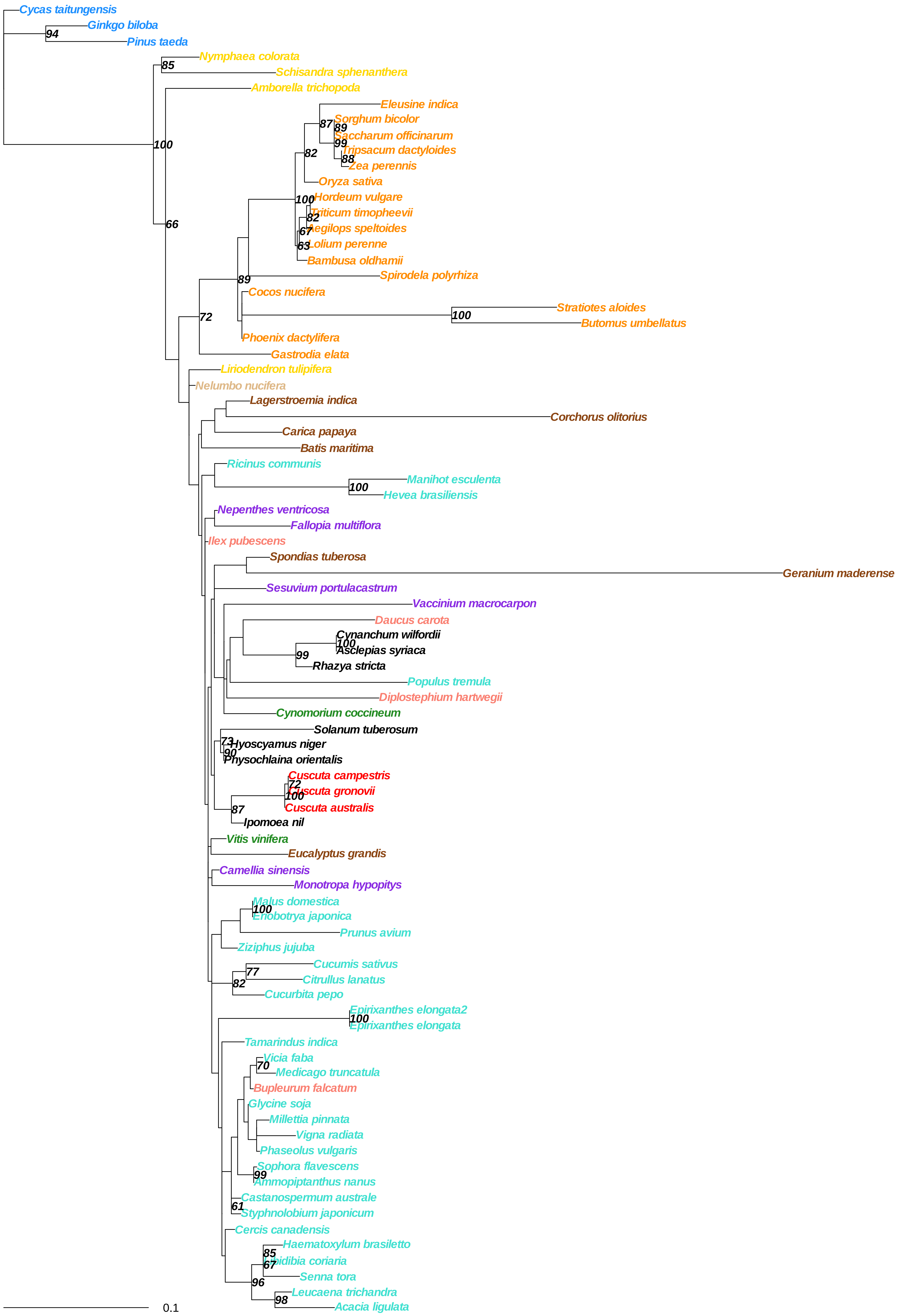
Cycas taitungensis
Ginkgo biloba
94
Pinus taeda
Nymphaea colorata
85
Schisandra sphenanthera
Amborella trichopoda
100
Eleusine indica
87
Sorghum bicolor
89
Saccharum officinarum
99
Tripsacum dactyloides
88
Zea perennis
82
Oryza sativa
100
Hordeum vulgare
82
Triticum timopheevii
67
Aegilops speltoides
63
Lolium perenne
Bambusa oldhamii
66
Spirodela polyrhiza
89
Cocos nucifera
Stratiotes aloides
100
Butomus umbellatus
72
Phoenix dactylifera
Gastrodia elata
Liriodendron tulipifera
Nelumbo nucifera
Lagerstroemia indica
Corchorus olitorius
Carica papaya
Batis maritima
Ricinus communis
Manihot esculenta
100
Hevea brasiliensis
Nepenthes ventricosa
Fallopia multiflora
Ilex pubescens
Spondias tuberosa
Geranium maderense
Sesuvium portulacastrum
Vaccinium macrocarpon
Daucus carota
Cynanchum wilfordii
100
Asclepias syriaca
99
Rhazya stricta
Populus tremula
Diplostephium hartwegii
Cynomorium coccineum
Solanum tuberosum
73
Hyoscyamus niger
90
Physochlaina orientalis
Cuscuta campestris
72
Cuscuta gronovii
100
Cuscuta australis
87
Ipomoea nil
Vitis vinifera
Eucalyptus grandis
Camellia sinensis
Monotropa hypopitys
Malus domestica
100
Eriobotrya japonica
Prunus avium
Ziziphus jujuba
Cucumis sativus
77
Citrullus lanatus
82
Cucurbita pepo
Epirixanthes elongata2
100
Epirixanthes elongata
Tamarindus indica
Vicia faba
70
Medicago truncatula
Bupleurum falcatum
Glycine soja
Millettia pinnata
Vigna radiata
Phaseolus vulgaris
Sophora flavescens
99
Ammopiptanthus nanus
Castanospermum australe
61
Styphnolobium japonicum
Cercis canadensis
Haematoxylum brasiletto
85
Libidibia coriaria
67
Senna tora
96
Leucaena trichandra
98
Acacia ligulata
0.1

### rps12

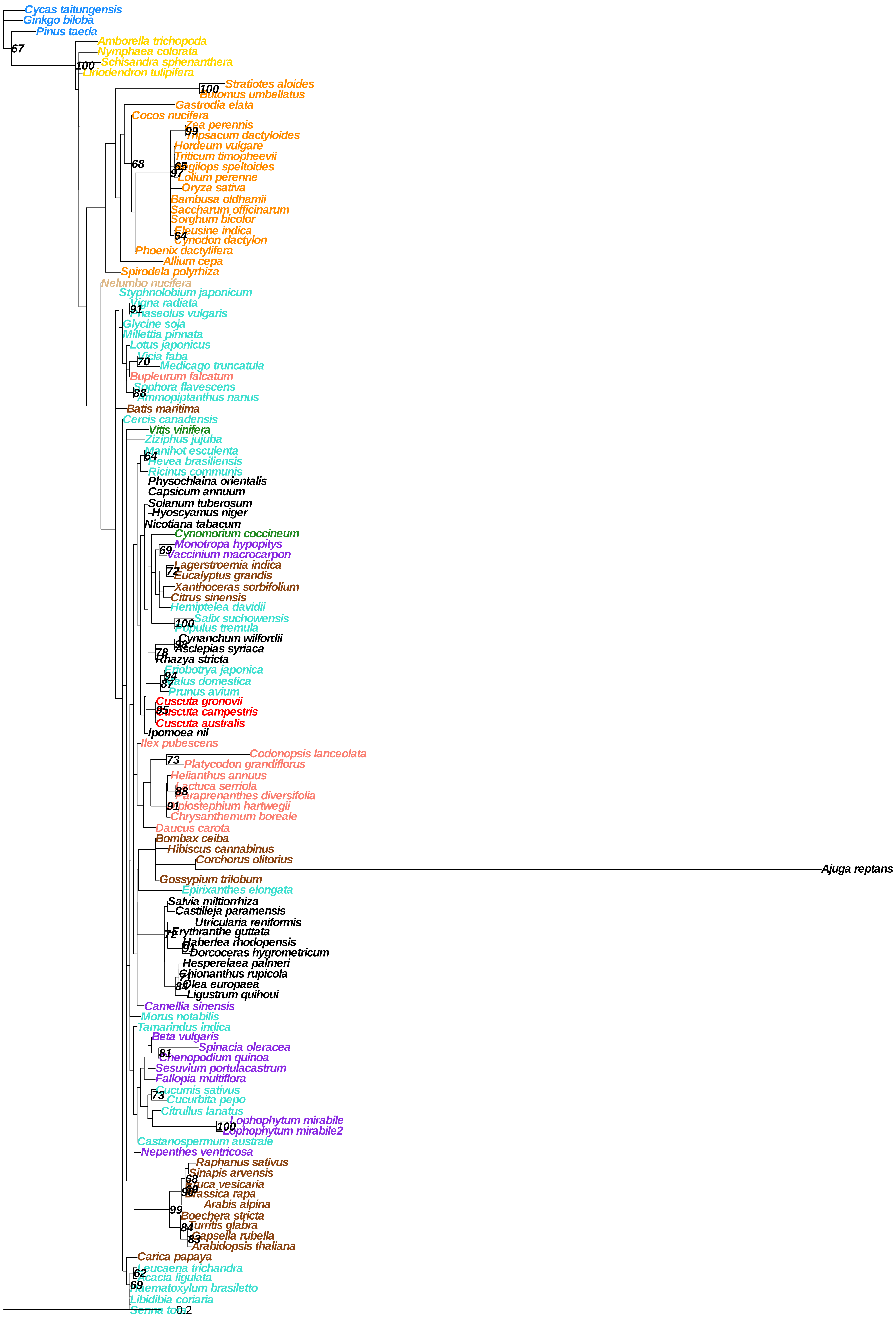

Cycas taitungensis
Ginkgo biloba
Pinus taeda
67
Amborella trichopoda
Nymphaea colorata
100
Schisandra sphenanthera
Liriodendron tulipifera
100
Stratiotes aloides
Butomus umbellatus
Gastrodia elata
Cocos nucifera
Zea perennis
99
Tripsacum dactyloides
Hordeum vulgare
Triticum timopheevii
65
Aegilops speltoides
97
Lolium perenne
68
Oryza sativa
Bambusa oldhamii
Saccharum officinarum
Sorghum bicolor
Eleusine indica
64
Cynodon dactylon
Phoenix dactylifera
Allium cepa
Spirodela polyrhiza
Nelumbo nucifera
Styphnolobium japonicum
Vigna radiata
91
Phaseolus vulgaris
Glycine soja
Millettia pinnata
Lotus japonicus
Vicia faba
70
Medicago truncatula
Bupleurum falcatum
Sophora flavescens
88
Ammopiptanthus nanus
Batis maritima
Cercis canadensis
Vitis vinifera
Ziziphus jujuba
Manihot esculenta
64
Hevea brasiliensis
Ricinus communis
Physochlaina orientalis
Capsicum annuum
Solanum tuberosum
Hyoscyamus niger
Nicotiana tabacum
Cynomorium coccineum
Monotropa hypopitys
69
Vaccinium macrocarpon
Lagerstroemia indica
72
Eucalyptus grandis
Xanthoceras sorbifolium
Citrus sinensis
Hemiptelea davidii
Salix suchowensis
100
Populus tremula
Cynanchum wilfordii
98
Asclepias syriaca
78
Rhazya stricta
Eriobotrya japonica
94
87
Malus domestica
Prunus avium
Cuscuta gronovii
95
Cuscuta campestris
Cuscuta australis
Ipomoea nil
Ilex pubescens
Codonopsis lanceolata
73
Platycodon grandiflorus
Helianthus annuus
Lactuca serriola
88
Paraprenanthes diversifolia
91
Atractylodes hartwegii
Chrysanthemum boreale
Daucus carota
Bombax ceiba
Hibiscus cannabinus
Corchorus olitorius
Ajuga reptans
Gossypium trilobum
Epirixanthes elongata
Salvia miltiorrhiza
Castilleja paramensis
Utricularia reniformis
72
Erythranthe guttata
Haberlea rhodopensis
91
Dorcoceras hygrometricum
Hesperelaea palmeri
Chionanthus rupicola
84
Olea europaea
Ligustrum quihoui
Camellia sinensis
Morus notabilis
Tamarindus indica
Beta vulgaris
Spinacia oleracea
81
Chenopodium quinoa
Sesuvium portulacastrum
Fallopia multiflora
Cucumis sativus
73
Cucurbita pepo
Citrullus lanatus
Lophophytum mirabile
100
Lophophytum mirabile2
Castanospermum australe
Nepenthes ventricosa
Raphanus sativus
Sinapis arvensis
68
Eruca vesicaria
99
98
Brassica rapa
Arabis alpina
99
Boechera stricta
84
Turritis glabra
83
Capsella rubella
Arabidopsis thaliana
Carica papaya
Leucaena trichandra
62
Acacia ligulata
69
Haematoxylum brasiletto
Libidibia coriaria
Senna tora
0.2

rps13

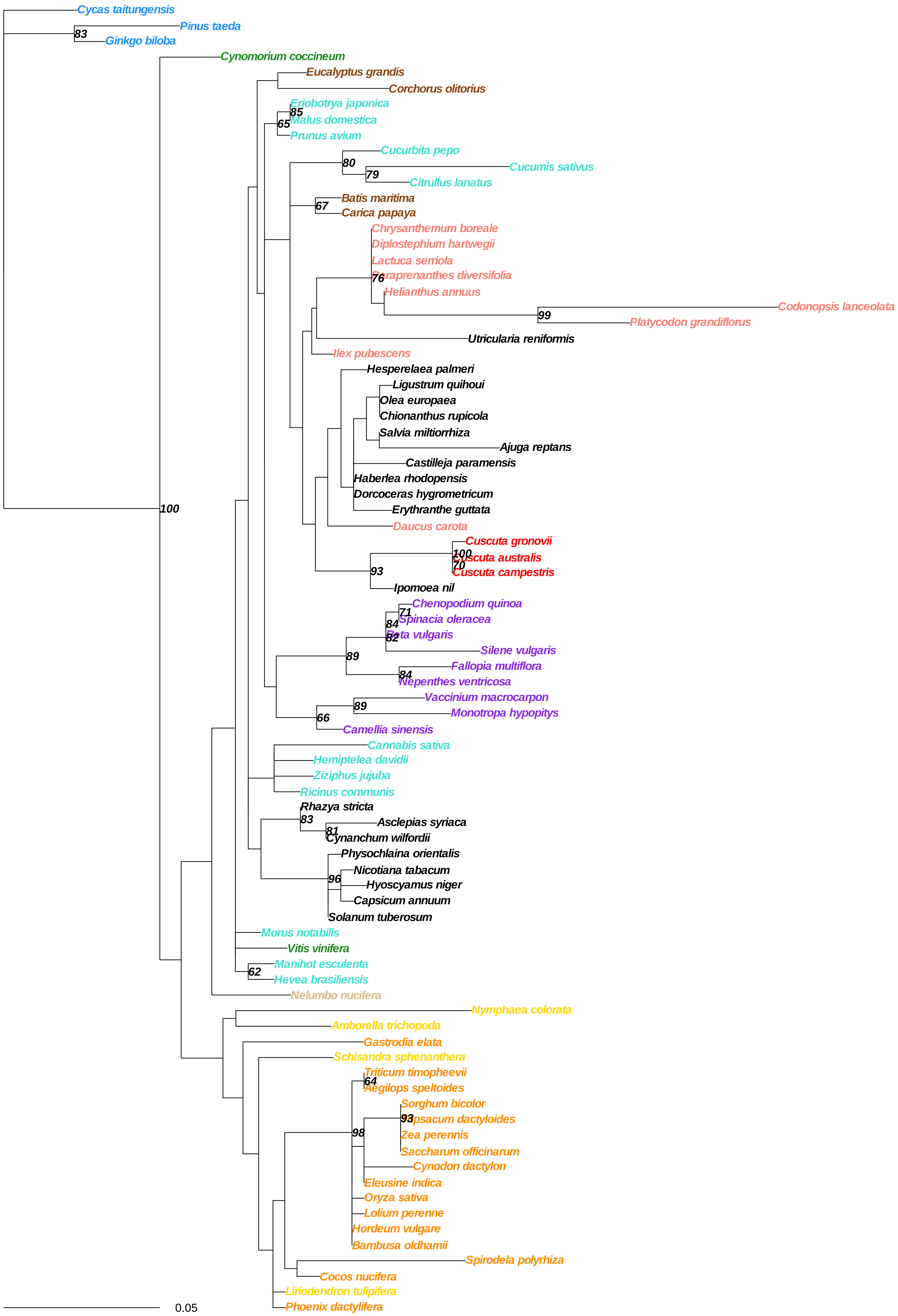

Cycas taitungensis
Pinus taeda
83
Ginkgo biloba
Cynomorium coccineum
Eucalyptus grandis
Corchorus olitorius
Eriobotrya japonica
85
65
Malus domestica
Prunus avium
Cucurbita pepo
80
Cucumis sativus
79
Citrullus lanatus
Batis maritima
67
Carica papaya
Chrysanthemum boreale
Diplostephium hartwegii
Lactuca serriola
Paraprenanthes diversifolia
76
Helianthus annuus
Codonopsis lanceolata
99
Platycodon grandiflorus
Utricularia reniformis
Ilex pubescens
Hesperelaea palmeri
Ligustrum quihoui
Olea europaea
Chionanthus rupicola
Salvia miltiorrhiza
Ajuga reptans
Castilleja paramensis
Haberlea rhodopensis
Dorcoceras hygrometricum
Erythranthe guttata
100
Daucus carota
Cuscuta gronovii
100
Cuscuta australis
70
Cuscuta campestris
93
Ipomoea nil
Chenopodium quinoa
71
84
Spinacia oleracea
82
Beta vulgaris
Silene vulgaris
89
Fallopia multiflora
84
Nepenthes ventricosa
Vaccinium macrocarpon
89
66
Monotropa hypopitys
Camellia sinensis
Cannabis sativa
Hemiptelea davidii
Ziziphus jujuba
Ricinus communis
Rhazya stricta
83
Asclepias syriaca
81
Cynanchum wilfordii
Physochlaina orientalis
Nicotiana tabacum
96
Hyoscyamus niger
Capsicum annuum
Solanum tuberosum
Morus notabilis
Vitis vinifera
Manihot esculenta
62
Hevea brasiliensis
Nelumbo nucifera
Nymphaea colorata
Amborella trichopoda
Gastrodia elata
Schisandra sphenanthera
Triticum timopheevii
64
Aegilops speltoides
Sorghum bicolor
93
Tripsacum dactyloides
Zea perennis
Saccharum officinarum
98
Cynodon dactylon
Eleusine indica
Oryza sativa
Lolium perenne
Hordeum vulgare
Bambusa oldhamii
Spirodela polyrhiza
Cocos nucifera
Liriodendron tulipifera
Phoenix dactylifera
0.05

**rps3**

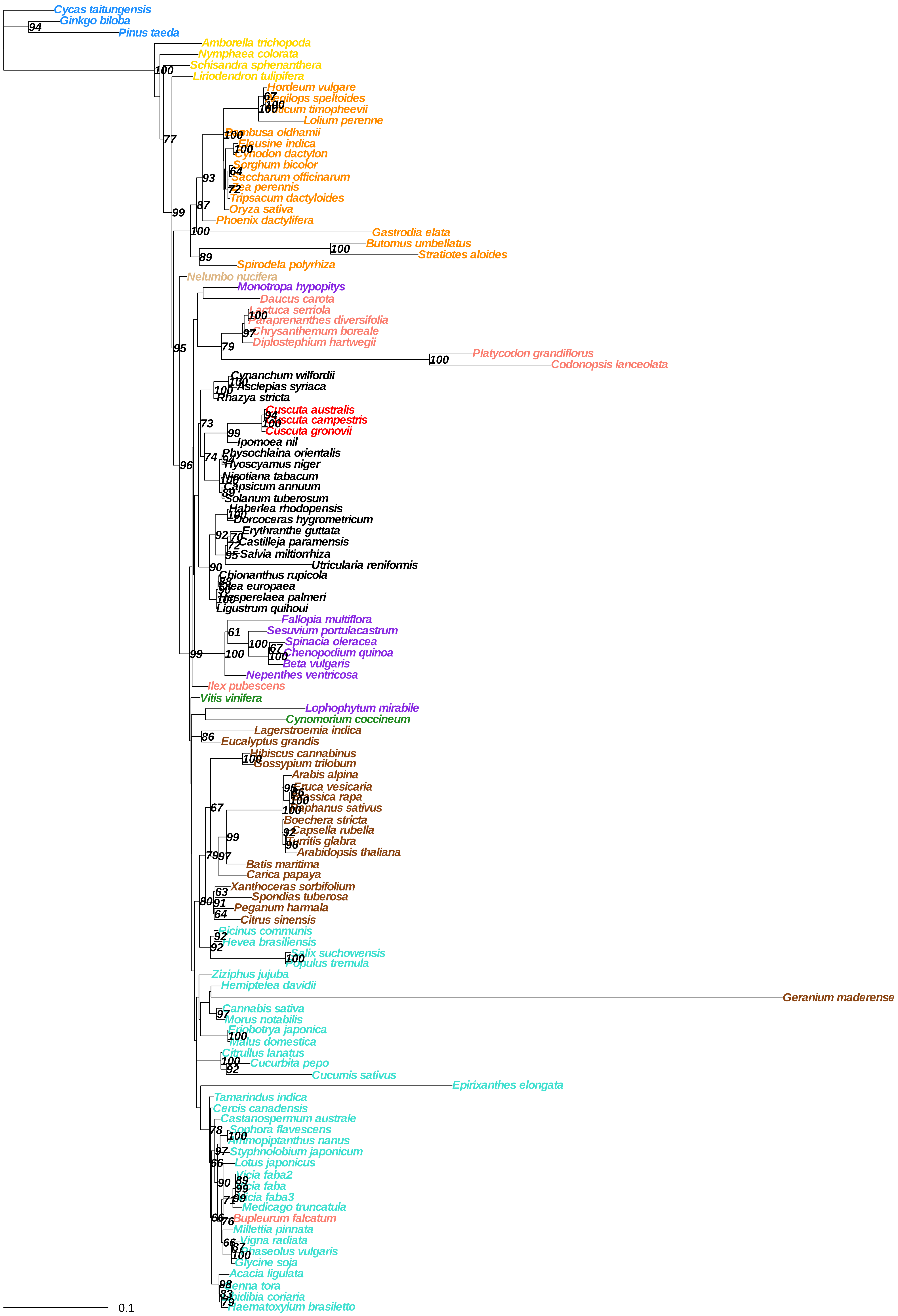

Cycas taitungensis
Ginkgo biloba
Pinus taeda
Amborella trichopoda
Nymphaea colorata
Schisandra sphenanthera
Liriodendron tulipifera
Hordeum vulgare
Aegilops speltoides
Triticum timopheevii
Lolium perenne
Bambusa oldhamii
Eleusine indica
Cynodon dactylon
Sorghum bicolor
Saccharum officinarum
Zea perennis
Tripsacum dactyloides
Oryza sativa
Phoenix dactylifera
Gastrodia elata
Butomus umbellatus
Stratiotes aloides
Spirodela polyrhiza
Nelumbo nucifera
Monotropa hypopitys
Daucus carota
Lactuca serriola
Paraprenanthes diversifolia
Chrysanthemum boreale
Diplostephium hartwegii
Platycodon grandiflorus
Codonopsis lanceolata
Cynanchum wilfordii
Asclepias syriaca
Rhazya stricta
Cuscuta australis
Cuscuta campestris
Cuscuta gronovii
Ipomoea nil
Physochlaina orientalis
Hyoscyamus niger
Nicotiana tabacum
Capsicum annuum
Solanum tuberosum
Haberlea rhodopensis
Dorcoceras hygrometricum
Erythranthe guttata
Castilleja paramensis
Salvia miltiorrhiza
Utricularia reniformis
Chionanthus rupicola
Olea europaea
Hesperelaea palmeri
Ligustrum quihoui
Fallopia multiflora
Sesuvium portulacastrum
Spinacia oleracea
Chenopodium quinoa
Beta vulgaris
Nepenthes ventricosa
Ilex pubescens
Vitis vinifera
Lophophytum mirabile
Cynomorium coccineum
Lagerstroemia indica
Eucalyptus grandis
Hibiscus cannabinus
Gossypium trilobum
Arabis alpina
Eruca vesicaria
Brassica rapa
Raphanus sativus
Boechera stricta
Capsella rubella
Turritis glabra
Arabidopsis thaliana
Batis maritima
Carica papaya
Xanthoceras sorbifolium
Spondias tuberosa
Peganum harmala
Citrus sinensis
Ricinus communis
Hevea brasiliensis
Salix suchowensis
Populus tremula
Ziziphus jujuba
Hemiptelea davidii
Geranium maderense
Cannabis sativa
Morus notabilis
Eriobotrya japonica
Malus domestica
Citrullus lanatus
Cucurbita pepo
Cucumis sativus
Epirixanthes elongata
Tamarindus indica
Cercis canadensis
Castanospermum australe
Sophora flavescens
Ammopiptanthus nanus
Styphnolobium japonicum
Lotus japonicus
Vicia faba2
Vicia faba
Vicia faba3
Medicago truncatula
Bupleurum falcatum
Millettia pinnata
Vigna radiata
Phaseolus vulgaris
Glycine soja
Acacia ligulata
Senna tora
Lupidibia coriaria
Haematoxylum brasiletto
0.1

#### rps4

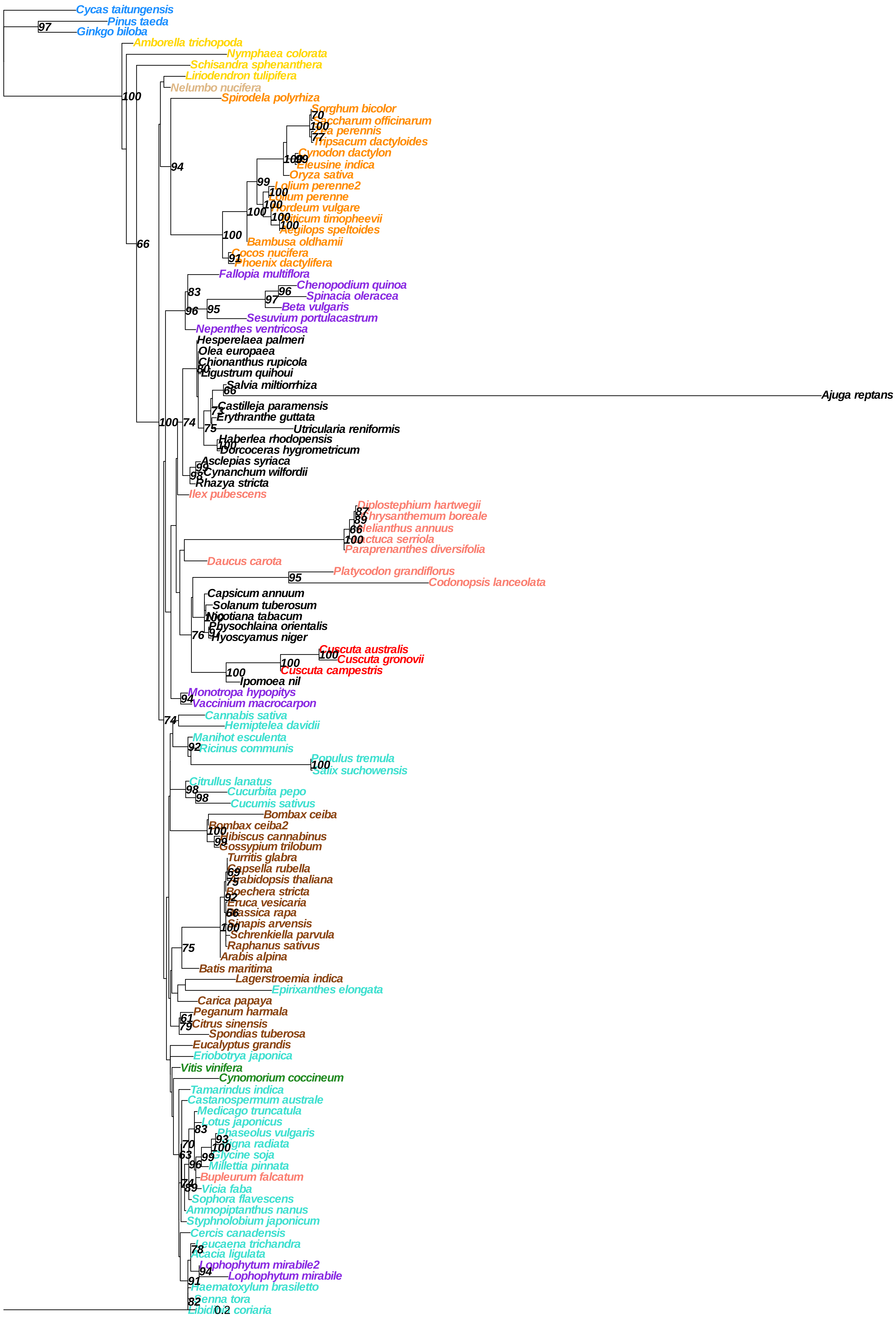
Cycas taitungensis
Pinus taeda
Ginkgo biloba
Amborella trichopoda
Nymphaea colorata
Schisandra sphenanthera
Liriodendron tulipifera
Nelumbo nucifera
Spirodela polyrhiza
Sorghum bicolor
Saccharum officinarum
Zea perennis
Tripsacum dactyloides
Cynodon dactylon
Eleusine indica
Oryza sativa
Lolium perenne2
Lolium perenne
Hordeum vulgare
Triticum timopheevii
Aegilops speltoides
Bambusa oldhamii
Cocos nucifera
Phoenix dactylifera
Fallopia multiflora
Chenopodium quinoa
Spinacia oleracea
Beta vulgaris
Sesuvium portulacastrum
Nepenthes ventricosa
Hesperelaea palmeri
Olea europaea
Chionanthus rupicola
Ligustrum quihoui
Salvia miltiorrhiza
Ajuga reptans
Castilleja paramensis
Erythranthe guttata
Utricularia reniformis
Haberlea rhodopensis
Dorcoceras hygrometricum
Asclepias syriaca
Cynanchum wilfordii
Rhazya stricta
Ilex pubescens
Diplostephium hartwegii
Chrysanthemum boreale
Helianthus annuus
Lactuca serriola
Paraprenanthes diversifolia
Daucus carota
Platycodon grandiflorus
Codonopsis lanceolata
Capsicum annuum
Solanum tuberosum
Nicotiana tabacum
Physochlaina orientalis
Hyoscyamus niger
Cuscuta australis
Cuscuta gronovii
Cuscuta campestris
Ipomoea nil
Monotropa hypopitys
Vaccinium macrocarpon
Cannabis sativa
Hemiptelea davidii
Manihot esculenta
Ricinus communis
Populus tremula
Salix suchowensis
Citrullus lanatus
Cucurbita pepo
Cucumis sativus
Bombax ceiba
Bombax ceiba2
Hibiscus cannabinus
Gossypium trilobum
Turritis glabra
Capsella rubella
Arabidopsis thaliana
Boechera stricta
Eruca vesicaria
Brassica rapa
Sinapis arvensis
Schrenkiella parvula
Raphanus sativus
Arabis alpina
Batis maritima
Lagerstroemia indica
Epirixanthes elongata
Carica papaya
Peganum harmala
Citrus sinensis
Spondias tuberosa
Eucalyptus grandis
Eriobotrya japonica
Vitis vinifera
Cynomorium coccineum
Tamarindus indica
Castanospermum australe
Medicago truncatula
Lotus japonicus
Phaseolus vulgaris
Vigna radiata
Glycine soja
Millettia pinnata
Bupleurum falcatum
Vicia faba
Sophora flavescens
Ammopiptanthus nanus
Styphnolobium japonicum
Cercis canadensis
Leucaena trichandra
Acacia ligulata
Lophophytum mirabile2
Lophophytum mirabile
Haematoxylum brasiletto
Senna tora
Eibidium coriaria
0.2

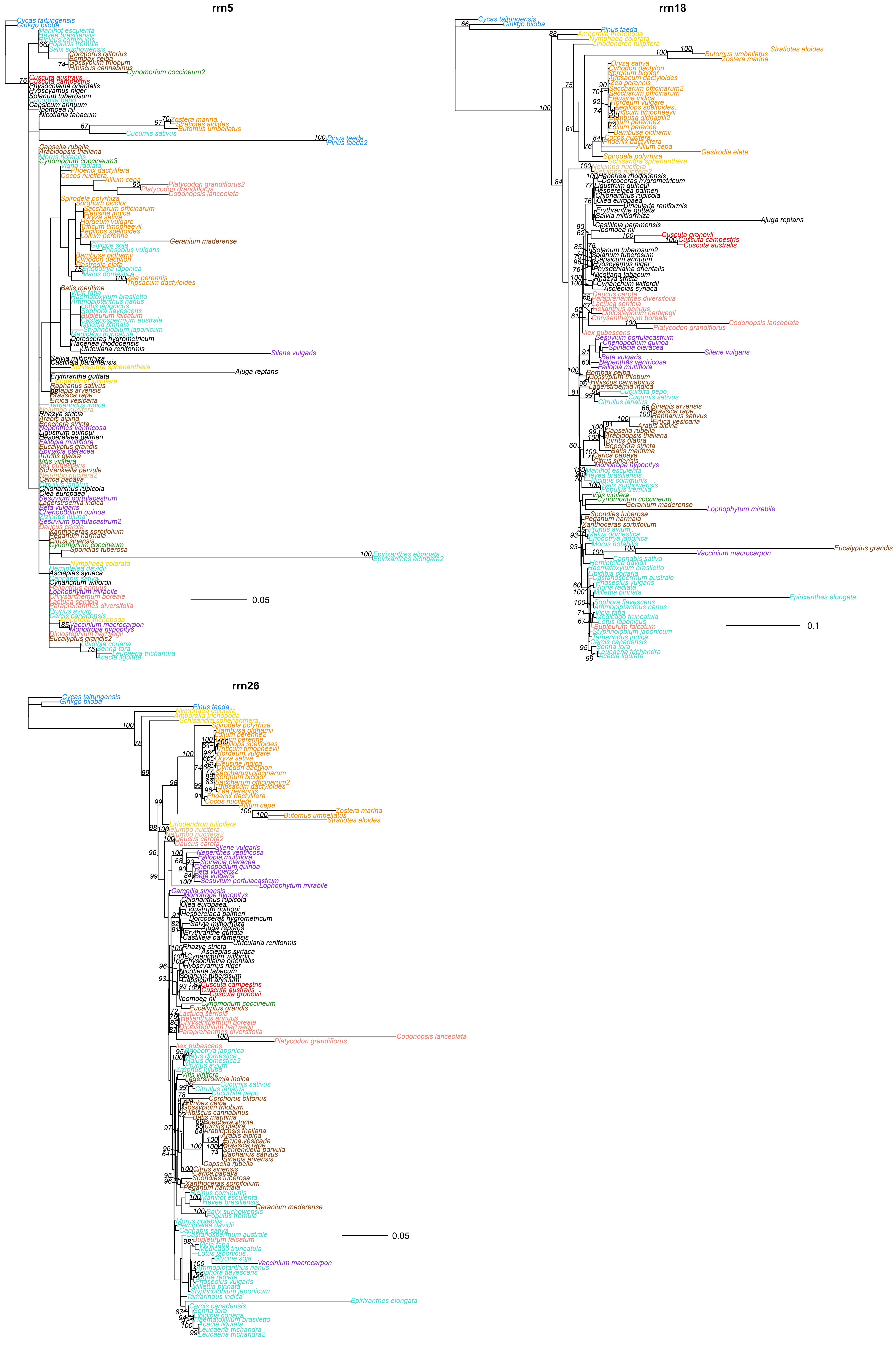

rrn5
rrn18
rrn26
0.05
0.1
0.05
